# Rational engineering of facultative anaerobiosis enables commensal survival in the oxygenated gut

**DOI:** 10.64898/2026.06.05.730451

**Authors:** Abigail E. Rose, Madison Langford-Butler, Luisella Spiga, Daniel W. Bak, Maxwell Neal, Muen Shen, M. Wade Calcutt, Ryan T. Fansler, Sanjay Kumar Rohaun, Mohamad Feron, Alexandra C. Schrimpe-Rutledge, Brittany Berdy, Simona G. Codreanu, Stacy D. Sherrod, Owen F. Hale, Jonathan Livny, John A. McLean, Karsten Zengler, Megan G. Behringer, Eranthie Weerapana, James A. Imlay, Wenhan Zhu

## Abstract

Life originated in the absence of oxygen. Despite its substantial energetic advantages, many modern microbes remain obligate anaerobes, confined to anoxic niches such as the mammalian gut. Why these organisms cannot tolerate oxygen has remained unresolved for more than two centuries. Here, using integrated multi-omics analyses, we identify a network of interlocking vulnerabilities in central metabolism, biosynthetic pathways, and redox homeostasis that together impose an aerobic growth barrier in the obligate anaerobic commensal *Bacteroides thetaiotaomicron*. Rational repair of these vulnerabilities restores metabolic integrity and progressively enhances oxygen tolerance, yielding engineered strains capable of robust growth at 10% O_2_ and markedly improved resilience in the oxygenated, inflamed gut. These findings define a molecular basis for obligate anaerobiosis and establish a framework for engineering commensal bacteria to function in oxidative environments, expanding their ecological range and therapeutic potential.

## Introduction

Life on this planet originated in an environment devoid of oxygen approximately 3.8 billion years ago^1,2^. It was not until about 1.3 billion years later that photosynthesis by cyanobacteria triggered the Great Oxygenation Event, leading to progressive accumulation of oxygen in the atmosphere^1,2^. This profound environmental transition drove the evolution and diversification of organisms capable of exploiting oxygen for metabolic gain^3^. At the same time, however, the rise of oxygen imposed a fundamental challenge for anaerobic life forms whose metabolic architectures were optimized for anoxic conditions and are inherently less compatible with oxygen and its associated redox chemistry^4^. Despite this evolutionary constraint, obligate anaerobes remain widespread in modern environments, occupying anoxic niches such as the mammalian gut^5^.

A central question that has persisted since the initial description of anaerobic organisms by Louis Pasteur nearly two centuries ago is why obligate anaerobes fail to grow when exposed to oxygen^6^. This question extends beyond evolutionary curiosity, as anaerobic bacteria exert important impacts on human health. Under homeostatic conditions, intestinal epithelial cells actively maintain luminal hypoxia by coupling short-chain fatty acid oxidation to mitochondrial oxygen respiration, thereby consuming oxygen diffusing from the underlying vasculature^7–9^. This hypoxic niche supports the dominance of obligate anaerobes, including members of the *Bacteroidetes* phylum^7–9^. However, during episodes of intestinal inflammation, a switch of the colonocyte metabolism from oxidative phosphorylation to anaerobic glycolysis reduces the consumption of oxygen, allowing its diffusion into the otherwise anaerobic gut lumen^10–15^.

In parallel, the immune response generates reactive oxygen species (ROS) as part of its defense mechanism^12,16–18^. Together, these changes impose acute oxidative stress on resident anaerobes and are closely associated with their depletion and the concomitant expansion of facultative anaerobes, driving the microbiota toward a dysbiotic state^13,19^. In contrast to a balanced gut microbial community that provides pivotal functions to human health, a dysbiotic microbiota causes and exacerbates disease conditions including infectious colitis and inflammatory bowel diseases in susceptible hosts^20–23^.

A complementary approach to defining the molecular basis of oxygen sensitivity in obligate anaerobes is to engineer representative commensals to withstand oxygen exposure. A goal-driven, engineering perspective provides a direct experimental framework to identify the metabolic pathways and biochemical processes that limit growth under oxic conditions, thereby revealing the mechanisms that underlie anaerobe-oxygen incompatibility. Enhancing oxygen tolerance also offers a strategy to improve commensal resilience in the inflamed gut, where elevated oxygen and reactive oxygen species disrupt microbial homeostasis^24,25^. Beyond host-associated contexts, increased oxygen tolerance may also facilitate the production and deployment of anaerobic bacteria for a wide range of applications by reducing the need for strictly anaerobic manufacturing and storage conditions^26^. Despite these potential advantages, efforts to engineer oxygen tolerance in obligate anaerobes have been limited, in part due to an incomplete understanding of the molecular determinants of oxygen sensitivity. Seminal studies from the Imlay laboratory and others have revealed that some catalytic mechanisms employed by anaerobic bacteria are intrinsically vulnerable to oxygen and its reactive byproducts^16^. For instance, anaerobes often rely on glycyl radical-based chemistry to metabolize growth substrates such as pyruvate. These radicals readily react with molecular oxygen to form peroxyl radicals, leading to irreversible cleavage of the polypeptide backbone^27^. In addition, anaerobic metabolism relies heavily on low-potential metal centers, including iron-sulfur (Fe-S) clusters in central metabolic enzymes and mononuclear iron centers in hydrogenases^16,28^. These cofactors are notably susceptible to oxygen through oxidative damage and adventitious electron transfer that generates reactive oxygen species (ROS), including superoxide and hydrogen peroxide^28–32^. Such processes can destabilize metal centers and, together with oxidation of redox-sensitive residues such as cysteines, propagate enzyme inactivation and metabolic failure^16^. In parallel, recent work has shown that natural variation in oxygen-response regulators can enhance oxygen tolerance in gut anaerobes^33^, suggesting that adaptation to oxidative stress can occur through both metabolic and regulatory mechanisms.

Despite these advances, a systems-level understanding of how oxygen disrupts anaerobic physiology remains incomplete. In particular, it is unclear which specific enzymatic vulnerabilities are consequential *in vivo* and how they collectively give rise to the aerobic growth barrier. Addressing this gap is essential for defining the molecular basis of obligate anaerobiosis and for developing strategies to manipulate anaerobic microbes in health and disease.

Here, we integrate multi-omics analyses with targeted genetic and biochemical analyses to systematically define oxygen-imposed vulnerabilities in the human gut commensal *Bacteroides thetaiotaomicron* (*B. theta*). We identify a set of interdependent metabolic and redox constraints that limit growth under oxic conditions and show that stepwise repair of these vulnerabilities progressively restores metabolic function and progressively extends oxygen tolerance, enabling robust growth at 10% O_2_. Moreover, these engineered strains exhibit enhanced resilience in the oxygenated, inflamed gut. Together, our findings provide a mechanistic framework for understanding obligate anaerobiosis and establish a strategy for engineering commensal bacteria to function in oxidative environments.

## Results

### Repairing electron carrier alone had a limited impact on cellular oxidative tolerance

A defining feature of anaerobic metabolism is the continuous transfer of electrons between redox-active reactions, a process that maintains intracellular redox balance and supports energy conservation^34^. In many anaerobes, this function is mediated by ferredoxin (Fd), a low-redox-potential electron carrier that uses iron-sulfur ([4Fe-4S] or [2Fe-2S]) clusters to shuttle electrons from reductive processes, such as pyruvate oxidation, to downstream electron sinks including hydrogen evolution^35–37^. While ferredoxins are well suited for anaerobic metabolic pathways, their solvent-exposed metal clusters could be highly susceptible to oxidative damage: interaction with molecular oxygen can inactivate the cofactor and promote the formation of partially reduced oxygen species, including superoxide (O_2_⁻·) and hydrogen peroxide (H_2_O_2_)^38–41^, thereby amplifying oxidative stress^42,43^. In contrast, flavodoxin (FldA) is an iron-free, flavin mononucleotide-based electron carrier that is utilized under iron-limiting conditions as an iron-sparing measure^44^. Notably, flavodoxin expression has been correlated with enhancing resistance to oxidative stress^45^, raising the possibility that replacing ferredoxin with flavodoxin could mitigate oxygen-induced redox damage in anaerobic cells. We therefore hypothesized that substituting this oxygen-labile electron carrier would alleviate a key redox vulnerability imposed by oxygen exposure.

To test this idea, we used *B. theta*, a genetically tractable obligate anaerobe representative of the human microbiota^46^. Consistent with its strict anaerobic lifestyle, *B. theta* growth is highly sensitive to oxygen exposure; growth was completely abolished at 0.5% O_2_ even under sealed conditions that permit gradual oxygen detoxification (**Fig. 1A**). To explore whether oxygen resistance could be rationally engineered into *B. theta*, we replaced the endogenous open reading frame of the ferredoxin with that of the endogenous flavodoxin (BT_0517), generating a “swap” mutant that lacks ferredoxin while flavodoxin is expressed under both anaerobic and aerobic conditions (**Supplementary Fig. S1A**). Exposure of wild-type cells to room air (20.9% O_2_) led to robust generation of H_2_O_2_, while the swap mutant exhibited reduced H_2_O_2_ production (**Fig. 1B**). This modification successfully conferred a competitive advantage over the wild-type strain at low oxygen tensions (0.5% and 1% O_2_; **Fig. 1C**) but not under anaerobic conditions (pO_2_ < 0.0001%), demonstrating an overall improvement in robustness.

**Fig. 1.**
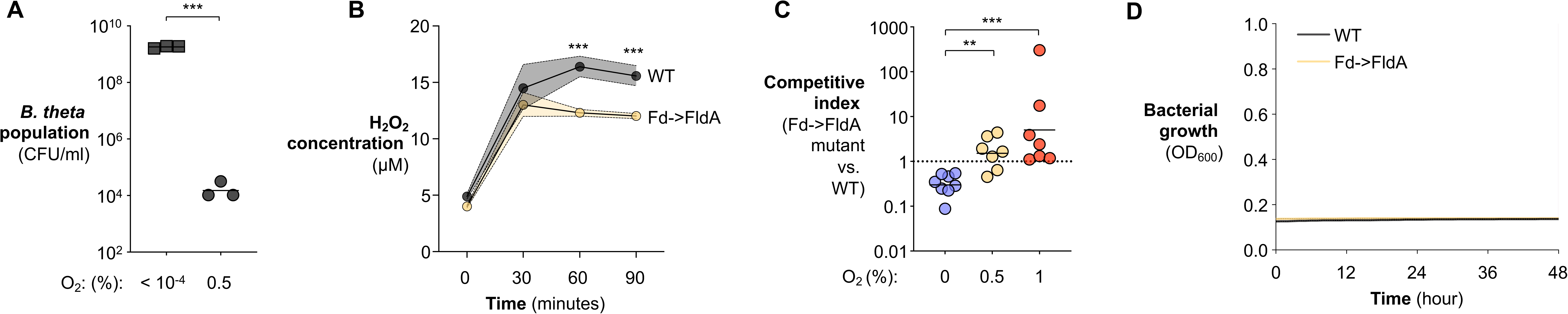
Replacing an Fe-S-containing electron carrier with an iron-independent equivalent confers limited oxygen tolerance in *B. theta*. (**A**) *B. theta* wild-type cells were inoculated into sealed Erlenmeyer flasks containing the indicated oxygen concentrations and incubated for 16 h at 37 °C. Viable bacterial counts were quantified by selective plating. (**B**) Anaerobically growing cultures of the indicated *B. theta* strains were washed and exposed to room air (20.9% O_2_) in glycylglycine-buffered glucose. Accumulation of H_2_O_2_ in the culture medium was monitored over time. (**C**) The ferredoxin-to-flavodoxin swap mutant (Fd→FldA) and wild-type *B. theta* were mixed at a 1:1 ratio and cultured for 16 h in sealed Erlenmeyer flasks containing the indicated oxygen concentrations. Competitive indices were determined by selective plating. (**D**) The indicated *B. theta* strains were cultured in an open atmosphere containing 2% O₂ with constant shaking, and growth was monitored by measuring OD_600_ over time. Bars represent geometric means. *, *P* < 0.05; **, *P* < 0.01.

To stringently assess whether mitigating this redox vulnerability was sufficient to support growth under higher oxygen tensions, we challenged *B. theta* strains at low inoculum density (1×10^5^ CFU) in an open hypoxia system with controlled oxygen concentrations (0.5-10% O_2_). This design minimizes population-level oxygen depletion caused by collective respiration (e.g., through the *B. theta* cytochrome *bd* oxidase) and oxygen scavenging at high bacterial density, both of which can progressively lower oxygen tension in sealed vessels^47–49^. To further reduce heterogeneous oxygen gradients caused by diffusion limitation, we monitored bacterial growth only in the outer wells of 96-well plates under constant shaking^49^. Under these conditions, the swap mutant failed to sustain any appreciable growth at 2% oxygen (**Fig. 1D**), a concentration 100-fold higher than its oxygen tolerance^48^. These results indicate that ferredoxin contributes to oxygen vulnerability in *B. theta*, but that repairing this single redox liability is insufficient to overcome the broader metabolic constraints that prevent aerobic growth in an obligate anaerobe. We therefore used this partial rescue as an entry point to identify additional oxygen-sensitive metabolic bottlenecks.

### Genome-wide identification of oxygen-imposed metabolic constraints

The limited rescue achieved by repairing electron transfer prompted us to pursue a broader, systems-level analysis to identify additional metabolic bottlenecks imposed by oxygen exposure. To achieve this, we profiled the *B. theta* transcriptome under ∼4% oxygen, approximating oxygen levels in the vascularized submucosa and conditions encountered during intestinal inflammation^50,51^. In parallel, we collected cell-free extracellular metabolites for untargeted metabolomic analysis (**Fig. 2A**). Together, these datasets revealed a robust oxidative stress response accompanied by widespread metabolic disruption^16,52^ (**Fig. 2B & Supplementary Fig. S1B-D, transcriptomics; Supplementary Fig. S1E-I, untargeted metabolomics**). Notably, genes encoding core glycolytic enzymes, including hexokinase and fructose-bisphosphate aldolase, were consistently downregulated (**Fig. 2C, D**), suggesting feedback inhibition of glycolysis in response to downstream metabolic constraints.

**Fig. 2.**
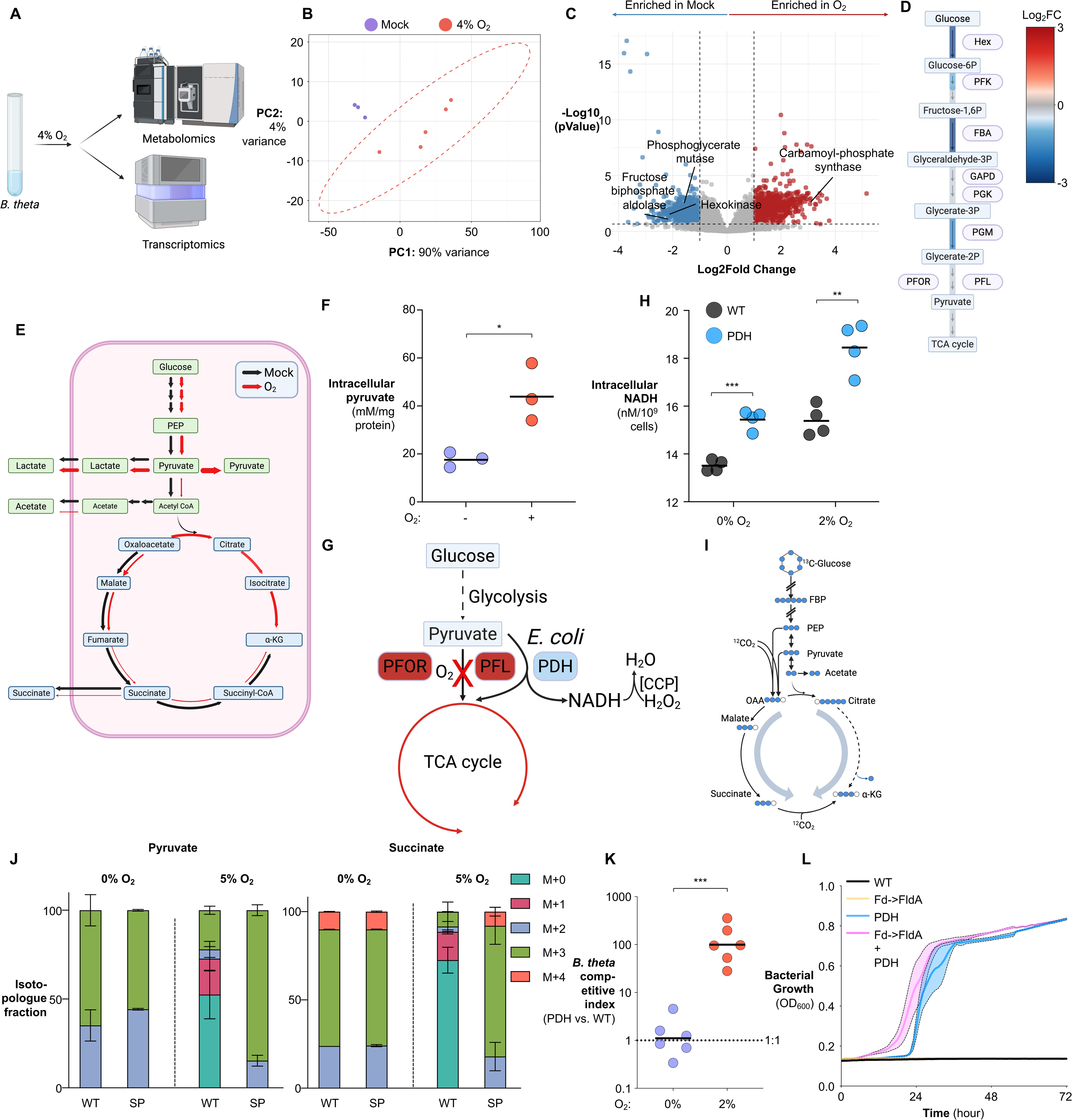
Oxygen impedes central metabolism in *B. theta*. (**A-E**) Anaerobically growing *B. theta* wild-type cells were exposed to 4% O₂ for 30 min at 37 °C, after which the transcriptome was profiled by RNA-seq. Transcriptomic data were integrated into a draft genome-scale metabolic model (GEM) to generate a context-specific metabolic model. (**A**) Experimental schematic. (**B**) Principal component analysis of RNA-seq datasets. Ellipse represents 95% confident interval. (**C**) Volcano plot depicting the transcriptional response of *B. theta* to oxygen exposure. (**D**) Oxygen-induced changes in transcript abundance of glycolytic genes. (**E**) Predicted flux through central metabolic pathways in the context-specific GEM under anaerobic versus oxygen-exposed conditions. (**F**) Anaerobically growing *B. theta* wild-type cells were either maintained under anaerobic conditions or exposed to room air (20.9% O_2_) for 30 min. Intracellular pyruvate levels were quantified by LC-MS. (**G**) Schematic of the engineered *B. theta* strain expressing an oxygen-resistant pyruvate dehydrogenase (PDH) complex. PFOR, pyruvate:ferredoxin oxidoreductase; PFL, pyruvate formate lyase; PDH, pyruvate dehydrogenase; CCP, cytochrome c peroxidase. (**H**) Anaerobically grown cultures of the indicated *B. theta* strains were either maintained anaerobically or exposed to 2% O₂ for 30 min. PDH activity was measured using intracellular NADH level as a proxy. (**I-J**) Anaerobically growing cultures of the indicated *B. theta* strains were exposed to 5% O_2_ for the indicated time and subsequently incubated with [U-^13^C]glucose. Label incorporation into metabolic intermediates was analyzed by high-resolution LC-MS/MS. (**I**) Schematic illustrating label propagation and inferred metabolic fluxes following the glucose isotope shift. Abbreviations: α-KG, α-ketoglutarate; FBP, fructose-1,6-bisphosphate; OAA, oxaloacetate; PEP, phosphoenolpyruvate. (**J**) Fractional abundance of ^13^C-labeled isotopologues following the shift from unlabeled glucose to [U-^13^C]glucose. [M+X] denotes a metabolite isotopologue containing X ^13^C atoms (maximum X equals the number of carbons in the molecule). (**K**) The PDH-expressing strain and wild-type *B. theta* were mixed at a 1:1 ratio and cultured for 16 h in an open atmosphere containing the indicated oxygen concentrations. Competitive indices were determined by selective plating. (**L**) The indicated *B. theta* strains were cultured in an open atmosphere containing 2% O_2_ with constant shaking, and growth was monitored by OD_600_. Bars represent geometric means. Statistical significance is indicated as follows: *. *P* < 0.05; *\*\*, P* < 0.01. \**\*\*, P* < 0.001.

To identify pathways selectively inactivated by oxygen across the genome, we integrated transcriptomic data into a genome-scale metabolic model (GEM)^53^. Using the heuristic thresholding algorithm StanDep^54^ to infer condition-specific sets of active enzymes, we generated context-specific metabolic models via model extraction method mCADRE^55,56^ and computed reaction flux distributions using Markov chain Monte Carlo sampling^57^. Comparing the flux distributions under anaerobic and O_2_-exposed conditions revealed broad impairment of central metabolism under oxygen stress (**Supplementary Fig. S2A-G**). Among the most pronounced defects was a marked restriction of flux from pyruvate to acetyl-CoA (**Fig. 2E**), consistent with the accumulation of phosphoenolpyruvate and pyruvate in oxygen-exposed cells (**Fig. 2F, Supplementary Fig. S2H**).

In anaerobes, conversion of pyruvate to acetyl-CoA is primarily catalyzed by pyruvate formate lyase (PFL)^58^ and pyruvate ferredoxin oxidoreductase (PFOR, **Fig. 2G**) ^59^. Both enzymes are intrinsically oxygen-sensitive: the glycyl radical required for PFL catalysis reacts with oxygen, leading to irreversible polypeptide backbone cleavage^58,60^, while PFOR is rapidly inactivated upon aeration, likely through oxidative damage to its iron-sulfur clusters^29^. These established vulnerabilities provide a mechanistic basis for the oxygen-induced blockade of pyruvate assimilation.

We therefore asked whether bypassing this oxygen-sensitive node could restore central metabolism under oxic conditions. We leveraged the evolutionary solution employed by facultative anaerobes such as *Escherichia coli*, which readily catabolizes pyruvate in room air (20.9% O_2_)^61^. *E. coli* encodes the pyruvate dehydrogenase (PDH) complex to split pyruvate, which, in contrast to PFL and PFOR, is aerotolerant and funnels pyruvate into the TCA cycle while generating reducing power in the form of NADH^62^. We introduced the *E. coli* PDH E1 and E2–E3 components into the *B. theta* genome under control of the native PFL promoter, generating a strain in which PDH expression mirrors that of the endogenous pyruvate-splitting machinery (**Fig. 2G, Supplementary Fig. S2I**). Because PDH catalysis generates NADH stoichiometrically during pyruvate oxidation^62^, intracellular NADH levels provide a readout of PDH-dependent metabolic activity. In contrast to PFOR, which lost activity after exposure to room oxygen^29^, the engrafted PDH retained catalytic activity under similar conditions (**Fig. 2H**). Consistent with its NADH-producing activity, PDH expression maintained intracellular NADH levels even during extreme oxygen challenge (20.9% O_2_) (**Supplementary Fig. S2J**) and increased pools of citrate, α-ketoglutarate, and ATP, indicating partial restoration of central metabolic activity (**Supplementary Fig. S2K-M**).

To directly assess the impact of PDH engraftment on central carbon flux, we performed a [U-^13^C_6_]glucose fingerprint assay^63^. Actively growing cells were exposed to oxygen, washed to remove unlabeled substrate, and supplied with isotopically labeled glucose, enabling high-resolution tracing of central carbon flux by mass spectrometry (**Fig. 2I**). Exposure to 2% O_2_ markedly reduced incorporation of ^13^C into pyruvate and downstream TCA-cycle intermediates, consistent with impaired flux through PFL and PFOR (**Supplementary Fig. S2N**). Expression of PDH partially restored isotopic incorporation into central carbon metabolites under oxygen exposure, demonstrating functional restoration of pyruvate assimilation under otherwise restrictive oxic conditions (**Supplementary Fig. S2N**). We next asked whether combining PDH expression with ferredoxin replacement further improved metabolic flux. The resulting SP strain exhibited enhanced isotopic incorporation into TCA-cycle intermediates even at higher oxygen tension (5% O_2_; **Fig. 2J**). Thus, PDH restores oxygen-impaired pyruvate assimilation, while ferredoxin replacement provides an additional but incomplete improvement in central carbon metabolism.

This metabolic repair translated into improved fitness under hypoxic conditions: the PDH-expressing strain outcompeted the wild type at 2% O_2_ and exhibited robust growth (**Fig. 2K-L**). Combining PDH expression with ferredoxin replacement further enhanced growth (SP strain), albeit modestly, consistent with additive relief of distinct oxygen-sensitive constraints (**Fig. 2L**). Importantly, this benefit was not attributable to reduction of oxidative stress, as PDH-expressing cells exhibited elevated ROS levels during oxygen exposure (**Supplementary Fig. S2O**). This increase likely reflects enhanced electron flow through restored central metabolism, which can amplify ROS production via oxygen interactions with low-potential metal centers or electron leakage from the respiratory chain^31,64^. Together, these results identify pyruvate assimilation as a critical oxygen-sensitive bottleneck in *B. theta* and demonstrate that partial restoration of central metabolic flux is sufficient to substantially enhance anaerobic growth under microaerobic conditions.

### Intracellular ROS impairs anaerobic growth upon oxygen exposure

Although PDH expression restored part of central carbon flux and improved growth at 2% O_2_, increasing oxygen levels still completely arrested growth (**Supplementary Fig. S3A**), indicating that restored pyruvate assimilation exposed additional oxygen-imposed constraints. One prominent candidate is the remarkable propensity of *B. theta* to generate intracellular reactive oxygen species (ROS), including superoxide, upon oxygen exposure^28^. Elevated intracellular ROS potently inactivate vulnerable enzymes, particularly [4Fe-4S] dehydratases and non-redox mononuclear iron enzymes, thereby constricting metabolic capacity even when carbon flux is partially restored (**Supplementary Fig. S3B**)^28,31,64^. Indeed, the cytosolic environment of aerated *B. theta* is so oxidative that even heterologously expressed *E. coli* fumarase, a [4Fe-4S] dehydratase that remains active in aerated *E. coli*, loses activity when expressed in *B. theta*^28^. Together, these observations suggest that oxygen sensitivity in *B. theta* reflects not only discrete pathway bottlenecks but also a cellular redox environment that amplifies ROS-mediated enzyme damage. Despite this pronounced phenotype, the molecular sources of excessive ROS generation in *B. theta* remain unclear, limiting our abilities to mitigate oxidative stress at its origin rather than buffering its downstream consequences.

Because ROS may be produced through incomplete reduction of oxygen by endogenous redox-active enzymes^65^, we hypothesize that limiting electron leakage into oxygen could enhance oxygen tolerance. To test this, we evolved the SP mutant strain under 3% O_2_, a condition that restricts growth yet permits survival, thereby allowing selection for variants with improved oxidative fitness (**Fig. 3A**). Strikingly, sequencing of independently evolved clones revealed a single recurrent mutation in all isolates, mapping to *oxe*, which encodes a predicted flavoprotein^66^. This mutation resulted in a cysteine-to-tyrosine substitution (C222Y; **Fig. 3B**).

**Fig. 3.**
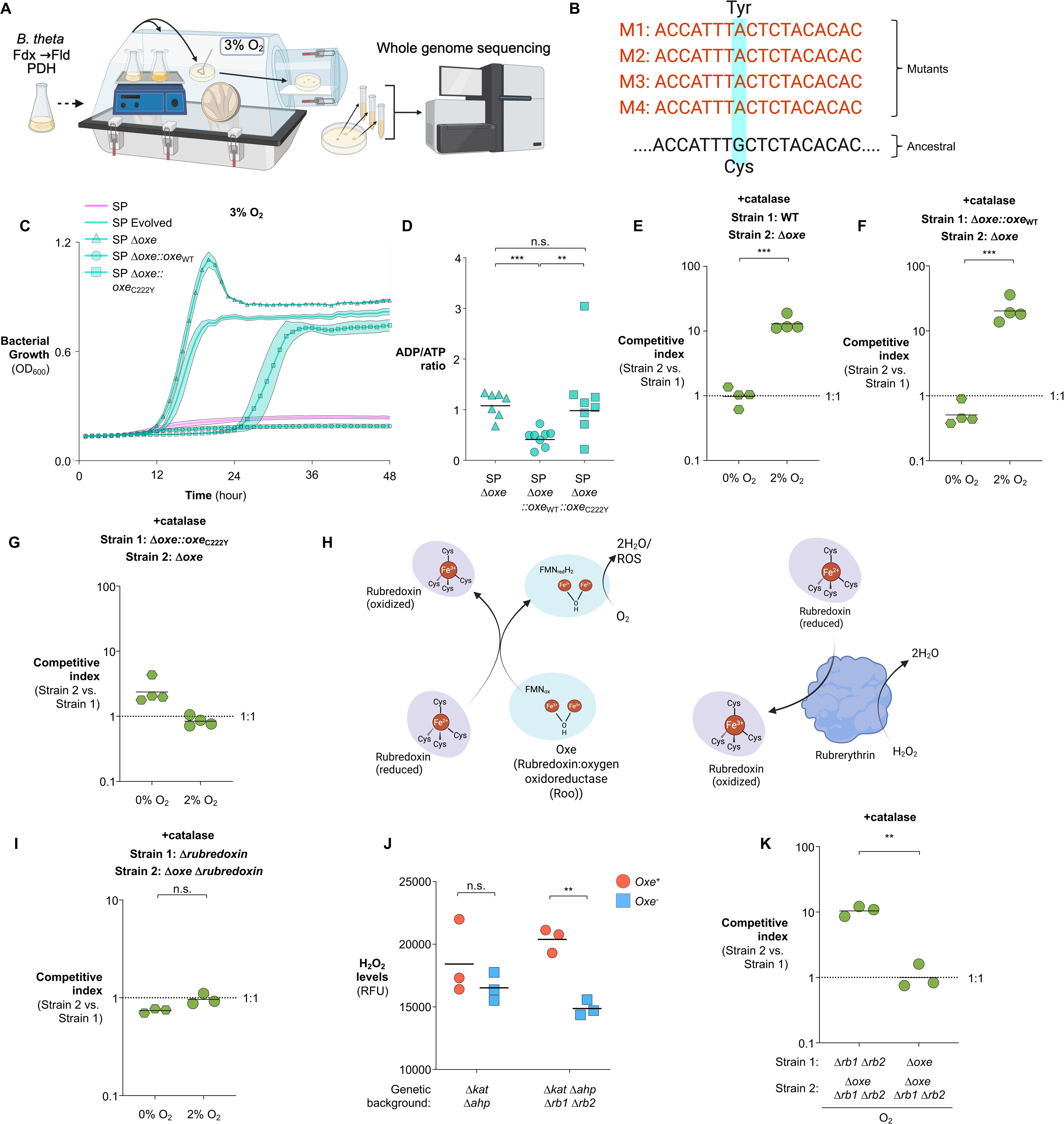
Oxe generates intracellular reactive oxygen species that limit microaerobic growth in *B. theta*. (**A-B**) The swap+PDH (SP) strain was experimentally evolved by repeated passaging in an open atmosphere containing 3% O_2_. Evolved populations were subjected to whole-genome sequencing to identify recurrent adaptive mutations. (**A**) Experimental schematic. (**B**) Mutations in *oxe* that arose reproducibly across all independently evolved isolates. (**C**) Growth of the indicated *B. theta* strains cultured in an open atmosphere containing 3% O_2_ with constant shaking, monitored by OD_600_. (**D**) The indicated *B. theta* strains were cultured for 16 h in an open atmosphere containing the indicated oxygen concentrations and intracellular ATP levels were quantified. (**E-G**) The indicated *B. theta* strains were mixed at a 1:1 ratio and cultured for 16 h under the indicated oxygen concentrations in an open atmosphere in the presence of extracellular catalase. Competitive indices were determined by selective plating. (**H**) Schematic model illustrating how Oxe may contribute to intracellular ROS generation during oxygen exposure. (**I**) Pairwise competition between rubredoxin-deficient and rubredoxin/oxe-deficient strains under the indicated oxygen conditions in the presence of catalase. (**J**) H_2_O_2_ accumulation in the indicated ROS-scavenging-deficient genetic backgrounds. (**K**) Pairwise competitions testing the interaction between oxe and rubrerythrins under oxic conditions in the presence of catalase. Bars represent geometric means. *, *P* < 0.05; **, *P* < 0.01; ***, *P* < 0.001.

Consistent with this result, prior studies have linked *oxe* inactivation to increased oxygen tolerance in *B. fragilis*^66^. Notably, we were unable to isolate any mutants when the wild-type strain was exposed to 3% O_2_, suggesting that prior genetic modifications in the SP strain were necessary to enable adaptation to microaerophilic conditions. To directly test the functional role of *oxe*, we generated a non-polar deletion mutant (Δ*oxe*) and complemented it with either the wild-type or mutant (C222Y) allele of *oxe*. Deletion of *oxe* conferred a significant fitness advantage over the parental SP strain at 3% O_2_, an effect that was partially reversed by complementation with wild-type *oxe* but not the C222Y variant, despite comparable expression levels (**Fig. 3C and Supplementary Fig. S3C**). Consistent with a central role in metabolic constraint, deletion of *oxe* in the SP background (SPO) improved cellular energy status, as reflected by a reduced ADP/ATP ratio, whereas complementation with the wild-type, but not the C222Y allele, restored the low-energy phenotype (**Fig. 3D**). Together, these findings identify Oxe activity as a key constraint on anaerobic metabolism.

To examine the consequences of Oxe activity in its native context in the absence of previous genetic modifications, we performed analogous experiments in the wild-type strain. Deletion of *oxe* similarly conferred a fitness advantage at lower oxygen levels and reduced peroxide accumulation during monoculture (**Supplementary Fig. S3D-E**), indicating that Oxe imposes an oxygen-dependent constraint independent of prior engineering.

Although Δ*oxe* improved growth in monoculture, pairwise competition assays revealed a more complex relationship between Oxe activity and population fitness. Across genetic backgrounds and oxygen levels, the magnitude of the Δ*oxe* advantage varied, and complementation with wild-type *oxe* did not uniformly restore the expected fitness defect, even in the SP background where increased metabolic flux amplified oxygen-dependent stress (**Supplementary Fig. S3F-K**). In contrast, strains complemented with the catalytically impaired *oxe* C222Y allele generally behaved more similarly to Δ*oxe* strains, consistent with reduced Oxe activity in this variant. Together, these findings suggested that cell-intrinsic growth capacity alone was insufficient to explain competitive fitness under oxic conditions and raised the possibility that Oxe imposes a shared environmental burden. Because catalytically active Oxe, but not the C222Y variant, was associated with peroxide production, we hypothesized that Oxe⁺ cells release H_2_O_2_ that diffuses through the population and constrains anaerobic growth. This model is consistent with the relatively long-lived and membrane-permeable nature of H_2_O_2_, in contrast to the short-lived, membrane-impermeable superoxide radical^67^.

To test this model, we repeated competition assays in the presence of membrane-impermeable catalase to selectively degrade extracellular hydrogen peroxide^68^. Catalase supplementation restored the competitive advantage of Δ*oxe* cells over Oxe⁺ strains (**Fig. 3E-G**), supporting the model that Oxe-derived H_2_O_2_ diffuses into the extracellular milieu and constrains anaerobic growth under oxic conditions.

Bioinformatic analysis revealed that Oxe shares sequence and structural homology with both nitric oxide reductase (NorV) and rubredoxin:oxygen oxidoreductase (Roo)^66^. Because nitric oxide was absent from our experimental conditions, we reasoned that Oxe may function as a Roo-like enzyme, transferring electrons to molecular oxygen (**Fig. 3H**), with incomplete reduction generating reactive oxygen species^69^. Consistent with rubredoxin acting upstream of Oxe, deletion of rubredoxin abolished the fitness advantage conferred by *oxe* loss under microaerobic conditions (**Fig. 3I**). To test whether Oxe promotes peroxide accumulation while minimizing confounding effects from endogenous scavenging systems, we compared H_2_O_2_ accumulation in a ROS-scavenging-deficient background (Δ*katE* Δ*ahpC* Δ*rbr1* Δ*rbr2*, Hpx^−^)^70^. Under 3% O_2_, *oxe*⁺ cells rapidly accumulated ROS whereas the Δ*oxe* mutant did not, an effect that became most apparent when rubrerythrins were removed (**Fig. 3J**), identifying Oxe as a key intracellular source of peroxide under oxic stress.

A non-mutually exclusive mechanism is that Oxe competes with rubrerythrins for rubredoxin-derived electrons^71^ (**Fig. 3H**). Because rubrerythrins reduce H_2_O_2_ to water^72,73^, Oxe could act as an electron sink that diverts reducing equivalents away from ROS detoxification. Consistent with this model, endogenous rubrerythrins partially buffered Oxe-dependent peroxide accumulation at 3% O_2_ (Δ*katE* Δ*ahpC rb1^+^ rb2^+^*vs. Δ*katE* Δ*ahpC* Δ*oxe rb1^+^ rb2^+^*, **Fig. 3J & Supplementary Fig. S4A**), and deletion of rubrerythrins markedly reduced the fitness benefit conferred by loss of *oxe* (**Supplementary Fig. S4B-C**). However, *oxe* deletion still provided a competitive advantage in the rubrerythrin-deficient background in the presence of extracellular catalase (**Fig. 3K**), indicating that the benefit of *oxe* loss cannot be explained solely by redirecting electrons toward rubrerythrin-dependent peroxide detoxification. Importantly, these phenotypes were not attributable to altered *oxe* expression (**Supplementary Fig. S4D**).

Together, these data identify Oxe as a key intracellular source of H_2_O_2_ that imposes an oxygen-dependent redox burden on anaerobic growth. Oxe likely contributes to oxygen sensitivity through both ROS generation and competition with peroxide-detoxification pathways, with the relative contribution of each mechanism depending on cellular redox state and detoxification capacity. Notably, phylogenetic analysis revealed that *oxe* is broadly conserved across *Bacteroides* and other gut-associated anaerobes, with its presence correlating with oxygen sensitivity (**Supplementary Fig. S4E**), suggesting that Oxe-dependent ROS generation represents a conserved determinant of oxygen vulnerability in this lineage.

### Sustained hydrogen utilization boosts *B. theta* fitness in oxic conditions

Although eliminating Oxe-mediated ROS production extended oxygen tolerance, SPO growth remained compromised at higher oxygen tensions (5% O_2_, **Supplementary Fig. S4F**), indicating that additional oxygen-sensitive processes constrain aerobic growth. Because redox balance is central to anaerobic metabolism, we asked whether oxygen disrupts key redox-active pathways. Hydrogenases represent a major redox node^74^, catalyzing reversible interconversion between molecular hydrogen and intracellular redox carriers^75^. In anaerobes, endogenous hydrogenases often function in hydrogen evolution by oxidizing reduced ferredoxin, thereby disposing of excess reducing equivalents^75^. In contrast, hydrogen can also serve as an abundant electron source in the gut^76,77^, with the potential to support reducing power and energy conservation^78^. However, hydrogenases, including those encoded by *B. theta*^79^, are highly susceptible to inactivation by even trace amounts of oxygen^74^. This pronounced sensitivity arises from either oxidation of the [Fe-Fe] catalytic center^74,80^ or the formation of inhibitory oxygen adducts at the [Fe-Ni] active site, rendering hydrogen metabolism nonfunctional under oxic conditions^81^.

We therefore tested whether installing an oxygen-tolerant hydrogen-oxidizing activity could alleviate this redox constraint by introducing the [Ni-Fe] hydrogenase Hyd-1 from *E. coli* into *B. theta* under control of the native PFL promoter (**Supplementary Fig. S5A**). Rather than serving as a direct replacement for endogenous hydrogen-evolving enzymes, Hyd-1 is expected to function primarily as an oxygen-tolerant H_2_ oxidation module, supplying electrons during oxic stress. Hyd-1 maintains activity in the presence of oxygen by redirecting electrons stored in iron-sulfur relay clusters to reduce the inhibitory oxygen adduct^81–83^ (**Fig. 4A**). Consistent with this model, the Hyd-1-expressing strain retained hydrogen oxidation activity and exhibited elevated intracellular NADH levels even at 5% O_2_ (**Fig. 4B-C**).

**Fig. 4.**
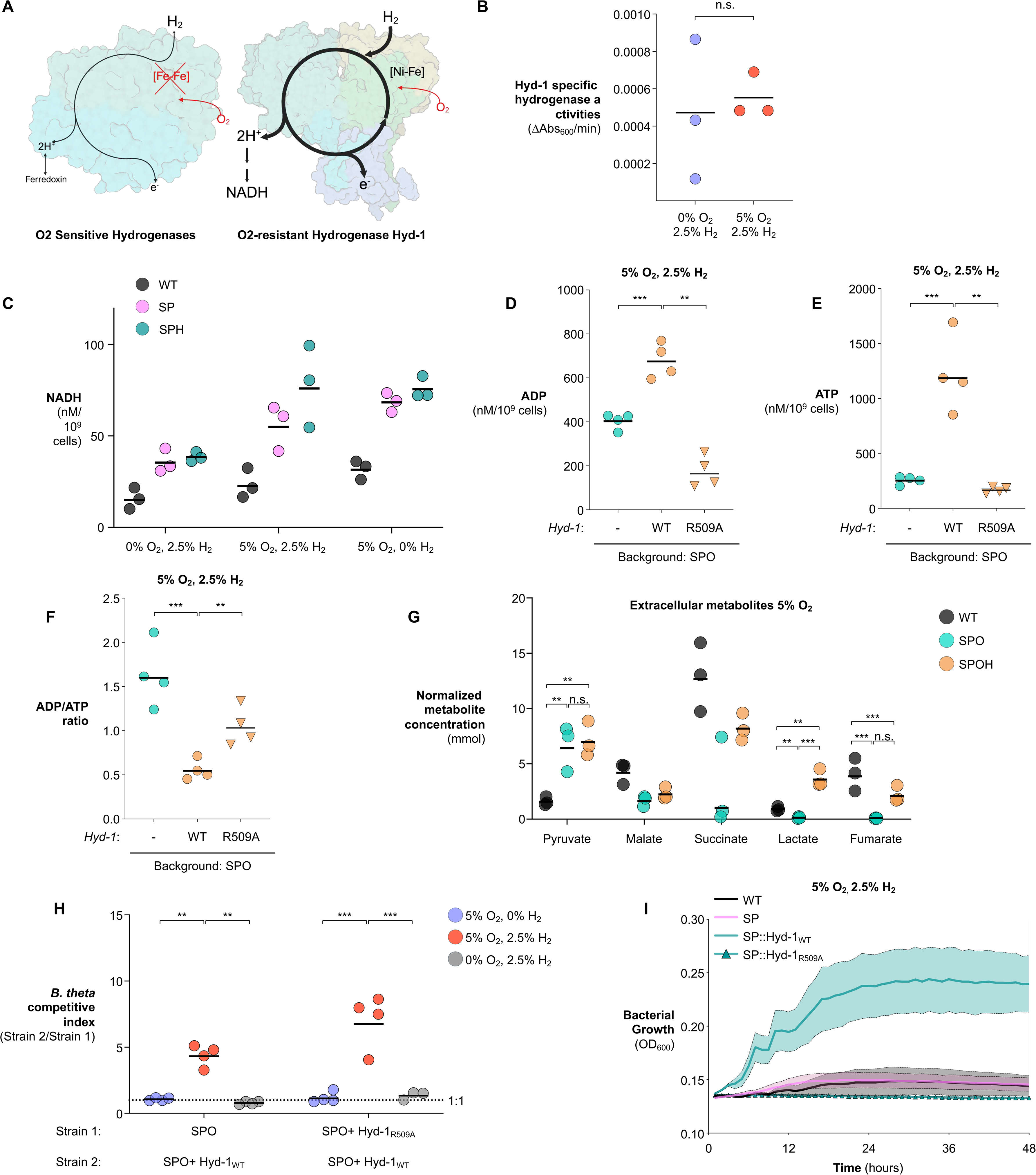
An oxygen-resistant hydrogenase preserves redox balance and improves *B. theta* fitness under oxic conditions. (**A**) Schematic comparison of oxygen-sensitive native hydrogenases and the oxygen-resistant [Ni-Fe] hydrogenase Hyd-1. (**B**) Hyd-1-specific hydrogenase activity measured under anaerobic conditions or following exposure to 5% O_2_ in the presence of 2.5% H_2_. (**C**) Intracellular NADH levels in the indicated strains under anaerobic or oxic conditions with or without H_2_. (**D-F**) Intracellular ADP (**D**), ATP (**E**), and ADP/ATP ratio (**F**) in the indicated SPO-background strains expressing no Hyd-1, wild-type Hyd-1, or catalytically inactive Hyd-1(R509A) following exposure to 5% O_2_ and 2.5% H_2_. (**G**) Extracellular TCA-related metabolites in indicated strains following exposure to 5% O_2_. (**H**) Pairwise competition assays between the indicated strains cultured under 0% or 5% O_2_ with or without 2.5% H_2_; competitive indices were determined by selective plating. (**I**) Growth of the indicated strains under 5% O_2_ with constant shaking, monitored by OD_600_. Bars represent geometric means. *, *P* < 0.05; **, *P* < 0.01; ***, *P* < 0.001.

We next introduced Hyd-1 into the SPO background, generating the SPOH strain (Swap + PDH + Δ*oxe* + Hyd-1), to determine whether hydrogen oxidation could improve cellular energy status after removal of Oxe-dependent ROS generation. Under 5% O_2_ and 2.5% H_2_, SPOH showed increased ATP production and a reduced ADP/ATP ratio relative to the parental SPO strain (**Fig. 4D-F**). These effects required both H_2_ and Hyd-1 catalytic activity, as hydrogen withdrawal or expression of an inactive Hyd-1 mutant (R509A) largely abolished the energetic benefits conferred by Hyd-1 expression (**Fig. 4D-F, Supplementary Fig. S5B**). Targeted analysis of TCA-cycle metabolites further showed that SPOH partially restored extracellular succinate, lactate, fumarate, and malate accumulation during oxygen exposure, despite continued pyruvate overflow (**Fig. 4G**). Similar oxygen-dependent buffering of central metabolite perturbations was observed in an independent WT versus SPH comparison (**Supplementary Fig. S5C-G**), supporting a general role for oxygen-tolerant hydrogen oxidation in stabilizing anaerobic metabolism under oxic stress.

We next asked whether preserving hydrogen utilization translated into a measurable fitness advantage under oxic conditions. In competition experiments performed in the presence of 2.5% H_2_ and 5% O_2_, SPOH significantly outcompeted both the parental SPO strain and the corresponding strain expressing the catalytically inactive Hyd-1 R509A variant (**Fig. 4H**). Consistent with this, expression of wild-type Hyd-1—but not the R509A mutant—conferred a growth advantage in monoculture under the same oxic conditions (**Fig. 4I**). Importantly, Hyd-1 expression had no measurable impact on growth under anaerobic conditions (**Supplementary Fig. S5H**), indicating that its functional contribution is specific to oxic stress. Together, these findings show that oxygen-tolerant hydrogen oxidation can provide a beneficial redox input during oxidative stress, enhancing *B. theta* fitness in oxygenated environments. These results prompted us to ask whether oxygen also disrupts essential anabolic pathways whose failure cannot be compensated by restoring redox nodes alone.

### Oxygen exposure impairs carbamoyl phosphate biosynthesis through widespread cysteine perturbation

Despite sequential repair of electron transfer, pyruvate assimilation, ROS generation, and hydrogen oxidation, engineered strains remained growth-limited at higher oxygen tensions (**Supplementary Fig. S6A**), suggesting the existence of additional oxygen-sensitive nodes. We therefore asked whether oxygen also disrupts anabolic enzymes through changes in cysteine reactivity. Cysteine residues play central roles in catalysis, metal coordination, and protein structure, and can be altered by direct oxidation, metal-cofactor disruption, or oxygen-induced conformational remodeling^84,85^ (**Fig. 5A**). Whether oxygen-induced cysteine perturbations contribute to the incompatibility between anaerobic metabolism and oxygen remains largely unresolved.

**Fig. 5.**
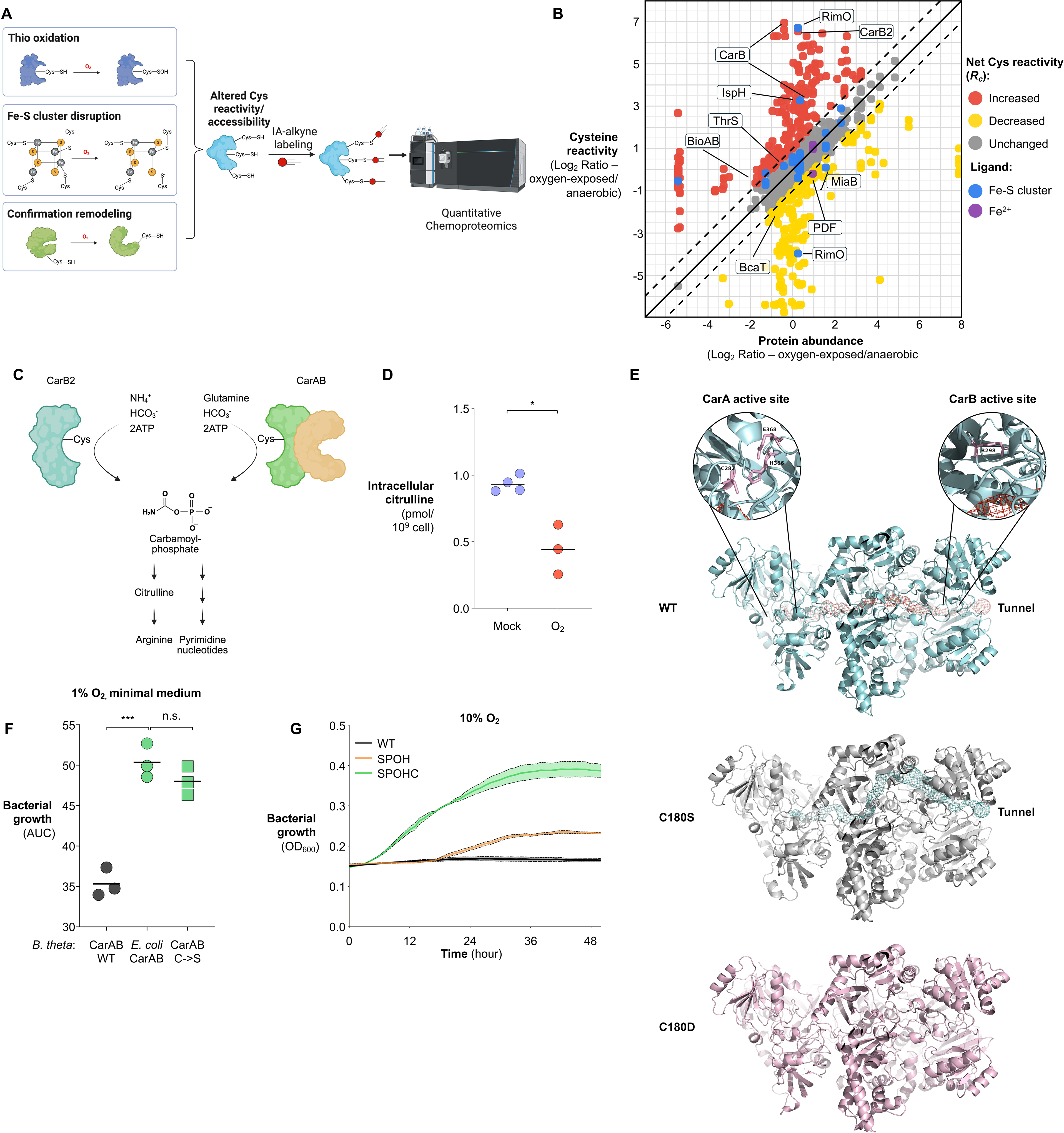
Oxygen impairs carbamoyl phosphate biosynthesis through cysteine modification in *B. theta*. (**A-B**) Anaerobically growing cultures of wild-type *B. theta* were either maintained under anaerobic conditions or exposed to 4% O_2_ for 90 minutes. Cell lysates were then labeled with isotopic iodoacetamide-alkyne (IA-alkyne) probes (mock: light; O_2_-exposed: heavy), a cysteine-reactive chemical probe, to quantify proteome-wide cysteine reactivity by mass spectrometry. After normalization to protein abundance, cysteine log_2_ (L/H) ratios (*R_C_*) report relative changes in labeling, with *R_C_* > 1 indicating increased cysteine reactivity upon oxygen exposure. (**A**) Schematic of the isoTOP-ABPP experimental workflow. (**B**) Two-dimensional proteomic plot of quantified cysteine residues from the *B. theta* proteome, highlighting cysteines with significantly altered reactivity upon oxygen exposure. (**C**) Schematic representation of carbamoyl phosphate biosynthesis and its integration into arginine and pyrimidine biosynthetic pathways. (**D**) Intracellular citrulline levels in wild-type *B. theta* following exposure to 10% O_2_ in the presence of NaHCO_3_ and glutamine to support carbamoyl phosphate synthesis. (**E**) Structural models of *B. theta* CarAB and predicted substrate tunnel architecture in wild-type, C180S, and C180D variants generated using CAVER. Insets highlight the CarA and CarB active sites. (**F**) Growth, quantified as area under the curve, of strains expressing endogenous CarAB, *E. coli* CarAB, or the CarAB C180S variant in wild-type background cultured in minimal medium at 1% O_2_. (**G**) Growth of the indicated strains cultured under 10% O_2_ with constant shaking, monitored by OD_600_. Bars represent geometric means. *, *P* < 0.05; **, *P* < 0.01; ***, *P* < 0.001.

Recent advances in high-throughput chemoproteomic profiling by mass spectrometry^86^ have enabled systematic analysis of stress-induced alterations in cysteine reactivity^87–90^. These approaches employ cysteine-reactive probes that exhibit reduced labeling when target cysteines undergo direct modification (e.g., oxidation or nitrosation). Conversely, disruption of metal cofactors such as Fe-S clusters can expose coordinating cysteines and increase their apparent reactivity^87,91,92^. Alternatively, oxidative perturbations can alter cysteine accessibility indirectly through conformational remodeling or changes in the local protein microenvironment^93–95^ (**Fig. 5A**).

To determine whether oxygen disrupts metabolic pathways through cysteine perturbation in *B. theta*, we applied a modified isotopic tandem orthogonal proteolysis-activity-based protein profiling (isoTOP-ABPP) platform^90^. This method employs iodoacetamide-alkyne (IA-alkyne), a cysteine-reactive chemical probe, to quantify changes to proteome-wide cysteine reactivity during oxygen exposure. Proteomes from anaerobic-versus 4% oxygen-exposed cultures are differentially labeled with isotopic IA-alkyne probes (light vs. heavy) and co-analyzed by mass spectrometry. After normalization to protein abundance, net log_2_ cysteine reactivity ratios (*R*_C_) report on relative changes in IA-alkyne labeling, with *R*_C_ > 1 indicating increased cysteine reactivity under oxygen exposure and *R*_C_ < 1 indicating decreased cysteine reactivity (**Fig. 5A**).

Remarkably, exposure to 4% oxygen altered the reactivity of 682 cysteine residues (392 with increased reactivity and 290 with decreased reactivity) across 376 proteins, including homologues of known oxygen-sensitive Fe-S proteins such as fumarase^96^. Several Fe-S-coordinating cysteines, including C97 in 4-hydroxy-3-methylbut-2-enyl diphosphate reductase (IspH, *R_C_* = 2.92), and C70 in lipoyl synthase (LipA, *R_C_* = 4.89) exhibited increased reactivity, consistent with oxygen-dependent cluster disruption (**Fig. 5B, Supplementary Fig. S6B, Supplementary Table 1**). In addition to canonical Fe-S-associated cysteines, oxygen exposure altered cysteine reactivity in enzymes lacking annotated metal-coordinating motifs (**Supplementary Fig. S6B)**, suggesting that oxygen perturbs anaerobic metabolism through broader structural and regulatory effects beyond classical cofactor damage.

Integration of the chemoproteomic, metabolomic, and transcriptomic datasets revealed convergent perturbation of a limited number of biosynthetic pathways, most prominently carbamoyl phosphate-dependent arginine and pyrimidine metabolism. Consistent with this, oxygen exposure altered cysteine reactivity in multiple enzymes linked to these pathways while simultaneously inducing compensatory transcriptional responses and accumulation of upstream metabolic intermediates (**Supplementary Fig. S1C-I, Supplementary Fig. S6C-D**). Together, these data suggested disruption of a shared metabolic node linking these pathways.

A central candidate emerging from this analysis was carbamoyl phosphate synthetase (CarAB), a heterodimer enzyme composed of a small (CarA) and a large subunit (CarB) that catalyzes the synthesis of carbamoyl phosphate, the shared precursor for pyrimidine and arginine biosynthesis (**Fig. 5C**). Because carbamoyl phosphate is chemically labile and difficult to quantify directly^97,98^, we supplemented cells with NaHCO_3_ and glutamine to support carbamoyl phosphate production and measured intracellular citrulline, a downstream product of carbamoyl phosphate utilization in arginine biosynthesis^99^ (**Fig. 5C**). Oxygen exposure markedly reduced citrulline abundance (**Fig. 5D**), consistent with impaired carbamoyl phosphate-dependent metabolic activity. Supporting this interpretation, a genome-scale metabolic model constrained by the metabolomic dataset predicted reduced carbamoyl phosphate-dependent flux together with increased flux through arginine and pyrimidine salvage pathways during oxygen exposure (**Supplementary Fig. S7A-D**). These changes are consistent with a compensatory metabolic shift away from *de novo* carbamoyl phosphate-dependent biosynthesis toward salvage metabolism during oxygen stress.

To determine whether this defect contributes to oxygen sensitivity, we supplemented arginine to bypass the blocked pathway. Arginine supplementation reduced accumulation of upstream intermediates (e.g., N-acetylglutamate) and improved growth under moderate oxygen (3% O_2_, **Supplementary Fig. S7E-F**) but failed to restore growth at higher oxygen levels (5% and 10% O_2_, **Supplementary Fig. S7G-H**), indicating that upstream dysfunction remained limiting. Supplementation with pyrimidines or dNTPs was similarly ineffective (**Supplementary Fig. S7H**), likely reflecting insufficient transport capacity to meet cellular demand.

We therefore sought to address the root cause of arginine and pyrimidine deficiency by repairing carbamoyl phosphate biosynthesis itself. Direct addition of carbamoyl phosphate did not restore growth (**Supplementary Fig. S7H**), likely reflecting its short half-life^98^ (∼5 mins at 37 °C). Instead, we focused on CarAB, which exhibited oxygen-sensitive cysteine reactivity at CarB C180 (*R_C_* = 4.41, **Fig. 5B**). CarAB subunits are connected by an internal ammonia channel that transfers ammonia generated by CarA to the catalytic site of CarB, enabling efficient carbamoyl phosphate synthesis (NH ^+^ + HCO ^−^ + ATP-> H NCO P^−^ + ADP)^100,101^. Substrate tunnel predictions (CAVER^102^) confirmed the presence of this inter-subunit conduit. Notably, substituting C180 with aspartate, which introduces a charged side chain that models perturbation at this position, disrupted predicted channel continuity, whereas substitution with serine preserved the channel architecture (**Fig. 5E**).

Next, we tested whether repairing this oxygen-sensitive feature of CarAB could enhance oxic growth. We reasoned that adaptation to oxygen may have favored acquisition of oxygen-resistant variants of key biosynthetic enzymes such as carbamoyl phosphate synthetase^103^. Consistent with this, the oxygen-responsive cysteine is highly conserved among obligate anaerobes, whereas non-reactive residues (e.g., alanine or valine) are conserved in aerotolerant lineages (**Supplementary Fig. S7I**), suggesting evolutionary selection against oxygen-sensitive features.

To directly test this model, we replaced the endogenous *carAB* with the facultative anaerobe *E. coli* ortholog. This substitution did not alter overall fitness under anaerobic conditions (**Supplementary Fig. S7J**) but conferred a competitive fitness advantage under oxic conditions in rich medium (wild-type background, **Supplementary Fig. S7K**). We next examined CarAB function in *Bacteroides* minimal medium, where cells rely more heavily on endogenous carbamoyl phosphate synthesis. Under these conditions, replacement of endogenous CarAB with *E. coli* CarAB or the oxygen-resistant C180S variant significantly improved growth at 1% O_2_ in a wild-type background (**Fig. 5F**), supporting the idea that oxygen-sensitive perturbation of CarAB contributes directly to impaired anabolic growth during oxygen exposure. Strikingly, introduction of the *E. coli* CarAB (SPOHC) into the previously engineered background enabled robust growth at 10% O_2_, a condition under which all previously engineered strains failed to thrive (**Fig. 5G**). Together with prior repairs to central metabolism and redox balance, these findings establish oxygen-sensitive carbamoyl phosphate synthesis as a key bottleneck limiting anaerobe growth at elevated oxygen tensions.

### Conditional cofactor limitation contributes to *B. theta* oxygen sensitivity

Even after repairing carbamoyl phosphate synthesis, growth at elevated oxygen remained incomplete, suggesting that additional metabolic processes become conditionally limiting during oxygen exposure. Many essential metabolic reactions depend on vitamin-derived cofactors, whose biosynthesis and utilization can be vulnerable to oxygen-sensitive chemistries, including Fe-S clusters, radical-mediated reactions, and redox-active protein states^104,105^. We therefore asked whether cofactor availability becomes conditionally limiting during oxygen stress in *B. theta*.

Integrated transcriptomic and untargeted metabolomic analyses revealed perturbation of vitamin-associated pathways during oxygen exposure (**Supplementary Fig. S1C-D, H-I**), including pantothenate (vitamin B5) and pyridoxal 5′-phosphate (vitamin B6) metabolism. Several enzymes connected to these pathways, including branched-chain amino acid aminotransferase and threonine synthase, also harbored cysteines with altered reactivity upon oxygen exposure (**Fig. 5B and Supplementary Table S1**). Although these data do not identify a specific limiting enzymatic step, they suggested that oxygen stress may create a relative cofactor limitation by impairing biosynthesis, increasing cofactor demand, perturbing utilization, or altering uptake.

We next tested whether vitamin supplementation could alleviate this conditional limitation. Under anaerobic conditions, supplementation with either vitamin had no measurable impact on fitness (**Supplementary Fig. S8A**), consistent with sufficient endogenous production and/or availability. In contrast, under 5% O_2_, vitamin B5 supplementation accelerated growth of the *B. theta* SPOH strain (**Fig. 6A**). This was accompanied by reduced ADP levels, while ATP levels and the ADP/ATP ratio were not significantly altered (**Fig. 6B-D**), suggesting that B5 availability improves metabolic performance under moderate oxygen stress without broadly increasing cellular energy charge. However, B5 supplementation failed to enhance fitness at 10% O₂ (**Supplementary Fig. S8C**), indicating that additional oxygen-imposed bottlenecks become dominant at higher oxygen tensions.

**Fig. 6.**
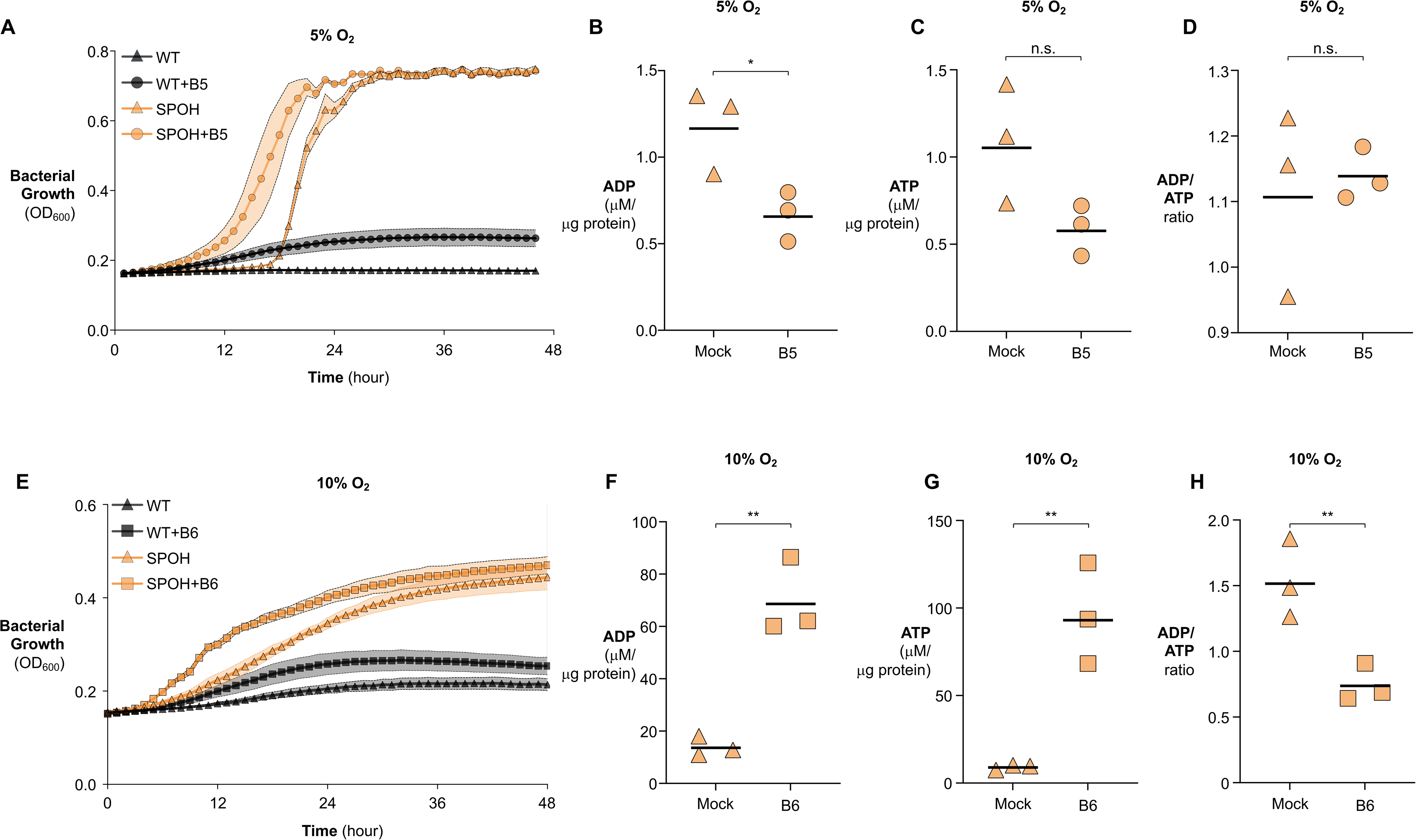
Oxygen impairs vitamin B5 and B6 biosynthesis in *B. theta*. (**A-D**) The indicated *B. theta* strains were cultured under 5% O_2_ with or without vitamin B5 supplementation. (**A**) Growth monitored by OD_600_. (**B-D**) Intracellular ADP (**B**), ATP (**C**), and ADP/ATP ratio (**D**) measured in SPOH cultures with or without vitamin B5 supplementation. (**E–H**) The indicated B. theta strains were cultured under 10% O_2_ with or without vitamin B6 supplementation. (**E**) Growth monitored by OD_600_. (**F-H**) Intracellular ADP (**F**), ATP (**G**), and ADP/ATP ratio (**H**) measured in SPOH cultures with or without vitamin B6 supplementation. For growth curves, lines represent geometric means and shaded regions indicate SEM. For endpoint measurements, bars represent geometric means. n.s., not significant; *, P < 0.05; **, P < 0.01.

Vitamin B6 supplementation had a stronger effect, enabling robust SPOH growth at 10% O_2_ (**Fig. 6E**). This rescue was accompanied by increased ATP and a reduced ADP/ATP ratio, despite elevated ADP levels (**Fig. 6F-H**), indicating increased metabolic activity and energy production under severe oxygen stress. This effect was not attributable to antioxidant activity^106^, as vitamin B6 did not alter H_2_O_2_ production (**Supplementary Fig. S8D**). Instead, the rescue is consistent with improved availability or function of B6-dependent metabolic processes during oxygen exposure. Together, these findings suggest that vitamin-derived cofactor availability can become conditionally limiting during oxidative stress, even in complex medium, and point to cofactor-dependent metabolism as an additional vulnerability that may constrain growth at elevated oxygen tensions.

### Rational engineering of facultative anaerobiosis in *B. theta* bolsters its resilience in the inflamed gut

Our genetic and metabolic interventions reveal that oxygen sensitivity in *B. theta* arises from multiple interlocking vulnerabilities, including excessive intracellular ROS generation and the oxygen lability of enzymes involved in redox balance, cofactor utilization, and central metabolic pathways. Sequential repair of these weak points progressively expanded the capacity of *B. theta* to grow in the presence of oxygen *in vitro* (**Figs. 1-6**). Importantly, these stepwise gains were observed reproducibly in pairwise competition assays specifically under oxic conditions (**Supplementary Fig. S8E**), supporting the conclusion that the engineered modifications, rather than stochastic suppressor outgrowth, are the primary drivers of enhanced oxygen tolerance.

In contrast to the homeostatic, anaerobic intestinal lumen, the inflamed intestine poses pronounced oxidative challenges. This arises from a metabolic shift in colonocytes toward anaerobic glycolysis, increasing oxygen diffusion into the otherwise anoxic gut lumen^10–14^. In parallel, immune activation generates reactive oxygen species (ROS), further amplifying oxidative stress for resident anaerobes^12,16–18^. Given that our engineering strategy conferred stepwise gains in oxygen tolerance *in vitro* (**Supplementary Fig. S8E**), we next asked whether these repairs enhance *B. theta* resilience in the oxygenated, inflamed gut.

To test this, we performed competitive colonization experiments in antibiotic-pretreated C57BL/6 mice to permit engraftment^107^. Mice were inoculated with an equal mixture of two *B. theta* strains, where the second strain (strain B) carried additional oxygen-resistance repairs relative to its otherwise isogenic counterpart (strain A; e.g., WT vs. S, S vs. SP). One day after colonization, mice were challenged with *Salmonella enterica serovar* Typhimurium (*S*. Tm), which induces robust gut inflammation and oxygen influx into the lumen via its Type III Secretion System (T3SS)^11,108^. Consistent with a previous report^11^, *S*. Tm infection drove oxygenation in the large intestine (**Fig. 7A-B**).

**Fig. 7.**
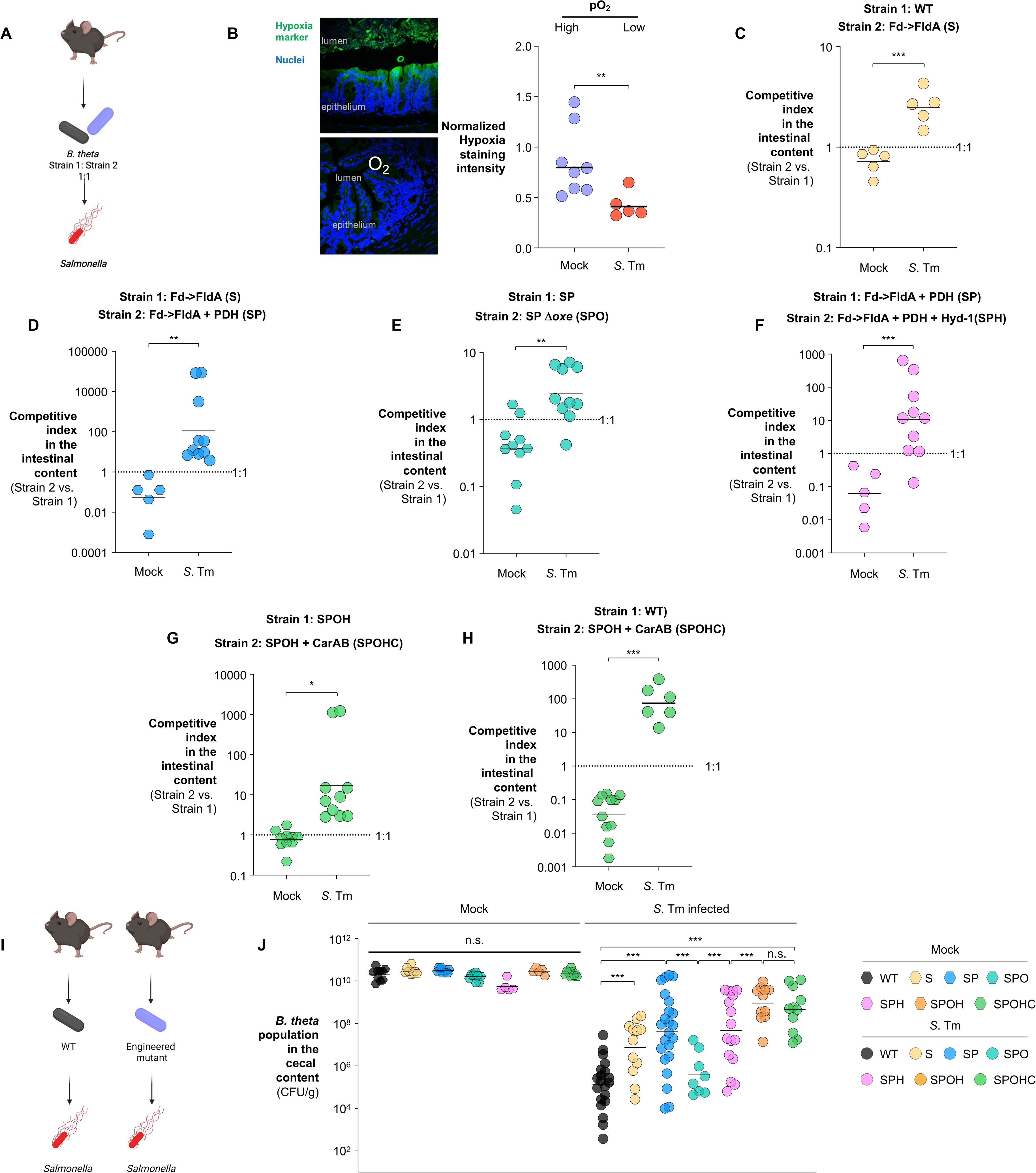
Stepwise enhancement of oxygen tolerance progressively improves *B. theta* resilience in the oxygenated, inflamed gut. (**A**) Schematic of the competitive colonization experimental design. Antibiotic-pretreated C57BL/6 mice were inoculated with a 1:1 mixture of two *B. theta* strains (strain 1 and strain 2), followed by mock treatment or infection with *Salmonella* Typhimurium (S. Tm) SL1344. (**B**) Representative images of tissue hypoxia in the cecum assessed by pimonidazole staining (green) with DAPI nuclear counterstaining (blue). (**C-H**) Competitive colonization assays in which mice were inoculated with equal mixtures of isogenic strains differing by sequential oxygen-resistance engineering steps and analyzed 4 days post-infection; competitive indices (strain 2 vs. strain 1, normalized to input) were determined from cecal contents. (**I-J**) Antibiotic-pretreated C57BL/6 mice were intragastrically inoculated with a single *B. theta* strain (either strain A or an isogenic derivative that carries additional oxygen-resistance modifications, strain B) and subsequently challenged with *S*. Tm SL1344. (**I**) Schematic of the single-strain colonization experiment. (**J**) Four days after infection, cecal contents were collected and bacterial abundances were quantified by selective plating. Bars represent geometric means. *, *P* < 0.05; **, *P* < 0.01; ***, *P* < 0.001.

Replacing ferredoxin with flavodoxin imposed a modest fitness cost under homeostatic conditions but conferred a larger competitive advantage during inflammation (WT vs. S; **Fig. 7C**), consistent with reduced ROS generation at the expense of less efficient electron transfer^109^. Introduction of an oxygen-tolerant pyruvate dehydrogenase complex provided a substantially greater benefit during inflammation (S vs. SP, **Fig. 7D**), highlighting impaired pyruvate assimilation as a major oxygen-imposed constraint *in vivo*.

In contrast, deletion of *oxe*, which markedly enhances oxygen tolerance in vitro, conferred only a modest fitness advantage during *S*. Tm infection (SP vs. SPO; **Fig. 7E**). This discordance indicates that Oxe exerts opposing effects *in vivo*. While it contributes to oxygen-dependent ROS burden, it may also support redox functions that become advantageous under inflammatory stress. Notably, Oxe shares homology with nitric oxide reductase NorV and has been implicated in nitric oxide detoxification^66^, an abundant antimicrobial mechanism during *S*. Tm infection^110^. Because major nitrosative stress defense pathways in *B. theta* are iron-dependent^111^, this phenotype became most apparent under iron-limited conditions, where redundant NO-detoxification systems are likely compromised. Consistent with this role, *oxe* deletion impaired nitric oxide resistance (**Supplementary Fig. S9A**). Together, these findings suggest that Oxe mediates a tradeoff between ROS generation and protection against host-derived reactive nitrogen species, resulting in only a modest net fitness benefit of *oxe* loss during infection.

Introduction of an oxygen-tolerant hydrogenase improved fitness in the *oxe*⁺ background (SP vs. SPH), but not after *oxe* deletion (SPO vs. SPOH) (**Fig. 7F** and **Supplementary Fig. S9B**). This epistasis indicates that the benefit of hydrogen oxidation depends on Oxe, suggesting that hydrogen-derived reducing equivalents are funneled through an Oxe-dependent redox network rather than acting independently. Thus, hydrogenase function is advantageous only within an intact Oxe-dependent electron transfer network. Although hydrogenase conferred a fitness advantage only in *oxe*^+^ backgrounds in this infection model, we retained it in subsequent constructs because its benefit is expected to depend on *in vivo* H_2_ availability and electron-sink capacity, which vary across gut microenvironments and can shift as additional oxygen-sensitive nodes are repaired.

In contrast, repairing carbamoyl phosphate synthesis provided an additional fitness gain during *S*. Tm infection (**Fig. 7G**), demonstrating that oxygen-sensitive biosynthetic capacity remains limiting *in vivo*. Notably, cumulative repair of these vulnerabilities offset the fitness cost associated with *oxe* deletion, enabling the fully engineered strain (SPOHC) to markedly outcompete the wild-type strain during inflammation (**Fig. 7H**). These differences were not attributable to altered host responses, as *S*. Tm burdens and inflammatory markers were comparable across groups (**Supplementary Fig. S9C-K**).

We next evaluated the performance of engineered strains in the context of pathogen challenge and competition with the resident microbiota (**Fig. 7I**). Using a single-colonization model, C57BL/6 mice were colonized with individual *B. theta* strains and subsequently mock-treated or infected with *S*. Tm. Similar to the competitive colonization experiments, successive engineering steps progressively increased *B. theta* abundance in the inflamed gut, with the exception of hydrogenase engraftment and *oxe* deletion alone (**Fig. 7J**). Remarkably, combining ferredoxin replacement with oxygen-tolerant pyruvate dehydrogenase increased abundance by more than 300-fold during inflammation, while subsequent deletion of *oxe* and introduction of hydrogenase further enhanced expansion to approximately 7,000-fold (**Fig. 7J**). Notably, incorporation of oxygen-resistant CarAB into this background did not further increase abundance, despite its clear fitness benefit in competition assays, consistent with the possibility that the SPHO strain approaches an ecological limit in the inflamed gut under these conditions.

Across all conditions, *S*. Tm colonized to similar levels and caused comparable gross inflammatory pathology, as reflected by colon shortening (**Supplementary Fig. S10A&B**). Although several inflammatory transcripts varied across groups (**Supplementary Fig. S10C-H**), these differences did not consistently track with the magnitude of engineered *B. theta* expansion, indicating that improved colonization was not simply explained by reduced pathogen burden or globally attenuated inflammation. Together, these results identify oxygen and ROS as dominant ecological pressures limiting obligate anaerobes in the inflamed gut and demonstrate that rational engineering of facultative anaerobiosis can fortify commensal resilience *in vivo*.

## Discussion

Following the Great Oxygenation Event, many organisms evolved strategies to tolerate oxygen by eliminating oxygen-labile enzymes, developing oxygen-resistant counterparts, or maintaining intracellular ROS at non-toxic levels^16^. Although billions of years of evolution have solved these challenges in aerobes and facultative anaerobes^33^, the molecular constraints that limit obligate anaerobes in the presence of oxygen remain incompletely defined. Here, by integrating transcriptomics, metabolomics, chemoproteomics, and genome-scale metabolic modeling, we identify multiple, interlocking molecular vulnerabilities that collectively impose an aerobic growth barrier in the strict anaerobe *Bacteroides thetaiotaomicron*. We demonstrate that rational repair of these vulnerabilities progressively enhances oxygen tolerance, enabling robust growth in up to 10% O_2_ and markedly improving fitness in the oxygenated, inflamed gut (**Figs. 1-7**). Together, these findings shift the problem of obligate anaerobiosis from a descriptive trait to a repairable metabolic problem set.

Repair of central carbon metabolism revealed a hierarchy of oxygen-sensitive constraints. Replacing the oxygen-labile, Fe-S-dependent electron carrier ferredoxin with flavodoxin modestly improved oxygen tolerance, consistent with reduced ROS generation at the cost of less efficient electron transfer (**Fig. 1**). In contrast, introduction of an oxygen-resistant pyruvate dehydrogenase (PDH) complex produced a substantially greater benefit (**Fig. 2**), identifying pyruvate assimilation as a dominant oxygen-imposed metabolic constraint. We initially hypothesized that this modification might impose a fitness cost under anaerobic conditions, because the native pyruvate-splitting enzymes do not generate NADH^29,112,113^. In contrast, the additional NADH produced by PDH needs to be reoxidized to maintain redox balance, which could reduce ATP yield if excess NADH were reoxidized through non-energy-conserving processes.

However, when nutrients are abundant, the additional NADH generated by PDH can be accommodated by redirecting reducing equivalents into existing anaerobic redox sinks, including fumarate reduction to succinate^114^. This provides a route for NADH reoxidation^115–117^ and is consistent with increased glycolytic flux toward succinate in the PDH-expressing strain (**Fig. 2J**). This metabolic flexibility, together with the auto-inhibitory regulation of PDH by NADH^118^, likely explains why PDH can be tolerated. Under nutrient-limiting conditions, however, where fumarate or other electron acceptor pools are constrained, excess NADH could lead to redox imbalance. This context dependence may help explain why obligate anaerobes typically rely on PFL and PFOR^16,119^, which avoid generating surplus NADH during pyruvate assimilation.

Beyond carbon flux, this study identifies Oxe as an important contributor to ROS generation and, therefore, to the incompatibility between anaerobic metabolism and oxygen^28^. In *B. theta*, Oxe likely transfers electrons from rubredoxin to oxygen, producing incompletely reduced intermediates (ROS), analogous to observations in *Desulfovibrio vulgaris*^69^. This contrasts with aerobic respiration, where cytochrome oxidases safely reduce oxygen to water, minimizing ROS formation^120^. In addition, Oxe may compete with rubrerythrins for rubredoxin-derived electrons, limiting peroxide detoxification and further amplifying oxidative stress. Why, then, is a ROS-generating enzyme retained in obligate anaerobes? Within its native niche, Oxe likely serves beneficial functions, including detoxification of reactive nitrogen species (**Supplementary Fig. S9A**), revealing a physiological tradeoff between ROS production and stress defense. This dual role suggests that Oxe is not simply a liability, but an evolutionary compromise optimized for life in predominantly anoxic environments. More broadly, these findings illustrate that oxygen sensitivity can emerge not only from intrinsic enzyme fragility, but also from tradeoffs in redox metabolism.

Our chemoproteomic analysis further reveals cysteine reactivity as a widespread and underappreciated determinant of oxygen sensitivity. Consistent with the enrichment of cysteine residues in anaerobic proteomes^121^, oxidative perturbation of cysteine residues extends beyond canonical metal-binding sites and affects diverse metabolic enzymes. Among these, we identify a cysteine (C180) in carbamoyl phosphate synthetase (CarB) as a key vulnerability. Structural modeling suggested that oxygen-dependent perturbation of the local environment surrounding C180 could influence the substrate tunnel connecting the CarA and CarB active sites, with potential consequences for substrate channeling and carbamoyl phosphate synthesis. Notably, this cysteine is replaced by valine in facultative anaerobes, consistent with evolutionary selection against oxygen-sensitive features. Functionally, replacement with *E. coli* CarAB or the C180S variant improved growth under oxygen exposure, supporting the idea that this site contributes to oxygen-sensitive pathway failure. Direct biochemical measurements will be required to determine whether this reflects increased intrinsic oxygen resistance of CarAB, altered substrate channeling, or improved pathway function *in vivo*. More broadly, CarAB represents one example of how redox-sensitive cysteines can encode latent liabilities in anaerobic enzymes, highlighting a general principle by which oxygen perturbs anaerobic metabolism.

Together, these findings establish an initial framework for understanding the molecular incompatibility between *B. theta* and oxygen. Rationally repairing vulnerabilities in central metabolism, ROS generation, and biosynthetic capacity yielded a *B. theta* strain capable of robust growth at 10% oxygen and markedly improved fitness in the oxygenated, inflamed gut. Although this study focuses on a single obligate anaerobe, many of the molecular liabilities identified here, e.g., oxygen-labile iron-sulfur enzymes, redox-active cysteine residues, and biosynthetic pathways optimized for anoxic metabolism, are broadly conserved across gut-resident and environmental anaerobes. These observations suggest that obligate anaerobiosis may arise not from a singular incompatibility with oxygen, but from the cumulative effect of multiple, interlocking vulnerabilities that are, as we demonstrated, repairable. Consistent with this view, our results demonstrate that facultative anaerobiosis can be experimentally enabled in a strict anaerobe within a tractable timeframe, recapitulating key evolutionary solutions to oxygen stress and expanding the ecological and therapeutic potential of anaerobic commensals.

### Limitation of the study

While this work provides proof of principle that facultative anaerobiosis can be rationally engineered into an obligate anaerobe, the resulting *B. theta* strains remain unable to grow under fully atmospheric oxygen (20.9%), in contrast to naturally facultative anaerobes such as *E. coli*. This limitation indicates that additional oxygen-imposed metabolic constraints remain unresolved. Indeed, our integrated multi-omics datasets reveal numerous additional candidate vulnerabilities including enzymes involved in cell envelope and biosynthetic processes that may further contribute to oxygen sensitivity and warrant future investigation. Of note, chemoproteomic measurements of cysteine reactivity report relative changes in probe labeling rather than absolute cysteine occupancy. Thus, further validation of high-occupancy residues will be necessary to determine whether these changes correspond to functionally relevant, oxygen-sensitive alterations in cysteine reactivity. Although the pathways defined here are broadly conserved across *Bacteroides* species, it remains unclear whether these engineering strategies can be generalized across the genus or extended to other major gut commensal phyla, such as *Firmicutes*, which possess distinct metabolic architectures. Finally, while the engineered *B. theta* strains exhibited markedly improved fitness during enteric infection, future efforts will be required to refine these strains for therapeutic application. In particular, minimizing traits that may adversely impact host physiology, such as mucus degradation or metabolic cross-feeding to pathogens, will be essential to ensure safety and translational viability.

## STAR METHODS

### Bacterial strains, and growth conditions

*Bacteroides thetaiotaomicron* VPI-5482 was cultured under strictly anaerobic conditions (90% N_2_, 5% CO_2_, 5% H_2_; Coy vinyl anaerobic chamber) at 37 °C in brain heart infusion supplemented (BHIS) medium or on BHIS agar plates for 48 h. BHIS medium consisted of 0.8% brain heart infusion solids, 0.5% peptic digest of animal tissue, 1.6% pancreatic digest of casein, 0.5% sodium chloride, 0.2% glucose, 0.25% disodium hydrogen phosphate, 0.005% hemin, and 0.0001% vitamin K (pH 7.4). BHIS agar plates were prepared by supplementing BHIS broth with 15 g/L agar. Where indicated, media were supplemented with antibiotics at the following concentrations: erythromycin (25 μg/mL), tetracycline (6 μg/mL), chloramphenicol (15 μg/mL), or 5-fluoro-2′-deoxyuridine (FUdR; 200 μg/mL).

*Escherichia coli* strains were routinely cultured in LB broth (10 g/L tryptone, 5 g/L yeast extract, 10 g/L sodium chloride) or on LB agar plates (supplemented with 15 g/L agar) at 37 °C. Where appropriate, media were supplemented with carbenicillin (100 μg/mL).

For in vitro competition assays, overnight *B. theta* cultures grown from single colonies in BHIS medium for ∼18 h were mixed at equal ratios and inoculated at the indicated total density (typically 1 × 10^5^–10^6^ CFU/ml). Cultures were exposed to defined oxygen conditions in an open atmosphere (1–10% O_2_, 5% CO_2_, 2.5% H_2_, balance N_2_) at 37 °C with continuous shaking (200 rpm). After 16 h, cultures were serially diluted and plated on selective media to enumerate colony-forming units, and competitive indices were calculated as the ratio of strain 2 over strain 1 normalized to the input ratio.

For microaerobic growth assays, single colonies of the indicated *Bacteroides thetaiotaomicron* strains were cultured overnight in BHIS under anaerobic conditions. Cultures were inoculated at low density (1 × 10^5^ CFU/mL) into BHIS and cultured under defined oxygen conditions (1-10% O_2_, 5% CO_2_, 2.5% H_2_, balance N_2_) at 37 °C in an open hypoxia system (Coy Laboratory Products) with continuous shaking (200 rpm), using a BioTek Epoch microplate reader for up to 72 h. Low initial inocula were used to minimize collective oxygen scavenging^47^ and the resulting reduction in effective oxygen tension at high cell densities^48^. Bacterial growth was monitored by measuring optical density at 600 nm (OD_600_). To ensure adequate gas exchange, only the outer wells of the plate were used.

### Plasmids

All primers and plasmids used in this study are listed in the Key Resources Table. Suicide plasmids were routinely propagated in *E. coli* S17-λpir. For construction of the Δoxe allele in *B. theta* VPI-5482, 500 bp regions upstream and downstream of *oxe* were amplified and assembled into pExchange-tdk using Gibson Assembly (New England Biolabs), yielding plasmid pAR8.

For genomic integration constructs (pKIs; e.g., pKI1, pKI2), flanking regions of two convergently oriented genes were amplified and assembled into the counter-selectable allelic exchange vector pExchange^122^. The resulting plasmids were linearized to introduce DNA fragments of interest, enabling single-copy genomic integration of expression cassettes at defined loci.

To replace oxygen-sensitive genes with oxygen-tolerant variants, flanking regions of the target gene (extending to the start and stop codons) were amplified and assembled into pExchange as described above. The resulting constructs were subsequently linearized to introduce the replacement gene, generating allelic exchange vectors for precise gene substitution.

### Construction of mutants by allelic exchange

All mutant strains generated in this study are listed in the Key Resources Table. For gene deletions in *B. theta*, suicide plasmids (pExchange) carrying 500 bp flanking regions of the target locus were introduced by conjugation using *E. coli* S17-λpir as the donor strain. For genomic insertions, pKI plasmids harboring the gene(s) of interest were similarly mobilized into *B. theta* via conjugation.

Following conjugation, single-crossover integrants were selected on BHIS agar supplemented with the appropriate antibiotics. Counterselection for the second homologous recombination event was performed on BHIS plates containing 5-fluoro-2′-deoxyuridine (FUdR; 200 μg/mL), yielding either deletion mutants or allelic exchange strains. Colonies were screened by PCR using locus-specific verification primers to confirm the desired genotype.

### ATP and ATP/ADP measurements

Intracellular ATP and ATP+ADP levels were quantified using a luciferase-based BacTiter-Glo Microbial Cell Viability Assay (Promega), with optional enzymatic conversion of ADP to ATP. Bacterial cultures were grown anaerobically or exposed to defined oxygen conditions as described above, in an open atmosphere containing 1–10% O_2_, 5% CO_2_, 2.5% H_2_ (balance N_2_) at 37 °C. Cultures were maintained in a 24 well plate with a large air-liquid interface, sealed with a gas permeable sealing membrane (Breathe-Easy, Sigma Aldrich), and continuous shaking (200 rpm) to facilitate efficient gas exchange. At indicated time points, cells were rapidly chilled on ice to quench metabolism and normalized to 1 × 10^6^ cells/mL prior to analysis.

For ATP measurements, 25 μL of normalized culture or metabolite extract was dispensed into white 96-well plates, followed by addition of an equal volume of BacTiter-Glo reagent. Plates were mixed briefly, incubated at room temperature for 5 min, and luminescence was measured using a plate reader. ATP concentrations were determined using a standard curve generated from serial dilutions of ATP.

For combined ATP+ADP measurements, metabolites were first extracted by pelleting 1 mL of normalized culture and quenching in ice-cold 80% methanol. Extracts were clarified, dried, and reconstituted in Tris buffer (50 mM, pH 7.5). ADP was enzymatically converted to ATP by incubation with pyruvate kinase in the presence of phosphoenolpyruvate and Mg^2+^ (final concentrations: 5 mM PEP, 5 mM MgCl_2_, 2 U/mL pyruvate kinase) for 10 min at room temperature prior to addition of BacTiter-Glo reagent. Luminescence from these reactions reflects total ATP+ADP, and ADP levels were calculated by subtraction of ATP-only measurements from ATP+ADP values. All measurements were normalized to cell number and are reported as relative intracellular nucleotide abundance per 10^6^ cells.

### NAD^+^ and NADH measurements

Intracellular NAD^+^ and NADH levels were quantified using the NAD/NADH-Glo assay (Promega) according to the manufacturer’s instructions, with modifications for bacterial samples. Cultures were grown anaerobically, subcultured to exponential phase, normalized to an equivalent cell density (OD_600_ = 0.8), and exposed to defined oxygen conditions as described above in an open atmosphere containing defined levels of % O_2_, 5% CO_2_, and 2.5% H_2_ (balance N_2_) at 37 °C. Cultures were maintained in flasks with a large air-liquid interface and continuous shaking (200 rpm) to facilitate efficient gas exchange.

For extraction and differential quantification, normalized cell suspensions (25 μL) were lysed in bicarbonate-containing alkaline buffer supplemented with dodecyltrimethylammonium bromide (DTAB). Lysates were then split and subjected to selective acid or base treatment coupled with heat to exploit the differential stability of NAD^+^ and NADH. For NAD^+^ measurements, samples were treated with 0.4 N HCl and incubated at 60 °C to selectively degrade NADH while preserving NAD^+^. For NADH measurements, samples were maintained under alkaline conditions and heated to 60 °C to selectively degrade NAD^+^ while preserving NADH. Following treatment, samples were neutralized and transferred to fresh plates for detection.

Equal volumes of NAD/NADH-Glo detection reagent were added to each well, and luminescence was recorded after incubation at room temperature. Signals from acid-treated samples correspond to NAD^+^, whereas signals from base-treated samples correspond to NADH. Absolute concentrations were determined using standard curves generated from serial dilutions of NAD standards and normalized to cell number. NAD^+^/NADH ratios were calculated for each condition.

### Pyruvate dehydrogenase (PDH) activity assay

Pyruvate dehydrogenase (PDH) activity was measured using a colorimetric assay (Abcam), with modifications for bacterial samples. *B. theta* strains were grown overnight in BHI under anaerobic conditions, subcultured to exponential phase, normalized to an equivalent cell density (OD_600_ = 0.8), and exposed to defined oxygen conditions as described above in an open atmosphere containing the indicated O_2_ concentrations, 5% CO_2_, and 2.5% H_2_ (balance N_2_) at 37 °C. Cultures were maintained in flasks with a large air–liquid interface and continuous shaking (200 rpm) to facilitate efficient gas exchange.

Cells were harvested by centrifugation, washed twice with PBS, and normalized to OD_600_ = 5. Cell pellets were resuspended in ice-cold PDH assay buffer and permeabilized by brief chloroform treatment (1% v/v, 5 s vortex) to enable substrate access while preserving enzymatic activity.

Permeabilized cells (10 μL) were added to assay wells and brought to a final volume of 50 μL with PDH assay buffer. Reactions were initiated by addition of PDH reaction mix containing assay buffer, substrate, and developer according to the manufacturer’s instructions. Plates were incubated at 37 °C, and absorbance at 450 nm (A_450_) was measured kinetically at defined intervals. PDH activity was quantified using a standard curve generated from serial dilutions of NADH standards prepared in parallel. Reaction rates were calculated from the linear range of signal accumulation prior to saturation and normalized to cell input.

### Whole-cell Hydrogenase Assay

Hydrogenase activity in *B. theta* was measured using a modified methyl viologen reduction assay^123^. Briefly, hydrogenase activity was quantified in whole cells by monitoring the reduction of methyl viologen spectrophotometrically at 600 nm. *B. theta* strains were grown overnight in BHI under anaerobic conditions, subcultured to exponential phase, normalized to an equivalent cell density (OD_600_ = 0.8), and exposed to defined oxygen conditions as described above. H_2_-free atmosphere was used as control.

Following oxygen exposure, cells were harvested by centrifugation, washed twice in assay buffer (50 mM Tris-HCl, pH 7.5, 50 mM NaCl, supplemented with 100 μM MgCl_2_ and 100 μM NiCl_2_), and resuspended to OD_600_ = 0.8. Aliquots (200 μL) of cell suspensions were transferred into 96-well plates under either anaerobic (0% O_2_) or defined oxygen conditions and equilibrated for 10 min prior to measurement.

Hydrogenase reactions were initiated by addition of methyl viologen-containing developing solution (final concentration 20 mM), followed by immediate kinetic measurement of absorbance at 600 nm at 30 s intervals for 5 min. Reduction of methyl viologen, which serves as an artificial electron acceptor, reflects hydrogenase-dependent electron transfer activity.

For each well, reaction rates were determined from the linear slope of absorbance increase over time. Background signals from no-cell controls were subtracted, and activities were normalized to cell density. Relative hydrogenase activity was calculated by normalizing to the mean activity of wild-type controls measured in parallel.

### Untargeted metabolomics

For the LC-MS/MS analysis, all culture supernatant samples were normalized to a total volume of 200 µL per sample. Isotopically labeled standards (phenylalanine and biotin) were spiked into individual samples prior to sample preparation to assess sample process variability. Following normalization, deproteinization was performed by cold methanol precipitation via overnight incubation at –80°C, followed by centrifugation at 4°C for 10 mins. The clear supernatant was then transferred to a new tube and dried down via cold vacuum centrifugation.

Prior to LC–MS/MS analysis, samples were reconstituted in 80µL of mobile phase A containing isotopically labeled internal standards (tryptophan, inosine, valine, and carnitine). Samples were centrifuged at 10K rpm for 5 min to remove insoluble material before analysis.

A pooled quality control (QC) sample was prepared by combining equal volumes (10µL) from each of the 20 individual experimental samples and was used for column conditioning (eight injections), retention time alignment, and evaluation of mass spectrometry instrument reproducibility throughout the analytical batch.

LC-MS and LC-MS/MS analyses were performed as previously described using a high-resolution Q-Exactive HF hybrid quadrupole-Orbitrap mass spectrometer (Thermo Fisher Scientific, Bremen, Germany) equipped with a Vanquish UHPLC binary system (Thermo Fisher Scientific, Bremen, Germany)^124,125^. Briefly, extracts (4 μL injection volume) were separated on an ACQUITY UPLC BEH Amide HILIC 1.7 μm, 2.1 ×□100 mm column (Waters Corporation, Milford, MA) held at 30 °C using the LC method as previously described^126–130^.

### Untargeted metabolomics data processing and statistical analysis

Raw mass spectrometry (MS) data were imported, processed, normalized, and evaluated using Progenesis QI software (version 3.0; Non-linear Dynamics, Newcastle, UK). Relative abundances of spiked, heavy-labeled QA/QC standards showed <17 and ≤20% variability for sample preparation and <6 and <4% for instrument variability for negative and positive ion modes, respectively. All MS and MS/MS runs were aligned to a quality control (QC) pooled reference sample. Detected ions (i.e., paired retention time and m/z features), were de-isotoped and de-adducted to produce distinct compounds. Data were normalized to all compounds, and statistical significance was evaluated using analysis of variance (ANOVA) on log transformed, normalized compound abundance values. Both tentative and putative metabolite annotations (Confidence Level 1-3)^131^ were assigned based on accurate mass measurements (mass error <5 ppm), isotopic pattern similarity, MS/MS fragmentation spectrum, and retention time matching by querying the Human Metabolome Database^132^ and an in-house reference library.

Metaboanalyst 6.0 (www.metaboanalyst.ca/) was used to generate heat maps^133^. The untargeted metabolomics data is available at the NIH Common Fund’s National Metabolomics Data Repository (NMDR) website, the Metabolomics Workbench, https://www.metabolomicsworkbench.org, under the assigned Study IDs ST003919 and ST004857. The data can be accessed directly via its Project DOIs: https://doi.org/10.21228/M8PR93 and http://dx.doi.org/10.21228/M8QC4Q.

### Targeted quantification of mRNA levels in intestinal tissue and contents

Colonic and cecal tissues were homogenized using a bead beater (Bertin Precellys 24 Touch Homogenizer), and total RNA was extracted using TRI reagent (Molecular Research Center) according to the manufacturer’s instructions. Residual genomic DNA was removed using the TURBO DNA-free Kit (Ambion). Complementary DNA (cDNA) was synthesized using the SuperScript VILO cDNA Synthesis Kit (Thermo Fisher Scientific). Quantitative real-time PCR (qRT-PCR) was performed using PowerUp SYBR Green Master Mix (Applied Biosystems), and fluorescence signals were acquired on a CFX Maestro 2.3 system (Bio-Rad). Primers listed in the Key Resources Table were used at a final concentration of 250 nM. Gene expression levels were normalized to *Gapdh* for host transcripts or *gyrB* for bacterial transcripts, and relative expression was calculated using the ΔCt method.

### Hypoxia staining

Tissue hypoxia was assessed using pimonidazole labeling (Hypoxyprobe). *S*. Tm-infected mice receiving the indicated treatments were administered pimonidazole hydrochloride (60 mg/kg) by intraperitoneal injection 30 min prior to euthanasia. Following euthanasia, cecal and colonic tissues were collected as described above and fixed in 10% neutral-buffered formalin, embedded in paraffin, sectioned, and mounted on glass slides.

Paraffin sections were deparaffinized, rehydrated, and subjected to antigen retrieval by incubation with proteinase K (New England Biolabs) in TE buffer (10 mM Tris, 1 mM EDTA, pH 8.0) for 10 min at 37°C. Slides were washed in PBS and blocked for 1 h at 4°C using M.O.M. mouse IgG blocking reagent (Vector Laboratories). Sections were then incubated overnight at 4°C with anti-pimonidazole rabbit primary antibody (Hypoxyprobe; 1:100 dilution). After washing, slides were incubated with Alexa Fluor 488–conjugated goat anti-rabbit secondary antibody (Cell Signaling Technology; 1:150 dilution) for 90 min at room temperature. Nuclei were counterstained with Hoechst for 5 min, followed by PBS washes and mounting.

Tiled fluorescence images (5 × 5 fields with 15% overlap) were acquired using a Nikon ECLIPSE Ti-2 microscope equipped with a 20× objective and Perfect Focus System. Image stitching was performed in real time using NIS-Elements software (version 6.10.01). Regions of interest (ROIs) were manually defined to exclude well-oxygenated lamina propria. Hypoxia signal intensity (pimonidazole staining) was quantified and normalized to the corresponding Hoechst signal within each ROI.

### Chemoproteomic assessment of cysteine reactivity in *B. theta* during oxygen exposure

The wild-type *B. theta* strain was grown overnight in BHI under anaerobic conditions, subcultured to exponential phase, and either maintained anaerobically or exposed to 4% O_2_. Cultures were harvested after 90 minutes of O_2_ exposure (5,000 × g, 10 min, 4□°C). Following oxygen exposure, subsequent protein isolation and cysteine labeling steps were performed in degassed buffers under a nitrogen atmosphere in an anaerobic chamber to preserve redox state. Cells were washed in PBS, lysed by sonication, and clarified by centrifugation (10,000 × g, 10 min, 4□°C). Protein concentrations were normalized to 2 mg mL⁻¹.

Cysteine reactivity was quantified using isotopic activity-based protein profiling (isoTOP-ABPP)^90^. Briefly, anaerobic and oxygen-exposed proteomes (2 mg) were labeled with isotopic alkyne-functionalized iodoacetamide probes^134^ (100 μM; anaerobic – iodoacetamide light (IAL); oxygen-exposed – iodoacetamide heavy (IAH)) for 1 h at room temperature in the dark. Probe-labeled proteins were conjugated to photocleavable (PC) biotin-azide capture reagents via Cu(I)-catalyzed azide-alkyne cycloaddition (CuAAC; 100 µM PC biotin-azide; 1 mM TCEP; 100 µM TBTA; and 2 mM CuSO_4_) for 1 h at room temperature in the dark^135^. Next, light and heavy samples were precipitated (6,500 × g, 10 min, 4□°C), combined, washed with methanol, and resolubilized in SDS-containing PBS buffer. Samples were diluted to 0.2 % SDS, enriched on streptavidin resin, and extensively washed to remove non-specific interactions. On-bead reduction (10 mM DTT, 20 min, 65 °C) and alkylation (20 mM iodoacetamide, 30 min, 37 °C) were performed prior to overnight trypsin digestion (1 µg trypsin; 2 M urea in PBS, 1 mM CaCl_2_, overnight, 37 °C). Probe-labeled peptides were released by photocleavage of the linker, acidified, and desalted for LC–MS/MS analysis.

For complementary quantification of protein abundance, whole-proteome samples were analyzed by reductive dimethylation (ReDiMe)^136^. Proteins were precipitated (5% TCA), acetone washed, and resuspended in urea buffer (8 M urea in 100 mM TEAB), followed by reduction (15 mM DTT, 15 min, 75 °C), alkylation (12.5 mM iodoacetamide, 30 min, 37 °C), and tryptic digestion (1 µg trypsin; 1 M urea in 100 mM TEAB, 1 mM CaCl_2_, overnight, 37 °C). Peptides from anaerobic and oxygen-exposed samples were isotopically labeled with either light (oxygen-exposed) or heavy (anaerobic) formaldehyde (1% H_3_^12^CO or D_2_H^13^CO, 50 mM sodium cyanoborohydride, 2 h, room temperature), quenched with 1% ammonium hydroxide, combined, desalted, and fractionated by off-line high-pH reversed-phase chromatography^137^ prior to LC-MS/MS.

Peptides were analyzed by nanoLC–MS/MS on Orbitrap Exploris 240 mass spectrometer coupled to a Dionex Ultimate 3000 RSLCnano system in data-dependent acquisition (DDA) mode. MS1 scans (400-1800 MW, 120,000 resolution, RF lens 65%, AGC target 300%, automatic maximum injection time) were acquired every two seconds with dynamic exclusion enabled (repeat count 2, duration 10 s), followed by a variable number of data-dependent MS2 fragmentation scans (15,000 resolution, AGC 75%, maximum injection time 100 ms) of the most abundant precursor ions. MS2 analysis consisted of the isolation of precursor ions (isolation window 2 m/z), filtered for monoisotopic peak determination, theoretical isotopic envelope fit, intensity (5E4), and charge state (+2 – +6), followed by higher-energy collision dissociation (HCD, collision energy 30%). Spectra were searched using Thermo Proteome Discoverer V2.4 software package against a *B. thetaiotaomicron* UniProt database (UP000001414) using SequestHT and Percolator algorithms with trypsin specified as the protease (max of 2 missed cleavages) and precursor mass tolerance set to 10 ppm with a fragment mass tolerance of 0.02 Da. In addition to dynamic modifications representing oxidation of methionine (+15.995), as well as acetylation (+42.011) and/or methionine-loss (+131.040) of the protein N-terminus, appropriate static and variable modifications were set for cysteine alkylation and isotopic labeling. For ReDiMe analysis, static modifications on lysine and the peptide N-terminus of +28.0313 (light) or +34.0632 (heavy) were specified. For isoTOP-ABPP, dynamic modifications on cysteine of +57.021 (alkylation) and either +285.159 (IAL) or +291.179 (IAH) were specified. Peptide and protein identifications were filtered to a false discovery rate ≤1%.

Relative cysteine reactivity and protein abundance were quantified from peptide light/heavy (ReDiMe) or heavy/light (isoTOP-ABPP) isotopic ratios using established pipelines (Proteome Discoverer)^90,138^. Protein-level ratios were calculated as the median of corresponding peptide measurements. All experiments were performed as two technical replicates of independent biological replicates (n = 3-5). Functional annotation of cysteine residues and protein localization was obtained from UniProt and integrated with quantitative datasets to identify oxygen-sensitive cysteine sites across the proteome. The mass spectrometry proteomics data have been deposited to the ProteomeXchange Consortium via the PRIDE^139^ partner repository with the dataset identifier PXD079134. Data are also available via ProteomeXchange^140^ with identifier PXD079134 (token: rzvuz8rH7YoT).

### TCA Cycle metabolite quantification

For targeted metabolite quantification, *B. theta* strains were grown overnight in SDM under anaerobic conditions, subcultured to exponential phase, and exposed to defined oxygen conditions as described above. Cultures were harvested after 1 hour of O_2_ exposure by centrifugation (5,000 × g, 10 min, 4□°C). The culture supernatant was filtered through a 0.22 μm filter, 10 μL of internal standard solution (250 μM of each: Succinate-d_4_, Pyruvate-^13^C_3_, Lactate-^13^C_3_) was added, and the cell-free supernatant was dried via vacufuge at ambient temperature, and reconstituted in 100 μL NaPO_4_ Buffer (250 mM Na_2_HPO_4_, 250 mM NaH_2_PO_4_ (pH ∼4) in acetonitrile:water (1:1)) for subsequent extracellular metabolite quantification.

Intracellular metabolites were collected in parallel by resuspending the cell pellet in 1 mL of ice-cold extraction solution (acetonitrile:methanol:water (2:2:1), vortexing, freezing at –20 for 1 hr. 10 μL of internal standard solution (250 μM of each: Succinate-d_4_, Pyruvate-^13^C_3_, Lactate-^13^C_3_. and cell extract was dried via vacufuge at ambient temperature, and reconstituted in 100 μL NaPO_4_ Buffer (250 mM Na_2_HPO_4_, 250 mM NaH_2_PO_4_ (pH ∼4) in acetonitrile:water (1:1)) was then added. Cell extract was reserved for normalization via protein quantification using a BCA assay (Pierce™ BCA Protein Assay, Thermo).

For stable isotope tracing of tricarboxylic acid cycle metabolites (pyruvate, succinate, malate, fumarate, α-ketoglutarate, citrate, and lactate), *B. theta* strains were grown overnight in *Bacteroides* Minimal Media (BMM) under anaerobic conditions, subcultured to exponential phase, and exposed to defined oxygen conditions as described above. 1 mL of oxygen exposed culture was filtered through 0.45 μm filter membranes (HVLP, Millipore) prewarmed with medium, and filtered bacteria were washed with 2 mL medium. Filters were rapidly transferred to 1 mL of extraction solution (acetonitrile:methanol:water (2:2:1)) and 10 μL of internal standard solution was added.

LC-MS Analysis: For measurement of tricarboxylic acid cycle metabolites and their respective isotopomers (pyruvate, succinate, malate, fumarate, α-ketoglutarate, citrate, and lactate), metabolites were derivatized to their corresponding dansyl hydrazides to improve their ionization efficiency and chromatographic resolution. Briefly, 50 μL of NaPO_4_ Buffer, 10 μL H_2_O, 12.5 μL 150 mg/mL EDC (1-Ethyl-3-(3-dimethylaminopropyl)carbodiimide) in H_2_O, 12.5 μL 50 mg/mL dansyl hydrazine in acetonitrile, and 25 μL of internal standard spiked and reconstituted cell free supernatant or clarified cell extract were combined. After shaking overnight at 22 °C, reactions were quenched with 12.5 μL x 5 % (v/v) TFA in water. Quenched reactions were centrifuged at 18,000 x g, then transferred to 2 mL autosampler vials equipped with low-volume polypropylene inserts and Teflon-lined rubber septa. The sample injection volume was 10 μL. TCA metabolite calibration standards were prepared in water and derivatized in the same manner. Liquid chromatography-high resolution mass spectrometry (LC-HRMS) analysis was performed using a Thermo Q Exactive HF hybrid quadrupole/orbitrap high resolution mass spectrometer interfaced to a Vanquish Horizon HPLC system (Thermo Fisher). High resolution mass spectra were acquired in positive ion mode over a precursor ion scan range of m/z 300 to 800 at a resolving power of 60,000 using the following HESI source parameters: spray voltage 4 kV; capillary temperature 300°C; HESI temperature 100 °C; Ss-lens 95; N_2_ sheath gas 40; N_2_ auxiliary gas 10. Extracted ion chromatograms were constructed for each TCA cycle metabolite based on the following [M+H]^+^ exact masses and a mass tolerance of +/− 5 ppm. Pyruvate: M, 583.1792; M+1, 584.1825; M+2, 585.1859; M+3, 586.1893; M+4, 587.1926. Succinate: M, 366.1118; M+1, 367.1152; M+2, 368.1185; M+3, 369.1219; M+4, 370.1252. Malate: M, 382.1067; M+1, 383.1101; M+2, 384.1134; M+3, 385.1168; M+4, 386.1202. Fumarate: M, 364.0962; M+1, 365.0995; M+2, 366.1029; M+3, 367.1062; M+4, 368.1096. Alpha-ketoglutarate: M, 641.1847; M+1, 642.1880; M+2, 643.1914; M+3, 644.1947; M+4, 645.1981.

Data acquisition and quantitative spectral analysis were done using Thermo Xcalibur version 4.1.31.9 and Thermo LCQuan version 2.7, respectively. Calibration curves were constructed by plotting peak areas against analyte concentrations for a series of eleven calibration standards, ranging from 0.15 to 1500 total pmol of each BA. A weighting factor of 1/C2 was applied in the linear least-squares regression analysis to maintain homogeneity of variance across the concentration range. An Acquity HSS C18 reverse phase analytical column (2.1 x 150 mm, 1.7 μm, Waters, Milford, MA) was used for all chromatographic separations. Mobile phases were made up of 0.2 % formic acid + 10 mM ammonium formate in (A) water/methanol/acetonitrile (8:1:1) and in (B) methanol/acetonitrile (4:1). Gradient conditions were as follows: 0-1.0 min, B = 10 %; 1-15 min, B = 10-100 %; 15-17.5 min, B = 100 %; 17.5-18 min, B = 100-10 %; 18-22 min, B = 10 %. The flow rate was maintained at 300 μL/min, and the total chromatographic run time was 22 min. A software-controlled divert valve was used to transfer the LC eluent from 0 to 5 min of each chromatographic cycle to waste.

### Targeted quantification of citrulline

Because carbamoyl phosphate is chemically unstable and undergoes rapid non-enzymatic hydrolysis during metabolite extraction and processing^97,98^, carbamoyl phosphate-dependent metabolic activity was assessed indirectly through citrulline production^99^. Mid-logarithmic phase *B. thetaiotaomicron* cultures were grown anaerobically to OD_600_ ≈ 0.8, exposed to either anaerobic conditions or 10% O_2_ for 1 h, and supplemented with NaHCO_3_ (10 mM final concentration) and L-glutamine (2 mM final concentration) to support carbamoyl phosphate synthesis. Cultures were then incubated for an additional 1 h under the same atmospheric conditions prior to harvest. Cells were rapidly chilled, pelleted by centrifugation, and lysates were prepared in methanol. For citrulline quantification, 100 μl of lysate was evaporated to dryness under nitrogen gas and reconstituted in 75 μl acetonitrile/water (1:1). Samples were then mixed with 20 μl of 0.1 M NaHCO3 and 20 μl of dansyl chloride solution (10 mg/ml in acetonitrile), incubated for 20 min at room temperature, and quenched with 20 μl of 5% acetic acid in water. Samples were clarified prior to LC-MS/MS analysis. Calibration standards were prepared by adding 10 μl of citrulline standards to matrix without internal standard and processed in parallel with experimental samples. Citrulline abundance was quantified from the calibration curve and normalized to cell number.

### Bulk RNAseq and analysis

For bacterial RNA-seq, *B. theta* strains were cultured anaerobically in BHIS and subcultured into lactate-supplemented BHIS under either anaerobic conditions or in the presence of 0.1% O₂ for 3 h at 37°C. RNAprotect Bacteria Reagent (Qiagen) was added immediately prior to harvest to stabilize transcripts. Cells were lysed using Lysing Matrix B (MP Biomedicals) in TRI reagent (Molecular Research Center), and total RNA was purified using the RNeasy Mini Kit (Qiagen) according to the manufacturer’s instructions. Strand-specific, paired-end (150 bp) libraries were generated using an RNAtag-seq strategy^141^ and sequenced on an Illumina NovaSeq platform.

Sequencing reads were trimmed using the BBMap software suite and aligned to the *B. theta* VPI-5482 reference genome using Bowtie2^141^. Gene-level counts were obtained using featureCounts^142^ and differential expression analysis was performed using DESeq2^143^. Raw sequencing data have been deposited in the European Nucleotide Archive under accession number PRJEB112738 (secondary accession ERP193206).

### Genome-Scale Metabolic Modeling

A genome-scale metabolic model (GEM) for *B. theta* VPI-5482 was reconstructed following established protocols^130^ using the COBRA Toolbox (v2.13.3) in MATLAB R2022b R^131^. An existing curated GEM for *B. theta* (doi.org/10.1016/j.ymben.2021.10.005) served as the primary template, supplemented by additional Gram-negative reference models from the BiGG database^132^, including multiple *E. coli* GEMs^133, 134^, *Shigella boydii*^134^, *Shigella dysenteriae*^134^, *Shigella flexneri*^134^, and *Klebsiella pneumoniae*^135^. Homologous metabolic genes were identified by BLAST (E-value < 1 × 10⁻³□, sequence identity > 50%), and corresponding reactions were incorporated into a draft reconstruction followed by manual curation, removing unsupported gene-reaction relationships. The model was curated to ensure growth under defined media conditions known to support *B. theta*, appropriate ATP yields, and mass and charge balance, consistent with prior reconstruction standards and recapitulating all validations performed in the original publication. Gene-protein-reaction associations were refined based on genome annotation and literature, and gap-filling was performed where necessary to support physiologically relevant growth. All additional reactions added through gap filling were iteratively curated and retained or rejected.

To generate context-specific models, transcriptomic data were integrated using the StanDep algorithm^53^, to determine condition-dependent reaction activity across distinct niches. To approximate the diverse, undefined nutrient environment of BHI, all exchange reactions were constrained to low uptake rates (≤0.5 mmol gDW⁻¹ h⁻¹), tuned to match the observed growth rate of *B. theta* in BHI culture. Resulting condition-specific models were optimized for biomass production using parsimonious flux balance analysis (pFBA)^136^, yielding minimal flux sets consistent with the observed expression profiles and without thermodynamically infeasible loops. Flux through each metabolite (**Supplementary Fig. S2**) was computed as the sum of reaction rates producing that compound scaled by the stoichiometric coefficient. All fluxes were normalized by condition-specific growth rate to make the values comparable.

### Phylogenetic analysis of genes involved in oxidative central metabolism

The accessions and metabolic traits for the genomes we analyzed were taken from the members of the class *Bacteroidia* described in Dataset S01 from Liao et al.^144^ The facultative outgroup *E. coli* Nissle 1917 (GCA_021559835.1) was also included. Genome assemblies were acquired from NCBI with the NCBI Datasets v18.19.0 tool^145^ and annotated with Prokka v.1.14.5^146^ with the compliant option. Orthology and species tree inference were performed with Orthofinder v3.1.3.post1.dev1^147–150^ with MAFFT v7.526^151^ set as the aligner. Paralogs of *carA*, *carB*, and *oxe* were identified and removed based on gene tree topologies. To construct these gene trees, protein sequences from each gene’s respective orthogroup were aligned with MAFFT, trimmed with ClipKIT v 2.11.4^152^, and then input to IQ-TREE v3.1.0^153^ for phylogenetic inference. This involved selecting the best model (including profile mixture models) with ModelFinder^154^ and calculating 1000 ultrafast bootstrap^155^ samples. Alignments and IQ-TREE gene trees were imported into R for visualization with ggtree v4.1.2^152^. Before visualization, further filtering was performed to remove remaining duplicate gene sequences.

### Animal Experiments

All animal experiments were conducted in accordance with protocols approved by the Institutional Animal Care and Use Committee at Vanderbilt University Medical Center. C57BL/6J wild-type mice (Jackson Laboratory, stock no. 000664) were maintained under specific pathogen–free conditions in sterile cages on a 12 h light/dark cycle with ad libitum access to food and sterile water.

Seven– to nine-week-old male and female mice were semi-randomly assigned to treatment groups prior to experimentation. To facilitate colonization, mice were administered an antibiotic cocktail consisting of ampicillin, metronidazole, vancomycin, and neomycin (5 mg of each antibiotic per mouse) by oral gavage daily for 2 days (competitive colonization) or 5 days (single colonization), as previously described^156^.

For single-colonization experiments, mice were orally inoculated on day 5 with 1 × 10^9^ CFU of the indicated *B. theta* strain or left uninfected as controls. For competitive colonization experiments, mice were inoculated with an equal mixture containing 0.5 × 10^9^ CFU of each indicated strain and its corresponding mutant. Twenty-four hours later, mice were inoculated intragastrically with 1 × 10^9^ CFU of *Salmonella enterica serovar* Typhimurium SL1344 or LB-mock control.

At the experimental endpoint, mice were euthanized and cecal and colonic tissues were collected, flash-frozen in liquid nitrogen for downstream mRNA analysis or fixed in 10% formalin for histopathological analysis. For culture-based quantification, cecal and colonic contents were collected in sterile PBS, serially diluted, and plated on selective media to determine bacterial burdens of *B. theta* and *S*. Tm.

### Quantification and Statistical analysis

Unless noted otherwise, data analysis was performed in GraphPad Prism v11.0.0. Values of bacterial population sizes, competitive indices, fold changes in mRNA levels, and normalized colon length were normally distributed after transformation by the natural logarithm. A two-tailed Student’s *t*-test was used for ln-transformed data. Unless otherwise stated, *, *P* < 0.05; **, *P* < 0.01; ***, *P* < 0.001; ns, not statistically significant. In all mouse experiments, *N* refers to the number of animals from which samples were taken. Sample sizes (i.e. the number of animals per group) were not estimated *a priori* since effect sizes in our system cannot be predicted. No predicted statistical outliers were removed since the presence or absence of these potential statistical outliers did not affect the overall interpretation. Mice that were euthanized early due to health concerns were excluded from analysis.

## Supporting information

Key Resource Table

## Acknowledgements

We thank Dr. Lynn Bry (Harvard University) for insightful discussions and comments on the conceptual framework of this work. Work in W. Z.’s lab was funded by the NIH (1R35GM147470, 1R01DK134692, R21AI187749, R21DK146015), The Pew Charitable Trust (2023-A-26048), V Foundation (V2022-032), Colorectal Cancer Alliance (10065978), and The G. Harold & Leila Y. Mathers Charitable Foundation (MF-2207-03128). Work in E. W.’s lab was funded by the NIH (R35GM134964). Work in M. G. B’s lab was funded by NIH (R35GM150625). Work in J. A. I.’s lab was funded by the NIH (GM158047). Untargeted metabolomics work was supported in part using the resources of the Center for Innovative Technology (CIT) at Vanderbilt University. This work is supported by Metabolomics Workbench/National Metabolomics Data Repository (NMDR) by NIH grants U2C-DK119886 and OT2-OD030544. Any opinions, findings, conclusions, or recommendations expressed in this material are those of the author(s) and do not necessarily reflect the views of the funding agencies. The funders had no role in study design, data collection, and interpretation, or the decision to submit the work for publication.

## Author contributions

A. E. R., J. A. I., and W. Z. designed the study. A. E. R., M. S., R. T. F. and S. K. R. performed and analyzed all the *in vitro* experiments. A. E. R., M. L-B, L. S. performed and analyzed all the *in vivo* experiments. D. W. B and E. W. performed and analyzed the chemoproteomics profiling. N. M. and K. Z. performed genome-scale metabolic flux integration. A. E. R. and M. W. C. performed targeted quantification of metabolites. A. E. R., A. C. S-R., S. G. C., J. A. M., and S. D. S. performed untargeted metabolomics analyses and analyzed the data. O. F. H. and M. G. B. performed phylogenetic analysis of metabolic genes. W. Z. wrote the manuscript, and all authors commented on the manuscript.

