## Supplemental Figures for "Rational engineering of facultative anaerobiosis enables commensal survival in the oxygenated gut"

### Supplemental Figure. S1

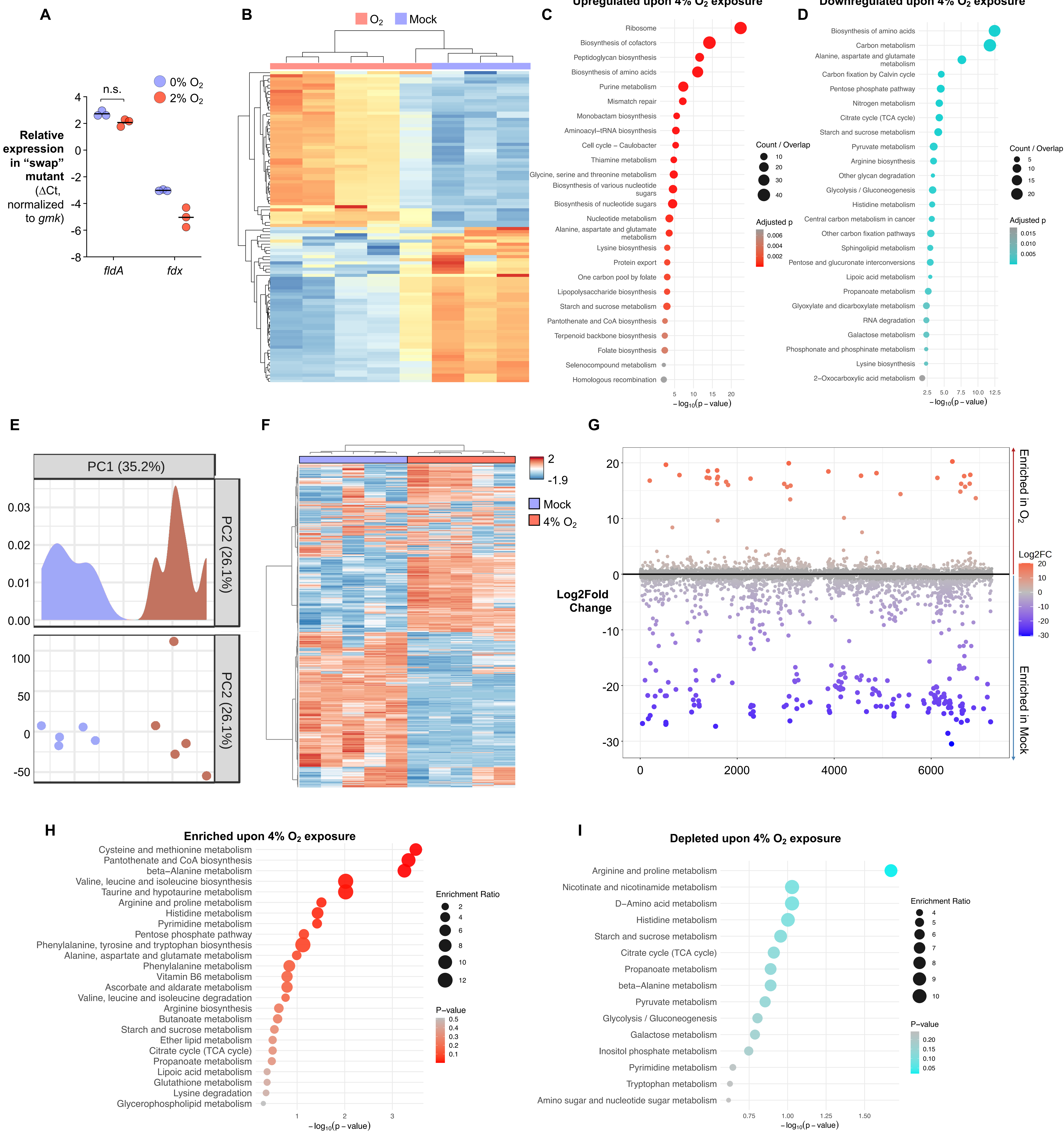

**Supplemental Fig. 1. Oxygen impedes central metabolism in *B. theta* (related to Fig. 1&2).** (**A**) *B. theta* wild-type cells or an isogenic ferredoxin-to-flavodoxin swap mutant (Fd→FldA) were cultured under anaerobic conditions or exposed to 2% $O_2$. (**A**) Expression of *fldA* was quantified by RT-qPCR. (**B-E**) Anaerobically growing wild-type *B. theta* cells were exposed to 4% $O_2$ for 30 min at 37 °C, after which global transcriptional responses were profiled by RNA-seq. (**B**) Heatmap depicting oxygen-responsive gene expression changes. Pathway enrichment analysis of (**C**) upregulated and (**D**) downregulated transcripts. (**E-I**) Anaerobically growing wild-type *B. theta* cells were either maintained under anaerobic conditions or exposed to 4% $O_2$ for 30 min, followed by untargeted analysis of extracellular metabolites by UHPLC-MS/MS. (**E**) Principal component analysis, (**F**) heatmap and (**G**) ranked dot plot showing differentially abundant metabolites during oxygen exposure. Pathway enrichment analysis of (**H**) metabolites enriched and (**I**) metabolites depleted upon oxygen exposure. Bars represent geometric means. *. $P < 0.05$; **, $P < 0.01$. ***, $P < 0.001$.

#### Supplemental Figure. S2

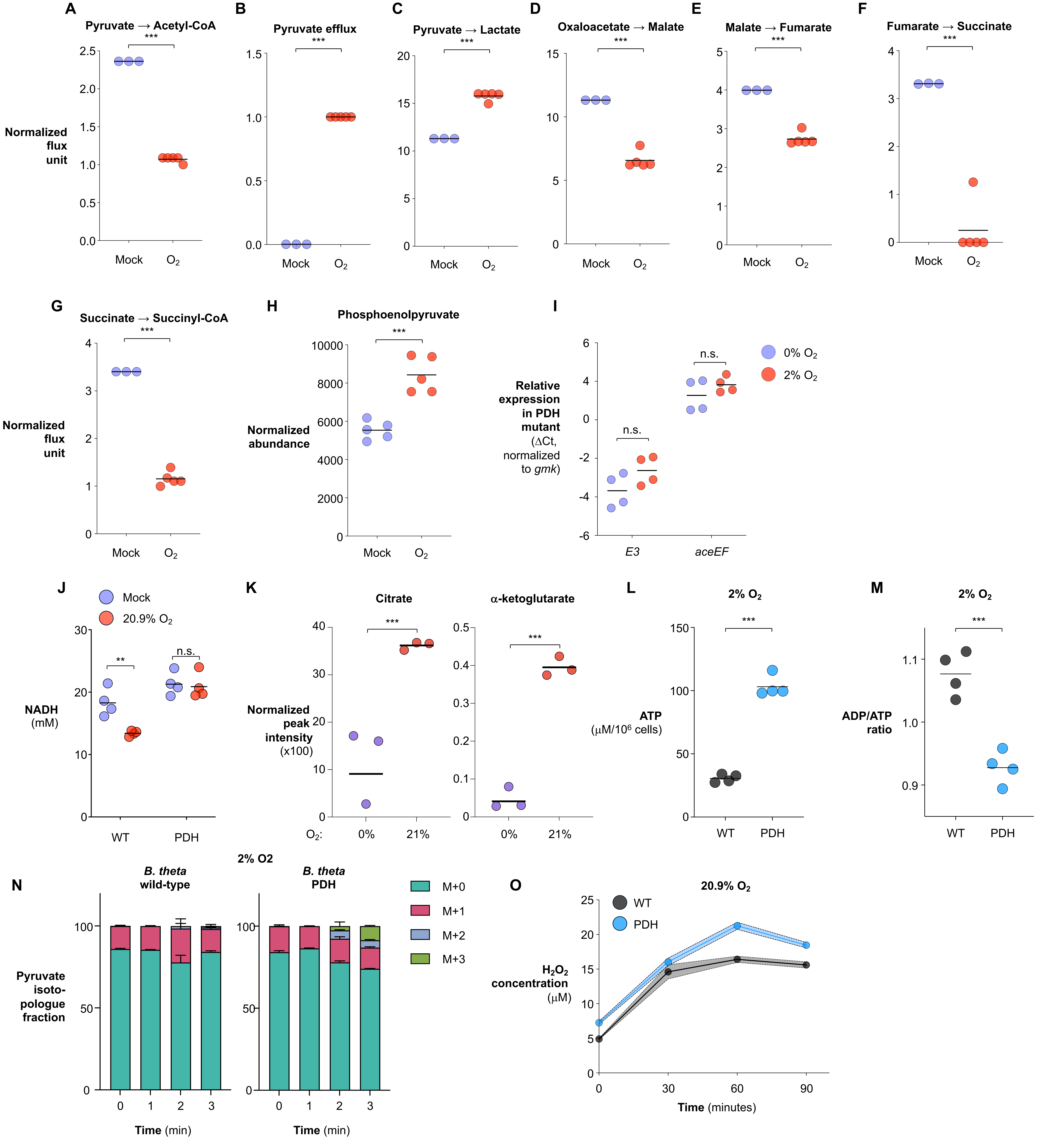

**Supplemental Fig. 2. Oxygen impedes central metabolism in *B. theta* (related to Fig. 2).** (**A-H**) Anaerobically growing wild-type *B. theta* cells were exposed to 4% $O_2$ for 30 min at 37 °C, after which global transcriptional responses were profiled by RNA-seq. Transcriptomic data were integrated into a draft genome-scale metabolic model (GEM) to generate a context-specific metabolic model. (**A-H**) Predicted relative fluxes through the indicated metabolic nodes under anaerobic and oxygen-exposed conditions. (**I**) Wild-type *B. theta* cells or an isogenic pyruvate dehydrogenase (PDH)-expressing mutant were cultured under anaerobic conditions or exposed to 2% $O_2$. *pdh* (*E3* and *aceEF*) expression was quantified by RT-qPCR. (**J-M**) Wild-type *B. theta* cells or an isogenic pyruvate dehydrogenase (PDH)-expressing mutant were cultured under anaerobic conditions or exposed to room air (20.9% $O_2$). Intracellular levels of (**J**) NADH, (**K**) citrate and α-ketoglutarate, (**L**) ATP, and (**M**) ADP/ATP ratio were measured as indicated. (**N**) Anaerobically growing cultures of the indicated *B. theta* strains were exposed to 2% $O_2$ for the indicated time and subsequently incubated with [U-$^{13}$C]glucose. Label incorporation into metabolic intermediates was analyzed by high-resolution LC-MS/MS. Shown are fractional abundance of $^{13}$C-labeled isotopologues following the shift from unlabeled glucose to [U-$^{13}$C]glucose. [M+X] denotes a metabolite isotopologue containing X $^{13}$C atoms (maximum X equals the number of carbons in the molecule). (**O**) Wild-type *B. theta* cells or an isogenic pyruvate dehydrogenase (PDH)-expressing mutant were cultured under anaerobic conditions or exposed to room air (20.9% $O_2$). $H_2O_2$ were measured as indicated. Bars represent geometric means. *. $P < 0.05$; **, $P < 0.01$. ***, $P < 0.001$.

Supplemental Figure. S3

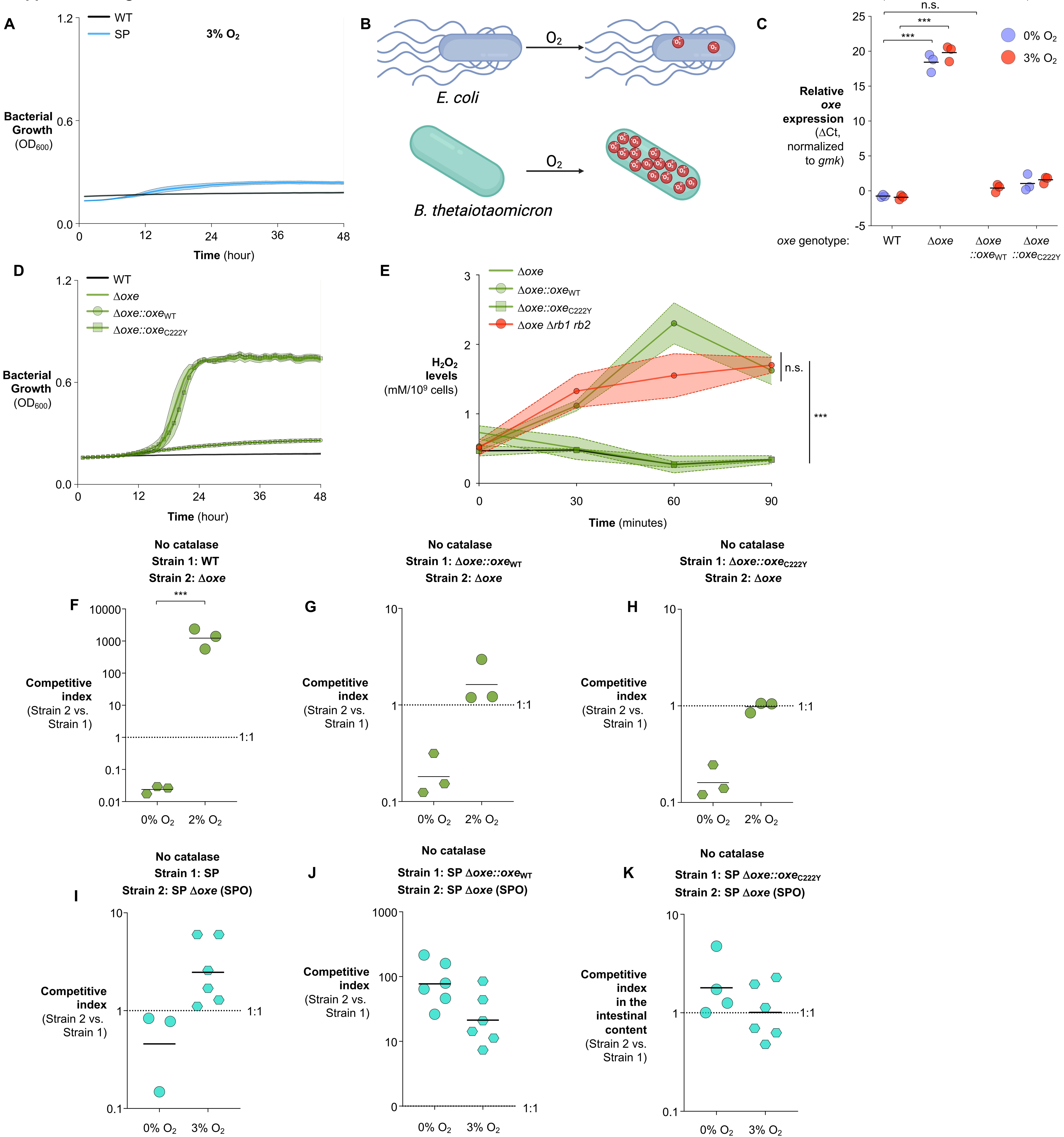

**Fig. S3. Oxe generates intracellular reactive oxygen species that limit microaerobic growth in *B. theta* (related to Fig. 3).** (**A**) Growth of wild-type (WT) and SP strains under 3% $O_2$ with constant shaking, monitored by $OD_{600}$ over time. (**B**) Conceptual model illustrating differential oxygen handling in *E. coli* and *B. theta*, highlighting intracellular ROS accumulation in *B. theta* during oxygen exposure. (**C**) Relative expression of PDH transcripts in the indicated *oxe* backgrounds under anaerobic or 3% $O_2$ conditions. (**D**) Growth of WT, Δ*oxe*, Δ*oxe*::$oxe_{WT}$, and Δ*oxe*::$oxe_{C222Y}$ strains under 3% $O_2$, monitored by $OD_{600}$. (**E**) Extracellular $H_2O_2$ accumulation following oxygen exposure in the indicated strains normalized to cell number. (**F-H**) Pairwise competition assays performed under 0% or 2% $O_2$ without catalase (initial inoculum $10^5$ CFU/ml). (**F**) WT versus Δ*oxe*. (**G**) Δ*oxe*::*oxe*WT versus Δ*oxe*. (**H**) Δ*oxe*::$oxe_{C222Y}$ versus Δ*oxe*. (**I-K**) Pairwise competition assays in the SP background performed under 0% or 3% $O_2$ without catalase. (**I**) SP versus SPO. (**J**) SP Δ*oxe*::$oxe_{WT}$ versus SPO. (**K**) SP Δ*oxe*::$oxe_{C222Y}$ versus SPO. Competitive indices were determined by selective plating. Bars represent geometric means. Dotted lines indicate a competitive index of 1 (no fitness difference). n.s., not significant; ***, $P < 0.001$.

**Supplemental Figure. S4**

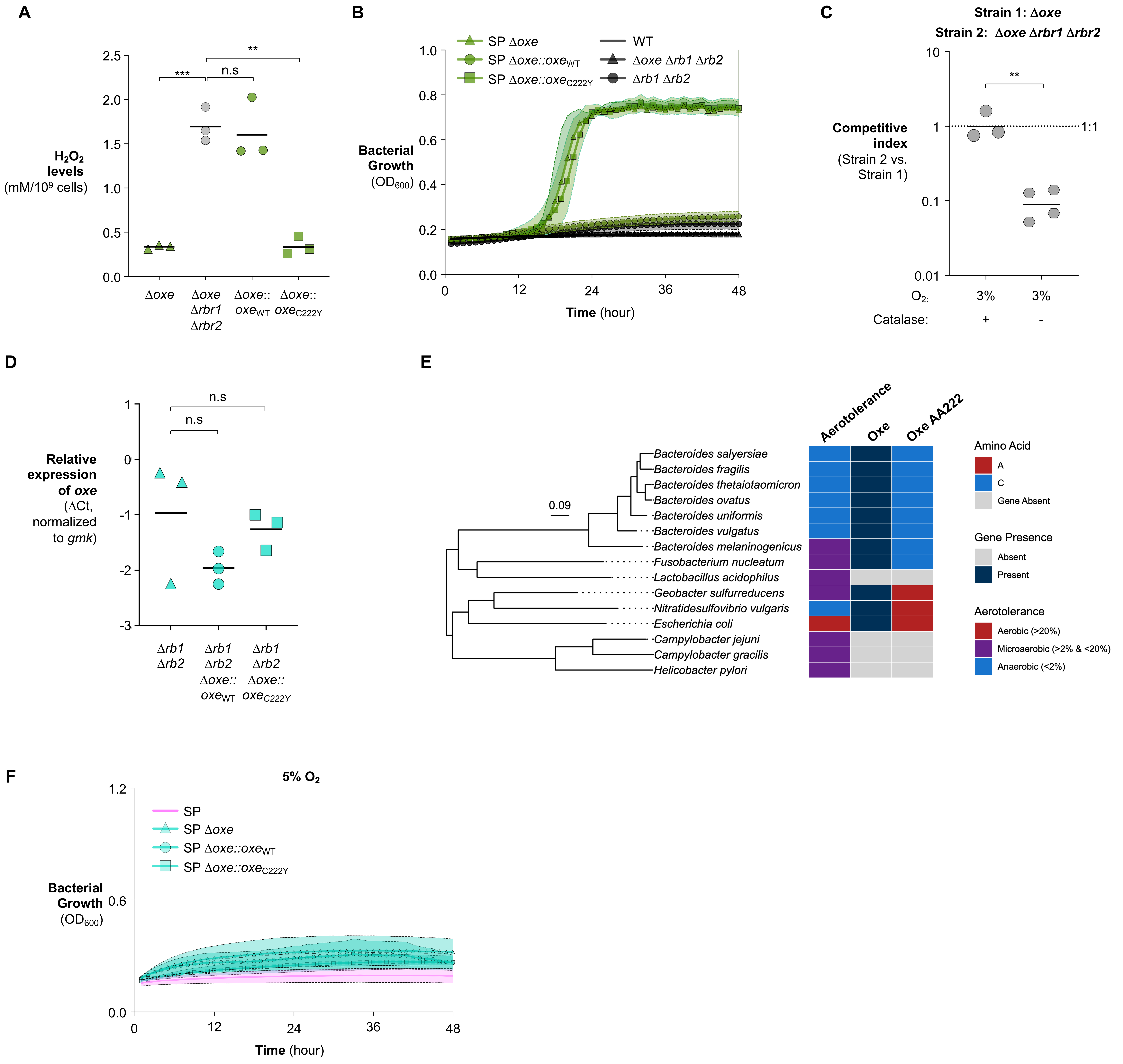

**Fig. S4. Oxe generates intracellular reactive oxygen species that limit microaerobic growth in *B. theta* (related to Fig. 3).** (**A**) $H_2O_2$ accumulation in strains expressing wild-type or C222Y *oxe* alleles in rubrerythrin-proficient or rubrerythrin-deficient backgrounds. (**B**) Growth of WT, Δ*oxe*, and Δ*oxe* Δ*rbr1* Δ*rbr2* strains under 3% $O_2$ with constant shaking, monitored by $OD_{600}$ over time. (**C**) Pairwise competition assays (Δ*oxe* versus Δ*oxe* Δ*rbr1* Δ*rbr2*) examining genetic interactions between *oxe* and rubrerythrins under 3% $O_2$ in the presence or absence of catalase. Competitive indices were determined by selective plating. (**D**) Relative *oxe* transcript levels in the indicated complemented strains measured by RT-qPCR. (**E**) Midpoint-rooted species tree generated using OrthoFinder and annotated with species aerotolerance, *oxe* carriage, and amino acid identity at the focal residue. Scale bar indicates substitutions per site. (**F**) Growth of SP, SPO, and SPO strains complemented with wild-type *oxe* or the $oxe_{C222Y}$ allele under 5% $O_2$ with constant shaking, monitored by $OD_{600}$ over time. Bars represent geometric means. Shaded regions indicate SEM where applicable. *, $P < 0.05$; **, $P < 0.01$; ***, $P < 0.001$.

#### Supplemental Figure. S5

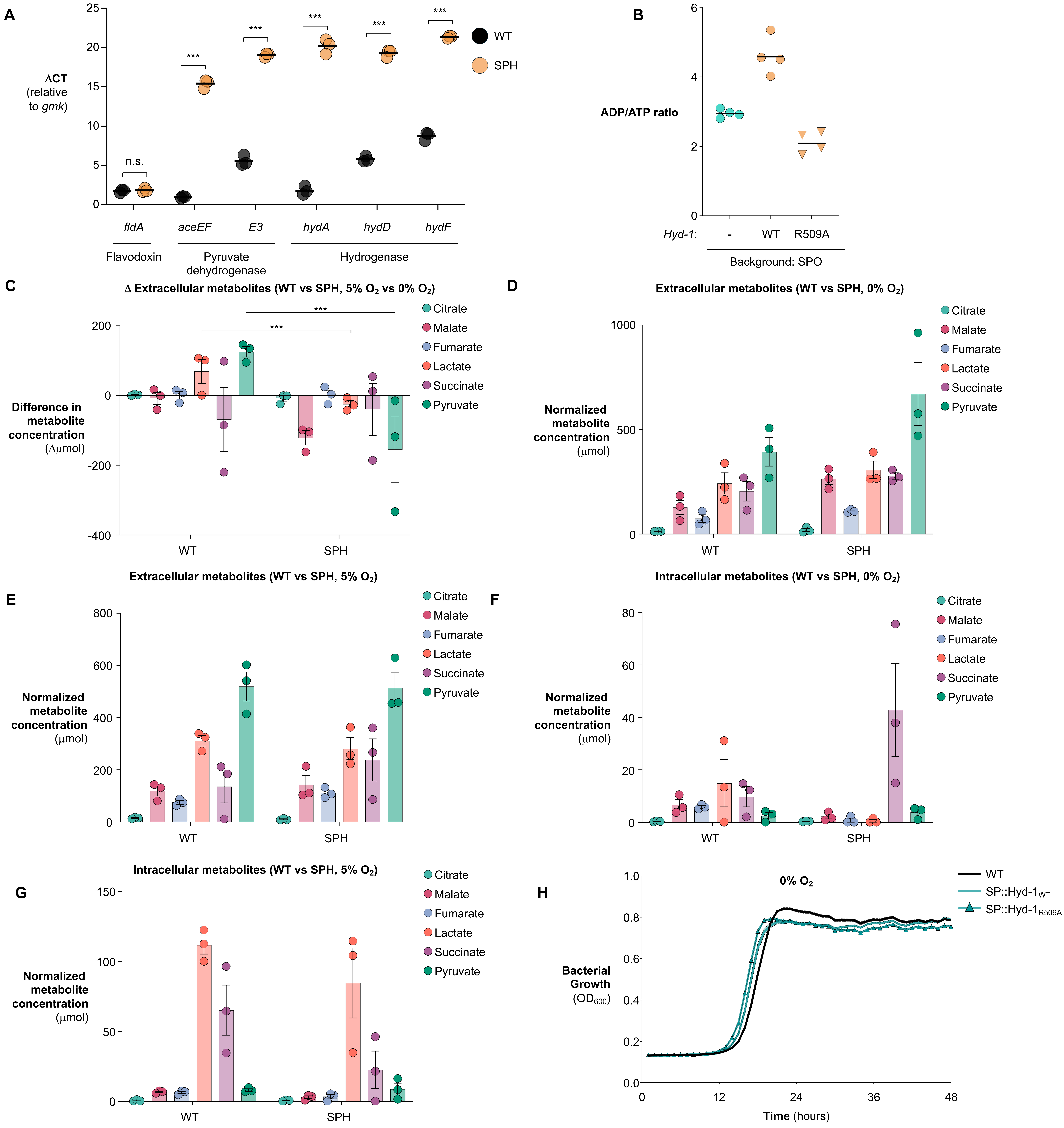

**Supplementary Fig. 5. An oxygen-resistant hydrogenase preserves redox balance and improves *B. theta* fitness under oxic conditions. (related to Fig. 4)** (**A**) *B. theta* wild-type or SPH strains were cultured under anaerobic conditions, and transcript levels of the indicated genes were quantified by RT-qPCR. (**B**) ADP/ATP ratios in SPO strains expressing wild-type Hyd-1 or the catalytically inactive Hyd-1 R509A mutant cultured under 5% $O_2$ in the absence of $H_2$. (**C**) Changes in extracellular metabolite abundance following exposure to 5% $O_2$ relative to anaerobic controls in WT and SPH strains. (**D,E**) Extracellular metabolite abundance in WT and SPH strains cultured under anaerobic conditions (**D**) or exposed to 5% $O_2$ for 30 min (**E**). (**F,G**) Intracellular metabolite abundance in WT and SPH strains cultured under anaerobic conditions (**F**) or exposed to 5% $O_2$ for 30 min (**G**). Metabolites were quantified by LC-MS/MS. (**H**) Wild-type *B. theta* or isogenic derivatives expressing either wild-type Hyd-1 or the catalytically inactive Hyd-1 mutant (R509A) were grown anaerobically, and bacterial growth was monitored by $OD_{600}$. Bars represent geometric means. *, $P < 0.05$; **, $P < 0.01$; ***, $P < 0.001$.

**Supplemental Figure. S6**

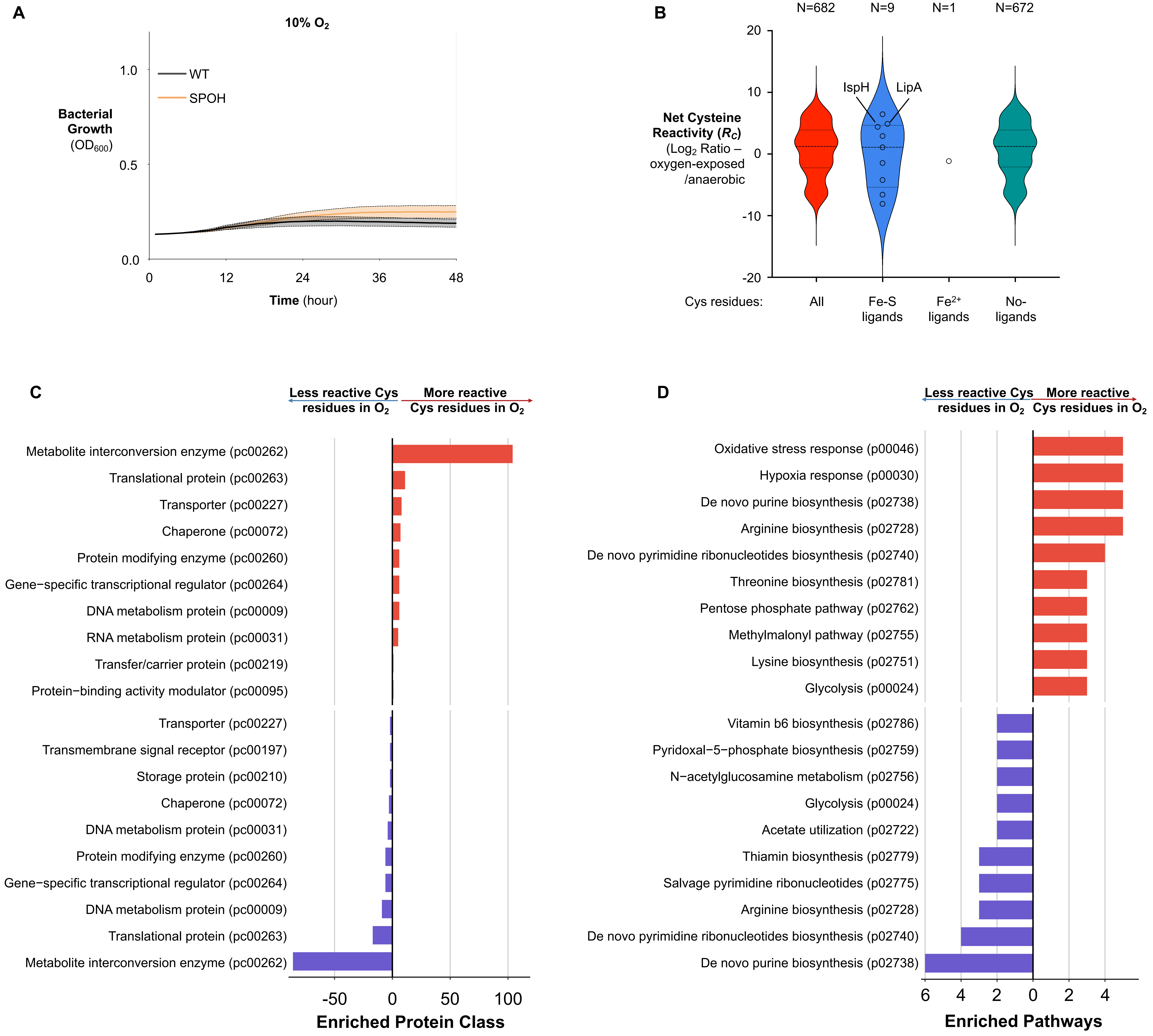

**Supplementary Fig. S6. Oxygen impairs carbamoyl phosphate biosynthesis through cysteine modification in *B. theta* (related to Fig 5).** (**A**) Growth of the indicated *B. theta* strains was monitored by $OD_{600}$ in an open atmosphere containing 10% oxygen concentrations with constant shaking. (**B-D**) Anaerobically growing cultures of wild-type *B. theta* were either maintained under anaerobic conditions or exposed to 4% $O_2$ for 3 h. Cell lysates were labeled with isotopic iodoacetamide-alkyne (IA-alkyne) probes (mock: light; $O_2$-exposed: heavy) to quantify proteome-wide cysteine reactivity by mass spectrometry. After normalization to protein abundance, cysteine $\log_2$(L/H) ratios ($R_C$) report relative changes in labeling, with $R_C > 1$ indicating increased cysteine reactivity upon oxygen exposure. Only cysteine residues with $|\log_2(R_C)| > 1$ are shown. (**B**) Distribution of differentially reactive cysteines across indicated protein functional categories. (**C**) Metabolic pathway and (**D**) molecular function enrichment analyses of proteins harboring cysteines with altered reactivity under oxygen exposure. Bars represent geometric means. *, $P < 0.05$; **, $P < 0.01$; ***, $P < 0.001$.

Supplemental Figure. S7

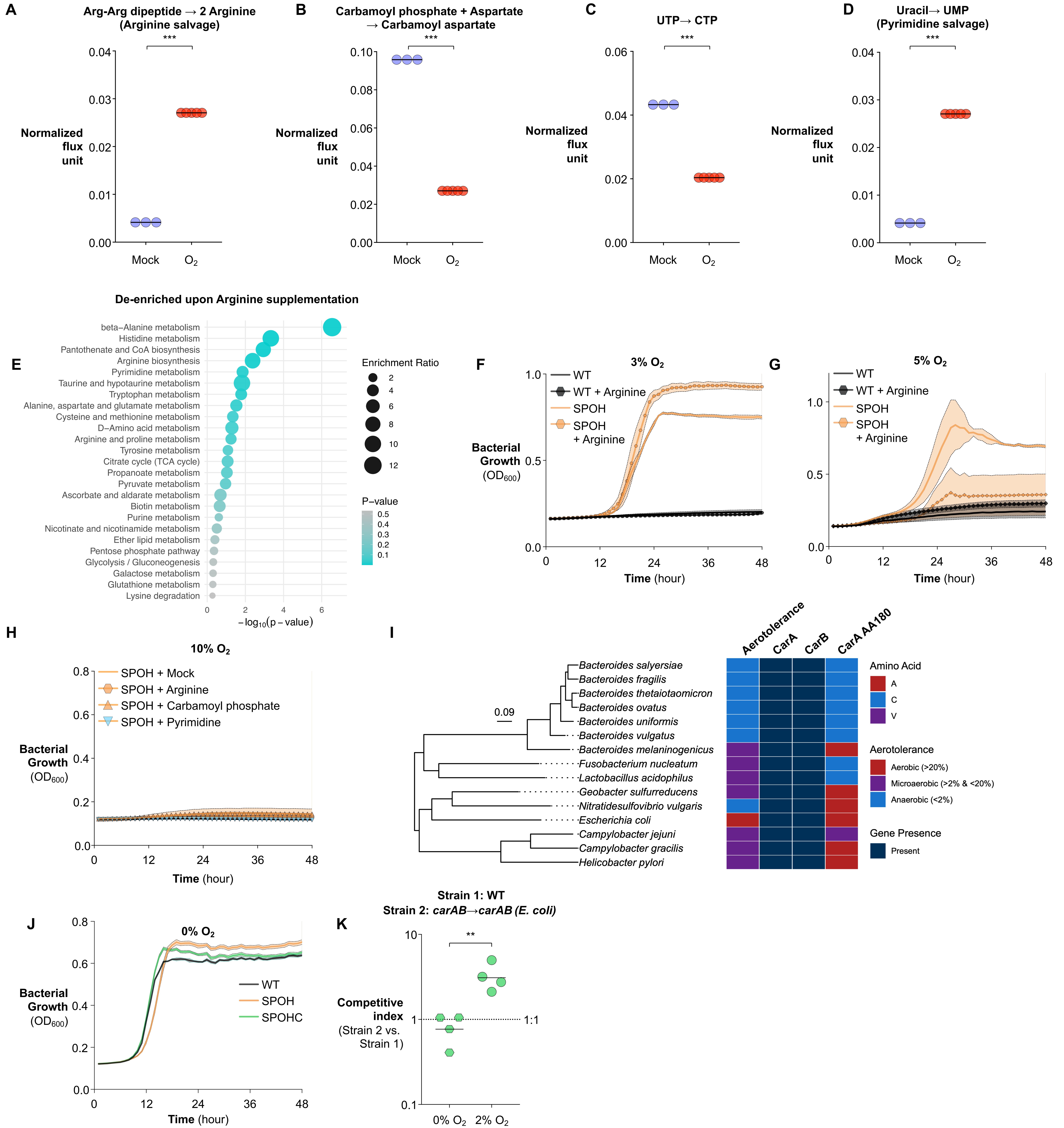

**Supplementary Fig. S7. Oxygen impairs carbamoyl phosphate biosynthesis through cysteine modification in *B. theta* (related to Fig 5).** (**A-D**) Flux balance modeling of CarAB-limited cultures under oxygen exposure predicts reduced de novo pyrimidine biosynthesis and increased salvage pathway activity. Shown are normalized modeled fluxes for (**A**) Arg-Arg dipeptide hydrolysis to arginine, (**B**) carbamoyl phosphate and aspartate conversion to carbamoyl aspartate, (**C**) UTP conversion to CTP, and (**D**) uracil conversion to UMP. (**E**) Anaerobically growing *B. theta* SPOH cultures supplemented with vehicle control or arginine were exposed to 5% $O_2$ for 30 min, and extracellular metabolites were quantified by untargeted UHPLC-MS/MS. Pathway enrichment analysis of metabolites depleted following arginine supplementation in SPOH cultures exposed to 5% $O_2$. (**F,G**) Growth of the indicated *B. theta* strains supplemented with vehicle control or arginine under (**F**) 3% $O_2$ or (**G**) 5% $O_2$ monitored by $OD_{600}$ over time. (**H**) Growth of SPOH cultures supplemented with vehicle control, arginine, carbamoyl phosphate, or pyrimidines under 10% $O_2$, monitored by $OD_{600}$ over time. (**I**) Midpoint-rooted species tree generated using OrthoFinder and annotated with species aerotolerance, *carA* and *carB* carriage, and amino acid identity at the focal residue. Scale bar indicates substitutions per site. (**J**) Growth of WT, SPOH, and SPOHC strains under anaerobic conditions, monitored by $OD_{600}$ over time. (**K**) Pairwise competition between WT and *carAB*→*carAB* (*E. coli*) strains cultured under the indicated oxygen concentrations for 16 h. Competitive indices were determined by selective plating. Bars represent geometric means. *, $P < 0.05$; **, $P < 0.01$; ***, $P < 0.001$.

**Supplemental Figure. S8**

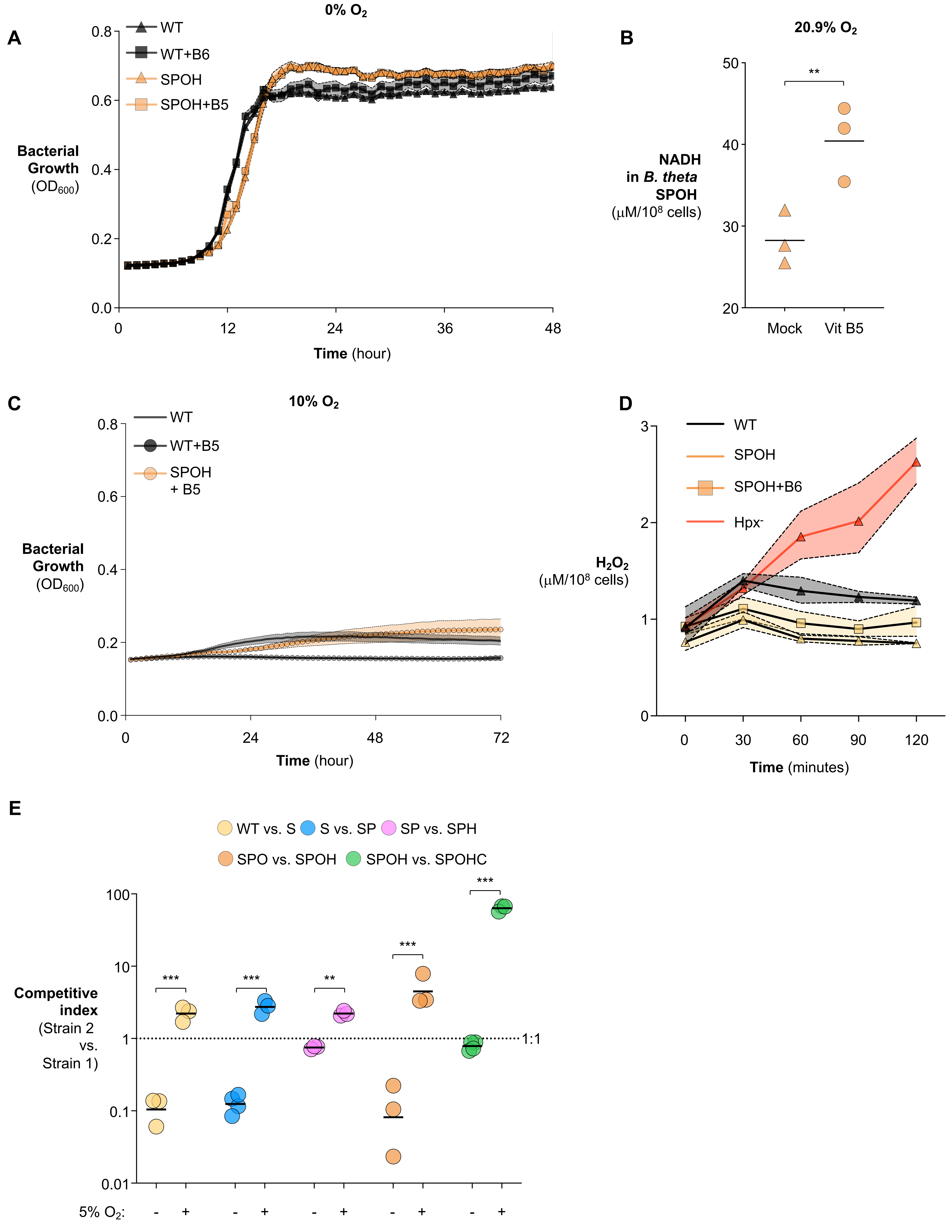

**Supplementary Fig. S8. Oxygen impairs vitamin B5 and B6 biosynthesis in *B. theta* (related to Fig. 6)**. (**A**) Growth of the indicated *B. theta* strains supplemented with vehicle control, vitamin B5 (pantothenate), or vitamin B6 under strictly anaerobic conditions, monitored by $OD_{600}$. (**B**) Actively growing *B. theta* SPOH cultures were supplemented with vitamin B5 or vehicle control and exposed to atmospheric oxygen (20.9% $O_2$); intracellular NADH levels were quantified. (**C**) Growth of *B. theta* cultures supplemented with vitamin B5 in an open atmosphere containing 10% $O_2$, monitored by $OD_{600}$. (**D**) Actively growing *B. theta* SPOH or ROS-scavenging-deficient ($Hpx^-$) strains were supplemented with vitamin B6 or vehicle control and exposed to atmospheric oxygen (20.9% $O_2$); accumulation of $H_2O_2$ in the culture supernatant was quantified. (**E**) Two isogenic *B. theta* strains, a parental strain (strain A) and a derivative strain harboring additional oxygen-resistance modifications (strain B), were mixed at a 1:1 ratio and cultured for 16 h in an open atmosphere containing the indicated oxygen concentrations. Competitive indices were determined by selective plating. Bars represent geometric means. *, $P < 0.05$; **, $P < 0.01$; ***, $P < 0.001$.

Supplemental Figure. S9

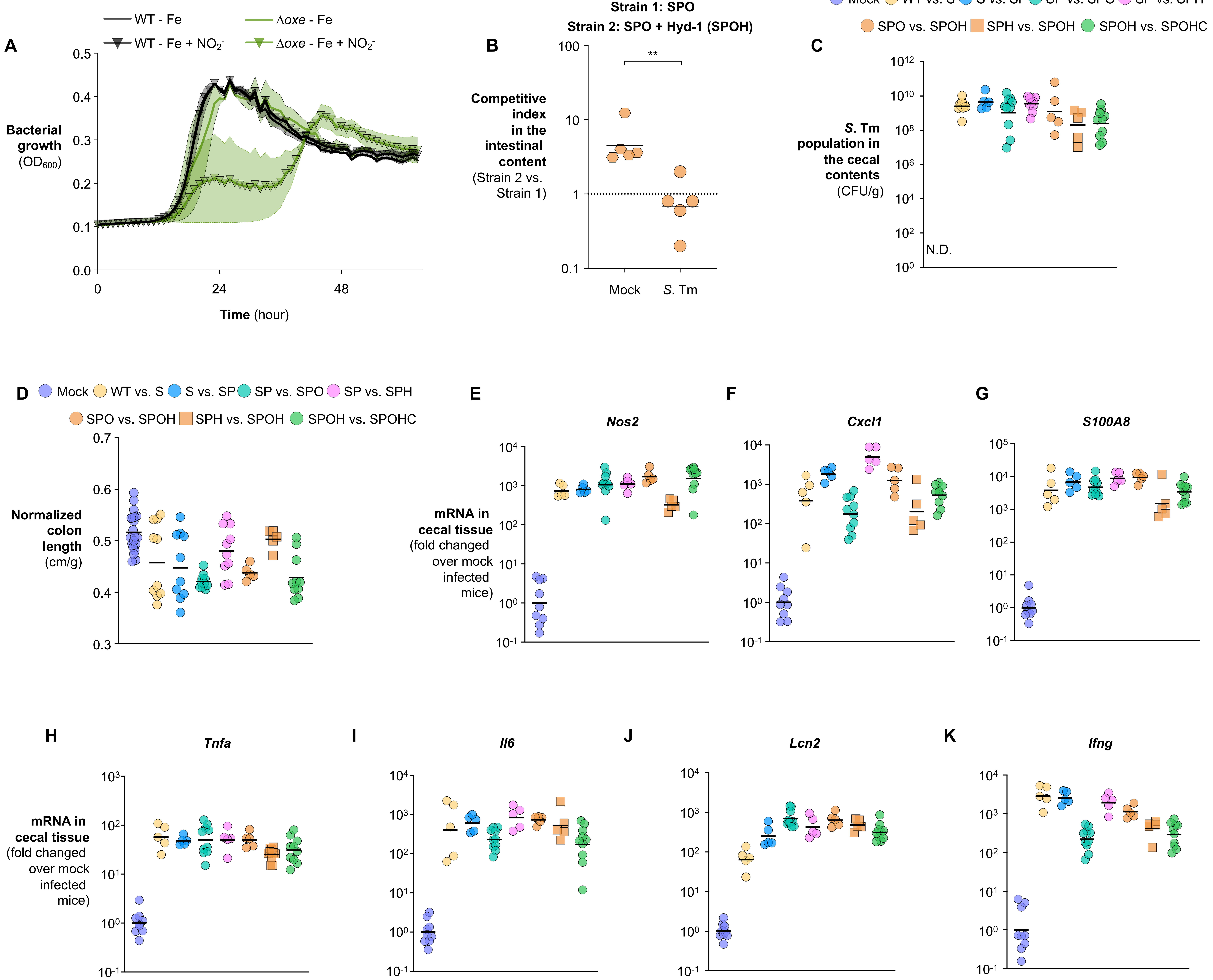

**Supplementary Fig. S9. Stepwise enhancement of oxygen tolerance progressively improves *B. theta* resilience in the oxygenated, inflamed gut** (**related to Fig. 7**) (**A**) The indicated *B. theta* strains were cultured under iron limiting (+BPS) and nitrosative conditions (+ $NO_2^-$). Bacterial growth was monitored by $OD_{600}$. (**B-K**) Antibiotic-pretreated C57BL/6 mice were intragastrically inoculated with an equal mixture of two isogenic *B. theta* strains and subsequently mock treated or infected with *Salmonella enterica* serovar Typhimurium (*S*. Tm) SL1344. Four days post infection, cecal contents and tissues were collected for analysis. (**B**) Competitive indices between SPO and SPOH strains recovered from cecal contents. (**C**) *S*. Tm abundance in cecal contents quantified by selective plating. (**D**) Normalized colon length as a gross indicator of intestinal inflammation. (**E-K**) Relative mRNA abundance of the indicated inflammatory markers in cecal tissue quantified by RT-qPCR: (**E**) *Nos2*, (**F**) *Cxcl1*, (**G**) *S100A8*, (**H**) *Tnfa*, (**I**) *Il6*, (**J**) *Lcn2*, and (**K**) *Ifng*. Bars represent geometric means. *, $P < 0.05$; **, $P < 0.01$; ***, $P < 0.001$.

**Supplemental Figure. S10**

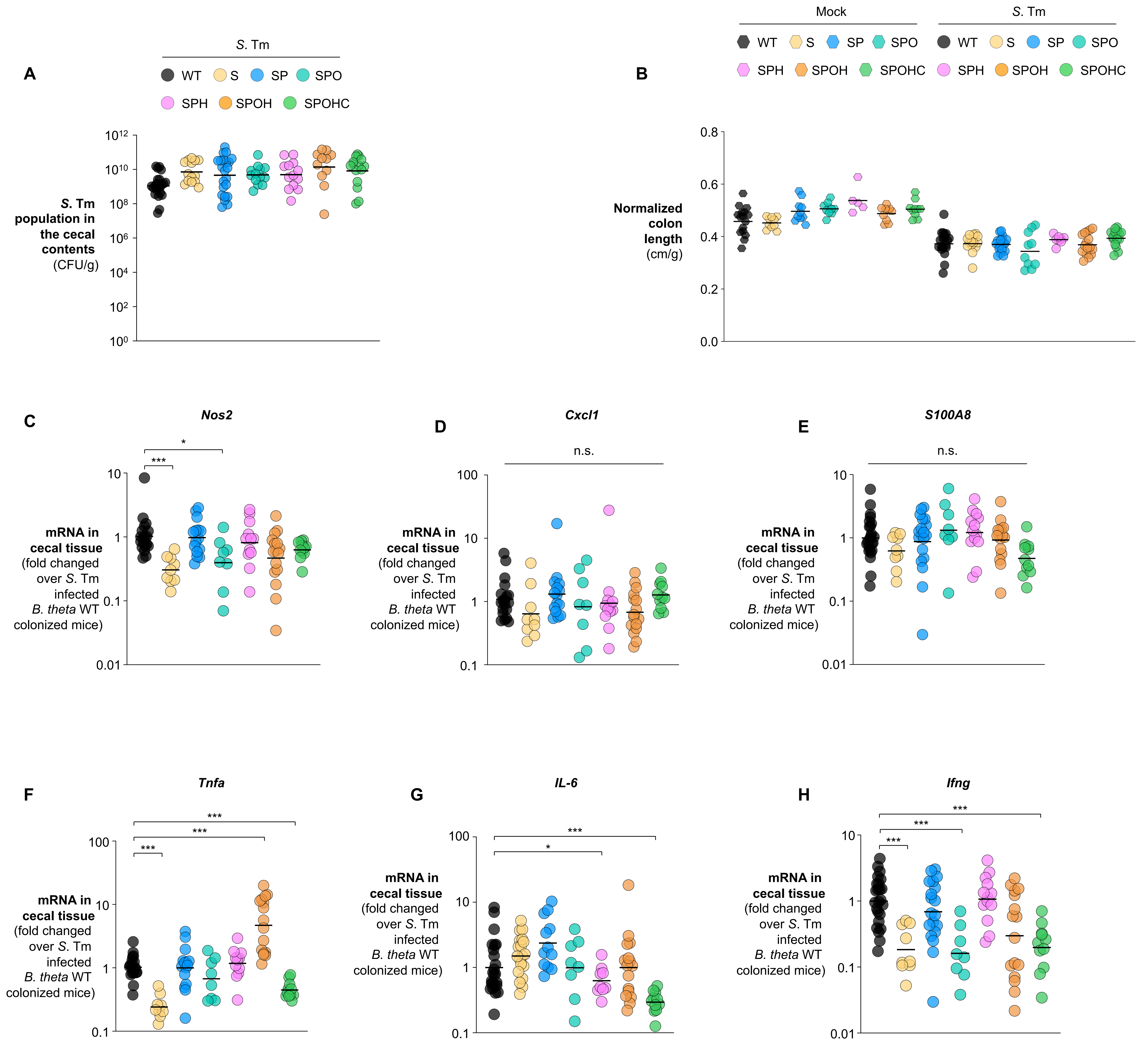

**Supplementary Fig. S10. Stepwise enhancement of oxygen tolerance progressively improves *B. theta* resilience in the oxygenated, inflamed gut (single colonization, related to Fig. 7)** (**A-H**) Antibiotic-pretreated C57BL/6 mice were intragastrically inoculated with individual *B. theta* strains carrying the indicated oxygen-resistance modifications and subsequently mock treated or infected with *Salmonella* enterica serovar Typhimurium (*S*. Tm) SL1344. Four days post infection, cecal contents and tissues were collected for analysis. (**A**) S. Tm abundance in cecal contents quantified by selective plating. (**B**) Normalized colon length as an indicator of intestinal inflammation. (**C-H**) Relative mRNA abundance of inflammatory markers in cecal tissue quantified by RT-qPCR: (**C**) *Nos2*, (**D**) *Cxcl1*, (**E**) *S100A8*, (**F**) *Tnfa*, (**G**) *Il6,* and (**H**) *Ifng*. Bars represent geometric means. *, $P < 0.05$; **, $P < 0.01$; ***, $P < 0.001$.
