## Supplementary material for "Rational engineering of facultative anaerobiosis enables commensal survival in the oxygenated gut": Key Resource Table

**KEY RESOURCES TABLE**

| REAGENT or RESOURCE | SOURCE | IDENTIFIER |
| --- | --- | --- |
| **Bacterial Strains** | | |
| *Bacteroides thetaiotaomicron* VPI-5482 Δ*tdk* (Gen^R^) | [1] | VPI-5482 |
| *S*. Tm wild-type strain (Strep^R^) | [2] | SL1344 |
| *E. coli* S17-1 *λpir*; *zxx::*RP4 2-(Tet^r^::Mu) (Kan^r^::Tn7) *λpir* | [3] | S17-1 *λpir* |
| S17-1 *λpir* + pKI3 | This Study | AR1 |
| S17-1 *λpir* + pKI4 | This Study | AR2 |
| S17-1 *λpir* + pKI3::P_*pfl*::E3 (PDH) | This Study | AR3 |
| S17-1 *λpir* + pKI4::P_*pfl*::*aceEF* (PDH) | This Study | AR4 |
| *B. theta* VPI-5482 *∆tdk*, pKI4::P_*pfl*::*aceEF* (PDH), pKI3::P_*pfl*::E3 (PDH) + BT_2414→BT_0517 | This Study | AR5 |
| *B. theta* VPI-5482 *∆tdk*, pKI4::P_*pfl*::*aceEF* (PDH), pKI3::P_*pfl*::E3 (PDH), BT_2414→BT_0517, pKI7::P_*pfl*::*hydABCDEF* | This Study | AR6 |
| *B. theta* VPI-5482 *∆tdk*, ∆*oxe* | This Study | AR7 |
| *S17-1 λpir +* pExchange*::* ∆*oxe* | This Study | AR8 |
| *B. theta* VPI-5482 *∆tdk*, Δ*katE*, Δ*ahpC1*, Δ*rbr1*, Δ*rbr2* | This Study | AR10 |
| *B. theta* VPI-5482 *∆tdk*, pKI4::P_*pfl*::*aceEF* (PDH), pKI3::P_*pfl*::E3 (PDH) + BT_2414→BT_0517, ∆*oxe* | This Study | AR14 |
| *B. theta* VPI-5482 *∆tdk*, pKI4::P_*pfl*::*aceEF* (PDH), pKI3::P_*pfl*::E3 (PDH), BT_2414→BT_0517, pKI7::P_*pfl*::*hydABCDEF*, ∆*oxe* | This Study | AR18 |
| pNBU2-*bla-ermG* | This Study | AR24 |
| pNBU2-*bla*-*supCfxA* | This Study | AR29 |
| pExchange::ΔBT_0556-BT_0557→*carAB* (*E. coli*) | This Study | AR37 |
| *B. theta* VPI-5482 *∆tdk*, pKI4::P_*pfl*::*aceEF* (PDH), pKI3::P_*pfl*::E3 (PDH), BT_2414→BT_0517, pKI7::P_*pfl*::*hydABCDEF*, ∆*oxe*, ΔBT_0556-BT_0557→*carAB* (*E. coli*) | This Study | AR80 |
| *B. theta* VPI-5482 *∆tdk*, Δ*katE*, Δ*ahpC1*, Δ*rbr1*, Δ*rbr2*, ∆*roo1* | This Study | AR88 |
| *B. theta* VPI-5482 *∆tdk*, Δ*katE*, Δ*ahpC1* | This Study | AR89 |
| pExchange::*B. theta carAB*→*carAB*(C180S) | This Study | AR91 |
| pExchange::*B. theta carAB*→*carAB*(C180D) | This Study | AR92 |
| *B. theta* VPI-5482 *∆tdk*, BT_2414→BT_0517 (pNBU2-ermG) | This Study | AR98 |
| *B. theta* VPI-5482 *∆tdk*, BT_2414→BT_0517 (pNBU2-cfxA) | This Study | AR99 |
| *B. theta* VPI-5482 *∆tdk*, pKI4::P_*pfl*::*aceEF* (PDH), pKI3::P_*pfl*::E3 (PDH) + BT_2414→BT_0517 (pNBU2-ermG) | This Study | AR100 |
| *B. theta* VPI-5482 *∆tdk*, pKI4::P_*pfl*::*aceEF* (PDH), pKI3::P_*pfl*::E3 (PDH) + BT_2414→BT_0517 (pNBU2-cfxA) | This Study | AR101 |
| *B. theta* VPI-5482 *∆tdk*, pKI4::P_*pfl*::*aceEF* (PDH), pKI3::P_*pfl*::E3 (PDH), BT_2414→BT_0517, pKI7::P_*pfl*::*hydABCDEF* (pNBU2-ermG) | This Study | AR102 |
| *B. theta* VPI-5482 *∆tdk*, pKI4::P_*pfl*::*aceEF* (PDH), pKI3::P_*pfl*::E3 (PDH), BT_2414→BT_0517, pKI7::P_*pfl*::*hydABCDEF* (pNBU2-cfxA) | This Study | AR103 |
| *B. theta* VPI-5482 *∆tdk*, pKI4::P_*pfl*::*aceEF* (PDH), pKI3::P_*pfl*::E3 (PDH), BT_2414→BT_0517, pKI7::P_*pfl*::*hydABCDEF*, ∆*oxe* (pNBU2-ermG) | This Study | AR104 |
| *B. theta* VPI-5482 *∆tdk*, pKI4::P_*pfl*::*aceEF* (PDH), pKI3::P_*pfl*::E3 (PDH), BT_2414→BT_0517, pKI7::P_*pfl*::*hydABCDEF*, ∆*oxe* (pNBU2-cfxA) | This Study | AR105 |
| *B. theta* VPI-5482 *∆tdk*, *catP* | This Study | AR106 |
| *B. theta* VPI-5482 *∆tdk*, pKI4::P_*pfl*::*aceEF* (PDH), pKI3::P_*pfl*::E3 (PDH), BT_2414→BT_0517, pKI7::P_*pfl*::*hydABCDEF*, ∆*oxe*, ΔBT_0556-BT_0557→*carAB* (*E. coli*) (pNBU2-ermG) | This Study | AR107 |
| *B. theta* VPI-5482 *∆tdk*, pKI4::P_*pfl*::*aceEF* (PDH), pKI3::P_*pfl*::E3 (PDH), BT_2414→BT_0517, pKI7::P_*pfl*::*hydABCDEF*, ∆*oxe*, ΔBT_0556-BT_0557→*carAB* (*E. coli*) (pNBU2-cfxA) | This Study | AR108 |
| *B. theta* VPI-5482 *∆tdk*, pKI4::P_*pfl*::*aceEF* (PDH), pKI3::P_*pfl*::E3 (PDH), BT_2414→BT_0517, ∆*oxe*, (pNBU2-ermG, pNBU2-cfxA) | This Study | AR109 |
| *B. theta* VPI-5482 *∆tdk*, pKI4::P_*pfl*::*aceEF* (PDH), pKI3::P_*pfl*::E3 (PDH), BT_2414→BT_0517, ∆*oxe*, (pNBU2-ermG) | This Study | AR110 |
| *B. theta* VPI-5482 *∆tdk*, *carAB*→*carAB*(C180S) | This Study | AR111 |
| *B. theta* VPI-5482 *∆tdk*, *carAB*→*carAB*(C180D) | This Study | AR112 |
| *B. theta* VPI-5482 *∆tdk*, ΔBT_0556-BT_0557→*carAB* (*E. coli*) | This Study | AR114 |
| *B. theta* VPI-5482 *∆tdk*, ∆*oxe*, Δ*rbr1*, Δ*rbr2* (pNBU2-cfxA) | This Study | AR121 |
| *B. theta* VPI-5482 *∆tdk*, Δ*rbr1*, Δ*rbr2* (pNBU2-cfxA) | This Study | AR122 |
| *B. theta* VPI-5482 *∆tdk*, Δ*katE*, Δ*ahpC1*, Δ*rbr1*, Δ*rbr2*, ∆*oxe* (pNBU2-cfxA) | This Study | AR123 |
| *B. theta* VPI-5482 *∆tdk*, Δ*katE*, Δ*ahpC1* (pNBU2-cfxA) | This Study | AR124 |
| *B. theta* VPI-5482 *∆tdk*, Δ*katE*, Δ*ahpC1*, Δ*rbr1*, Δ*rbr2*, ∆*oxe*, pNBU2::*oxe*(C222Y) | This Study | AR125 |
| *B. theta* VPI-5482 *∆tdk*, Δ*katE*, Δ*ahpC1*, Δ*rbr1*, Δ*rbr2*, ∆*oxe*, pNBU2::*oxe* | This Study | AR126 |
| *B. theta* VPI-5482 *∆tdk*, pKI4::P_*pfl*::*aceEF* (PDH), pKI3::P_*pfl*::E3 (PDH), BT_2414→BT_0517, pKI7::P_*pfl*::*hydABCDEF*(HydF R509A) (pNBU2-ermG) | This Study | AR131 |
| *B. theta* VPI-5482 *∆tdk*, pKI4::P_*pfl*::*aceEF* (PDH), pKI3::P_*pfl*::E3 (PDH), BT_2414→BT_0517, ∆*oxe*, pNBU2::*oxe* | This Study | AR132 |
| *B. theta* VPI-5482 *∆tdk*, pKI4::P_*pfl*::*aceEF* (PDH), pKI3::P_*pfl*::E3 (PDH), BT_2414→BT_0517, ∆*oxe*, pNBU2::*oxe*(C222Y) | This Study | AR136 |
| *B. theta* VPI-5482 *∆tdk*, ∆*rbx* (pNBU2-tetQ) | This Study | AR138 |
| *B. theta* VPI-5482 *∆tdk*, ∆rbx (pNBU2-ermG) | This Study | AR140 |
| *B. theta* VPI-5482 *∆tdk*, ∆oxe ∆rbx ∆rbr1 ∆rbr2 (pNBU2-ermG) | This Study | AR146 |
| Evolved *B. theta* VPI-5482 *∆tdk*, pKI4::P_*pfl*::*aceEF* (PDH), pKI3::P_*pfl*::E3 (PDH), BT_2414→BT_0517 | This Study | MS13 |
| *B. theta* VPI-5482 *∆tdk*, ∆*oxe* (pNBU2-ermG) | This Study | MS30 |
| *E. coli* S17-1 *λpir* (pNBU2*-bla-ermG*::*oxe*(C222Y)) | This Study | MS32 |
| *B. theta* VPI-5482 *∆tdk*, ∆*oxe* (pNBU2*-bla-ermG*::*oxe*(C222Y)) | This Study | MS34 |
| *E. coli* S17-1 *λpir* + pNBU2*-bla-ermG*::*oxe* | This Study | MS46 |
| *B. theta* VPI-5482 *∆tdk*, ∆*oxe* + pNBU2*-bla-ermG*::*oxe* | This Study | MS49 |
| *E. coli* S17-1 *λpir* + pKI7 | This Study | ML 43 |
| pKI7:: P_*pfl*::*hydABCDEF* | This Study | ML218 |
| *B. theta* VPI-5482 *∆tdk* pKI7::P_*pfl*::*hydABCDEF* | This Study | ML 83 |
| *B. theta* VPI-5482 *∆tdk*, pKI4::P_*pfl*::*aceEF* (PDH), pKI3::P_*pfl*::E3 (PDH), BT_2414→BT_0517, pKI7::P_*pfl*::*hydABCDEF* (pNBU2-tetQ) | This Study | ML 111 |
| *E. coli* S17-1 *λpir* + pExchange::∆*rbr2* | This Study | ML 176 |
| *E. coli* S17-1 *λpir* + pExchange::∆*rbr1* | This Study | ML 179 |
| *B. theta ∆tdk* BT_2414→BT_0517 | This Study | WZ911 |
| *B. theta* VPI-5482 *∆tdk* (pNBU2-tetQ) | This Study | WZ1334 |
| **Chemicals** |  |  |
| Brain Heart Infusion Broth | Becton, Dickinson and Company | 237200 |
| Agar | Becton, Dickinson and Company | 214010 |
| Luria-Bertani (LB) Broth | Becton, Dickinson and Company | 244620 |
| LB Agar, Miller (Luria-Bertani) | Becton, Dickinson and Company | 244510 |
| Menadione (Vitamin K3) | Sigma-Aldrich | M5625-25G |
| Hemin, from Porcine, >96% (HPLC) | Sigma-Aldrich | 51280-1G |
| Erythromycin | Acros Organics | 227330250 |
| Tetracycline | Alfa Aesar | B21408-22 |
| Chloramphenicol | Fisher Scientific | BP904100 |
| Cefoxitin sodium salt | Fisher Scientific | VC00021 |
| Carbenicillin disodium salt | VWR | J358 |
| 5-Fluoro-2′-deoxyuridine (FUdR) | Chem-Impex | 867 |
| Gibson Assembly Master Mix | New England Biolabs | E2611L |
| Adenosine 5′-triphosphate | Sigma-Aldrich | A2383 |
| Methanol, anhydrous, 99.8% | Fisher Scientific | 322415 |
| Tris(hydroxymethyl)aminomethane (TRIS, Trometamol) | Alfa Aesar | J65594 |
| Pyruvate kinase from rabbit muscle | Sigma-Aldrich | P1506 |
| Phosphoenolpyruvate (PEP) | Sigma-Aldrich | P0564 |
| Dodecyltrimethylammonium bromide (DTAB) | Sigma-Aldrich | D8638 |
| PBS | Sigma-Aldrich | P38135 |
| Chloroform, anhydrous, >99% | Sigma-Aldrich | 288306-1L |
| Chloroform | Sigma-Aldrich | C2432 |
| β-Nicotinamide adenine dinucleotide | Sigma-Aldrich | N8285 |
| Methyl viologen | Acros Organics | 227320050 |
| Nickel Chloride Hexahydrate | MP Biomedicals | 219471380 |
| Sodium bicarbonate | Sigma-Aldrich | S6014 |
| L-glutamine | Sigma-Aldrich | G3126 |
| DTT [DL-Dithiothreitol] | RPI | D11000 |
| Hydroxylamine hydrochloride | Sigma-Aldrich | 159417 |
| Sodium carbonate | Sigma-Aldrich | 223530 |
| 2,3-Butanedione monoxime | Sigma-Aldrich | B0753 |
| Antipyrine | Sigma-Aldrich | A5882 |
| Carbamyl phosphate disodium salt | Sigma-Aldrich | C4135 |
| Acetonitrile | EMD Millipore | AX0156 |
| Ammonium bicarbonate | Sigma-Aldrich | A6141 |
| Formic acid | Fisher Scientific | A117 |
| Ampicillin | Cayman Chemical | 69-52-3 |
| Metronidazole | Sigma-Aldrich | M3761 |
| Vancomycin | Chem-Impex | 00315 |
| Neomycin | Sigma | 974104 |
| Formalin | Fisher Scientific | SF100 |
| Tri Reagent® | Sigma-Aldrich | T9424 |
| TURBO DNA-free Kit | Ambion | AM1907 |
| SuperScript VILO cDNA Synthesis Kit | Invitrogen | 11754050 |
| PowerUp SYBR Green Master Mix | Life Technologies | 4309155 |
| Hypoxyprobe Omni Kit (200 mg pimonidazole HCl plus 1 unit of 2627 rabbit antisera) | Hypoxyprobe | HP3-200Kit |
| Hypoxyprobe-Green Kit (100 mg pimonidazole HCl plus 1 unit of 4.3.11.3 mouse FITC-MAb) | Hypoxyprobe | HP6-100Kit |
| Proteinase K, Molecular Biology Grade, 800 U/mL | New England Biolabs | P8107S |
| M.O.M. (Mouse on Mouse) Immunodetection Kit, Basic | Vector Laboratories | BMK-2202 |
| a-pimonidazole rabbit antisera | Hypoxyprobe | Pab2627 |
| Alexa Fluor 488 goat anti-rabbit secondary antibody | Cell Signaling Technology | 4412S |
| Hoechst stain | TargetMol | T21627 |
| Succinate-d4 | Sigma-Aldrich | 293075 |
| Pyruvate-13C3 | Sigma-Aldrich | 490733 |
| Lactate-13C3 | Sigma-Aldrich | MDS0880209 |
| Sodium phosphate dibasic (Na2HPO4) | Fisher Scientific | 13437 |
| Sodium phosphate monobasic (NaH2PO4) | Sigma-Aldrich | S9638 |
| N-(3-Dimethylaminopropyl)-N'-ethylcarbodiimide hydrochloride | Sigma-Aldrich | 03449 |
| Dansyl hydrazine | Sigma-Aldrich | 30434 |
| Trifluoroacetic acid (TFA) | Sigma-Aldrich | 80457 |
| Ammonium formate | Sigma-Aldrich | 70221 |
| L-arginine | Sigma-Aldrich | A5006 |
| Pyridoxal-5-Phosphate Monohydrate | Sigma-Aldrich | P20750 |
| **Critical Commercial Assays** |  |  |
| Gibson Assembly Cloning Kit | NEB | Cat# E2611 |
| Q5 Hot Start 2x Master Mix | NEB | Cat# M0494L |
| NAD/NADH-Glo Assay | Promega | Cat# G9072 |
| BacTiter-Glo Microbial Cell Viability Assay | Promega | Cat# G8230 |
| Pierce™ BCA® Protein Assay Kit | Thermo Scientific | Cat# 23225 |
| **Deposited Data** |  |  |
| Chemoproteomic assessment of cysteine reactivity | PRIDE[4]/ ProteomeXchange[5] | PXD079134 (token: rzvuz8rH7YoT) |
| Bulk RNAseq | European Nucleotide Archive | PRJEB112738 (secondary accession ERP193206) |
| Untargeted Metabolomics data | NIH Common Fund’s National Metabolomics Data Repository (NMDR)/ the Metabolomics Workbench | Study IDs ST003919 and ST004857, Project DOIs: <https://doi.org/10.21228/M8PR93>, <http://dx.doi.org/10.21228/M8QC4Q>. |
| **Experimental Models: Organisms/Strains** |  |  |
| SPF C57BL/6 mice (wild-type) | The Jackson Laboratory | Cat# 000664 |
| **Oligonucleotides** |  |  |
| pExchange_AR_fwd: cactagttctagagcggccg | This Study | N/A |
| pExchange_AR_rev: ccccgggctgcaggaatt | This Study | N/A |
| BT_2414 1 fwd: cgaattcctgcagcccggggATTTCAGTAATTTCCTCCC | This Study | N/A |
| BT_2414 2 rev: caattttattcatTACTTGGTATGCTTCTTTG | This Study | N/A |
| BT_2414→0517 fwd: ataccaagtaATGAATAAAATTGGAGTATTTTATGG | This Study | N/A |
| BT_2414→0517 rev: gtcgattgtaTTAGCTGATTTCTTGTTTTACTTG | This Study | N/A |
| BT_2414 2 fwd: aatcagctaaTACAATCGACAAAAAAGTATAGG | This Study | N/A |
| BT_2414 2 rev: cggccgctctagaactagtgTGCTTACGTAGGTACACAAG | This Study | N/A |
| BT_2414 ver 1: CAGTGGATACTTGGGAACTC | This Study | N/A |
| BT_2414 ver 2: GTTTCATTGCTGGTCCGGAA | This Study | N/A |
| PDH pKI3 US fwd: cacaactgtcGGATCCGCCCTCTTTTGC | This Study | N/A |
| PDH pKI3 US rev: cggccgctctagaactagtgTAATATTCAGGTAAGAGAAATAAAAGAGATCCTG | This Study | N/A |
| PDH pKI3 DS fwd: cgaattcctgcagcccggggGTATGTATATGCTTTTCCAGG | This Study | N/A |
| PDH pKI3 DS rev: gggcggatccGACAGTTGTGAAACTCTC | This Study | N/A |
| PDH pKI3 Open fwd: ggatccgccctcttttgc |  |  |
| PDH pKI3 Open rev: gacagttgtgaaactctcgc |  |  |
| PDH pKI3 proPFL fwd: gcgagagtttcacaactgtctaaaatgataaaattagaggttgtag | This Study | N/A |
| PDH pKI3 proPFL rev: gaagaagaagaaaacttccttatcagaagtg | This Study | N/A |
| PDH pKI3 E3 fwd: aggaagttttcttcttcttcgctttcgg | This Study | N/A |
| PDH pKI3 E3 rev: acgcaaaagagggcggatccatgagtactgaaatcaaaactc | This Study | N/A |
| PDH pKI4 US fwd: tctggacaagaaataaaatgCAAAACTTCCTTATCAGAAGTG | This Study | N/A |
| PDH pKI4 US rev: gttctgacatTAAAATGATAAAATTAGAGGTTGTAG | This Study | N/A |
| PDH pKI4 DS fwd: tatcattttaATGTCAGAACGTTTCCCAAATG | This Study | N/A |
| PDH pKI4 DS rev: ctgacggcgattcttgaaggCATTCATGAGATTACCAGAAAAAAG | This Study | N/A |
| PDH pKI4 Open fwd: cgaattcctgcagcccggggatatgaaaacaagaaaacaaatcc | This Study | N/A |
| PDH pKI4 Open fwd: cggccgctctagaactagtgGGCGGTACCATCCGTGATG | This Study | N/A |
| PDH pKI4 proPFL fwd: tctggacaagaaataaaatgCAAAACTTCCTTATCAGAAGTG | This Study | N/A |
| PDH pKI4 proPFL rev: gttctgacatTAAAATGATAAAATTAGAGGTTGTAG | This Study | N/A |
| PDH pKI4 *aceEF* fwd: tatcattttaATGTCAGAACGTTTCCCAAATG | This Study | N/A |
| PDH pKI4 *aceEF* rev: ctgacggcgattcttgaaggCATTCATGAGATTACCAGAAAAAAG | This Study | N/A |
| pKI7 US fwd: cgaattcctgcagcccggggCTTTCGTGGTGGAAATCC | This Study | N/A |
| pKI7 US rev: aggcttcgggTATGATTACCGGTAGCCTTTTATTG | This Study | N/A |
| pKI7 DS fwd: ggtaatcataCCCGAAGCCTTTCACTTTTATC | This Study | N/A |
| pKI7 DS rev: cggccgctctagaactagtgGTTCGATTATATACGTGATCATCTG | This Study | N/A |
| pKI7 Open fwd: CCCGAAGCCTTTCACTTTTATC | This Study | N/A |
| pKI7 Open rev: TATGATTACCGGTAGCCTTTTATTG | This Study | N/A |
| pKI7 TWIST proPFL HyaAB rev: aaaggctaccggtaatcataCAAAACTTCCTTATCAGAAGTGC | This Study | N/A |
| pKI7 TWIST proPFL HyaAB fwd: actttgcgccAGTTCGGAGGTTTTCCGC | This Study | N/A |
| pKI7 HyaCDEF fwd: cctccgaactGGCGCAAAGTCTCTCCTC | This Study | N/A |
| pKI7 HyaCDEF rev: taaaagtgaaaggcttcgggTTACGTCGGTGCAGCTTC | This Study | N/A |
| D BT4126 1 fwd: cgaattcctgcagcccggggGGGAGAGACACATGATAC | This Study | N/A |
| D BT4126 1 rev: gtgccaggttTCCCACATAGTGGACATTTC | This Study | N/A |
| D BT4126 2 fwd: ctatgtgggaAACCTGGCACGTGCGATG | This Study | N/A |
| D BT4126 2 rev: cggccgctctagaactagtgCCACCCATGAAAAGGTCGATG | This Study | N/A |
| BT4126 ver fwd: TGGTGGAATATGGCATTGAG | This Study | N/A |
| BT4126 ver rev: AGCTGTCTTCGGAACTTTGT | This Study | N/A |
| pNBU2*-bla-ermG*b_fwd: CCCCGGGCTGCAGGAATT | This Study | N/A |
| pNBU2*-bla-ermG*b_rev: GATCCACTAGTTCTAGAGCGGC | This Study | N/A |
| BT4126 Complement fwd: cgctctagaactagtggatctggacgaatatattgacctg | This Study | N/A |
| BT4126 Complement rev: cgaattcctgcagcccggggagtccggctgaataataag | This Study | N/A |
| BT_0216 KO Ver fwd: GATATTGGGACTTTCTATATTTGCATCAG | This Study | N/A |
| BT_0216 KO Ver rev: GCAGCTTTTCCGTGGAGC | This Study | N/A |
| BT_3182 KO Ver fwd: TCATCAAGAAAAGGGAGC | This Study | N/A |
| BT_3182 KO Ver rev: TAGGAGCTGTTTATAAAACG | This Study | N/A |
| BT_3182 Up_fwd: cgaattcctgcagcccggggAACGGCATCCGGCAATTC | This Study | N/A |
| BT_3182 Up_rev: tcttcacctcGGTTCCTTTAATACTTTTAGTCATAATTCTAATCTATATTTAAATTG | This Study | N/A |
| BT_3182 Down_fwd: taaaggaaccGAGGTGAAGAAAGAAAACTATTAATAATAAG | This Study | N/A |
| BT_3182 Down_rev: cggccgctctagaactagtgAAGAGACAAAGAAGCACTTTC | This Study | N/A |
| BT_0216 Up_fwd: cgaattcctgcagcccggggGAACGGACAAGCTCTGGC | This Study | N/A |
| BT_0216 Up_rev: cgaaataacgTACAGTACATCTAAATTTTTTCATTTTCTTCTCAATTTTAAATTAATTAG | This Study | N/A |
| BT_0216 Down_fwd: atgtactgtaCGTTATTTCGGAGATAAGAAATAATAAAG | This Study | N/A |
| BT_0216 Down_rev: cggccgctctagaactagtgCTGCCTTGTTTAACGCAAC | This Study | N/A |
| BT_CarAB_500_us_fwd: ttcctgcagcccgggggatctcggcattctcgaccgtc | This Study | N/A |
| BT_CarAB_500_us_rev: ttattatttctcttatttttgaatcgttgtatggttgttag | This Study | N/A |
| BT_CarAB_500_ds_fwd: tcaaaaataagagaaataataaattgttgtataataagttgg | This Study | N/A |
| BT_CarAB_500_ds_rev: cggccgctctagaactagtgaggatattgctttcaggaatc | This Study | N/A |
| CarAB_Swap_Open fwd: gaaataataaattgttgtataataa | This Study | N/A |
| CarAB_Swap_Open rev: tcttatttttgaatcgttgtatggt | This Study | N/A |
| Nissle_CarAB_fwd: acaacgattcaaaaataagaatgaccggttatcaagaaatc | This Study | N/A |
| Nissle_CarAB_rev: tatacaacaatttattatttcttatttgatctgcgcgtg | This Study | N/A |
| 180_500US_fwd: gatatcgaattcctgcagcccgggggccacaaccagccggtac | This Study | N/A |
| 180_500US_rev: accttccggcttttctttctggattactttctctacaaaatacgg | This Study | N/A |
| 180_TWIST_fwd: aatccagaaagaaaagccggaaggtattatg | This Study | N/A |
| 180_TWIST_rev: aaagttctccatattacaaactgtaatacagttgtc | This Study | N/A |
| 180_500DS_fwd: tacagtttgtaatatggagaactttgacccac | This Study | N/A |
| 180_500DS_rev: ggtggcggccgctctagaactagtgatcatacggaggcccttc | This Study | N/A |
| *AceEF* qPCR fwd: GATGCTGCCGATTTCTCTCT | This Study | N/A |
| *AceEF* qPCR rev: GTCAGACAGCGTGTTGTTAATG | This Study | N/A |
| E3 qPCR fwd: ACCAAGCGTATCAGCAAGAA | This Study | N/A |
| E3 qPCR rev: TGCCTTCCATCGTCACATAAA | This Study | N/A |
| Oxe qPCR fwd: TTCCAGGGTGATCGGTATCT | This Study | N/A |
| Oxe qPCR rev: CCGCCATCTGTTCCGTATTT | This Study | N/A |
| HydA qPCR fwd: CACCTGCTGTACCGAATCTT | This Study | N/A |
| HydA qPCR rev: ATCGTCGTAATCGAGGGAAATC | This Study | N/A |
| HydB qPCR fwd: CCGCAGGTAAGTTGCAGTAT | This Study | N/A |
| HydB qPCR rev: AGGTTGCAGGTTCCCATTT | This Study | N/A |
| HydC qPCR fwd: GTTGATGGTCACCGGATACTT | This Study | N/A |
| HydC qPCR rev: CCTGATGTAGCCCATATAGAACAG | This Study | N/A |
| HydD qPCR fwd: GAATCGCGCTGTCTCAATTATG | This Study | N/A |
| HydD qPCR rev: TCCTGTGTCATCCGGTACT | This Study | N/A |
| HydE qPCR fwd: CAGACGGCGTGGTGTTATTA | This Study | N/A |
| HydE qPCR rev: TGCCATGTATAGTCGGGAAAC | This Study | N/A |
| HydF qPCR fwd: ACGCCATGTCTGGCATTTA | This Study | N/A |
| HydF qPCR rev: CTTCCGGTATTGGGCAGATT | This Study | N/A |
| **Recombinant DNA** |  |  |
| pKNOCK-*bla*-*ermGb*::*tdk* | [1] | pExchange-*tdk* |
| pExchange-*tdk*::Upstream BT_2414→BT_0517::Downstream BT_2414 | This Study | pExchange-*tdk::* *BT_2414->BT_0517* |
| pExchange-*tdk*::intergenic region of BT_4719 and BT_4720 | This Study | pKI3 |
| pExchange-*tdk*::intergenic region of BT_1200 and BT_1201 | This Study | pKI4 |
| pExchange-*tdk*::Upstream *oxe*::*oxe* | This Study | pExchange-*tdk::* *∆oxe* |
| pNBU2-bla-ermG::*oxe* | This Study | pNBU2*::oxe*^WT^ |
| pNBU2-bla-ermG::*oxe*(C222Y) | This Study | pNBU2*::oxe*^C222Y^ |
| pExchange-*tdk*::intergenic region of BT_3019 and BT_3020 | This Study | pKI7 |
| pExchange-*tdk*::Upstream BT_0556::ECN_CarAB::Downstream BT_0557 | This Study | pExchange-*tdk::* BT_*carAB->carAB* (*E. coli*) |
| pExchange-*tdk*::Upstream BT_0557:: BT_0557[C180S]::Downstream BT_0557 | This Study | pExchange-*tdk::carAB*→*carAB*(C180S) |
| pExchange-*tdk*::Upstream BT_0557:: BT_0557[C180D]::Downstream BT_0557 | This Study | pExchange-*tdk::carAB*→*carAB*(C180D) |
| **Software and Algorithms** |  |  |
| Excel for Mac | Microsoft | v16.109 |
| Prism | Graph Pad | v11.0.0 |
| AlphaFold3 | [6] |  |
| Pymol | The PyMOL Molecular Graphics System, Schrödinger | v3.1.6.1 |
| UCSF Chimera X | [7] | v1.11.1 |
| BioRender | BioRender.com | N/A |
| Protenix Server | [8] | V2 |
| CAVER | [9] | 3.0.3 |
| ConSurf-DB | [10] | N/A |
| MetaboAnalyst 6.0 | [11] | 6.0 |
| Progenesis QI | Waters | v3.0 |
| NIS-Elements software | Nikon | v6.10.01 |
| isoTOP-ABPP | [12] | N/A |
| ReDiMe | [13] | N/A |
| Thermo Proteome Discoverer | Thermo Fisher Scientific | v2.4 |
| SequestHT | Thermo Fisher Scientific | V3.1 |
| Percolator | [14] | Rel-3-09 |
| COBRA Toolbox | [15] | v2.13.3 |
| MATLAB | MathWorks | R2022b |
| Foldseek | [16] | N/A |
| Thermo Xcalibur | Thermo Fisher Scientific | v4.1.31.9 |
| Thermo LCQuan | Thermo Fisher Scientific | v2.7 |
| BBMap | Bushnell B. - sourceforge.net/projects/bbmap/ | Ver 39.88 |
| Bowtie2 | [17] | Ver 2.3.5 |
| featureCounts | [18] | 2.0.5 |
| DESeq2 | [19] | V1.50.2 |
| StanDep | [20] | V1.0 |
| mCADRE | [21] | N/A |
| NCBI Datasets Tool | [22] | v18.19.0 |
| Prokka | [23] | v1.14.5 |
| Orthofinder | [24-27] | v3.1.3.post1.dev1 |
| MAFFT | [28] | v7.526 |
| ClipKIT | [29] | v2.11.4 |
| IQ-TREE | [30] | v3.1.0 |
| ModelFinder | [31] | N/A |
| ggtree | [29] | v4.1.2 |

**Key Resource Table Reference:**

1.	Koropatkin, N.M., et al., *Starch catabolism by a prominent human gut symbiont is directed by the recognition of amylose helices.* Structure, 2008. **16**(7): p. 1105–15.

2.	Hoiseth, S.K. and B.A. Stocker, *Aromatic-dependent Salmonella typhimurium are non-virulent and effective as live vaccines.* Nature, 1981. **291**(5812): p. 238–9.

3.	Simon, R., U. Priefer, and A. Puhler, *A Broad Host Range Mobilization System for In*

*Vivo Genetic Engineering: Transposon Mutagenesis in Gram Negative Bacteria.* Nature Biotechnology, 1983. **1**: p. 784–791.
